# Cross-species phosphoproteomic mapping of mTORC1 signalling reveals RNA splicing regulation during neurodevelopment

**DOI:** 10.64898/2026.11.16.766904

**Authors:** Simeon R. Mihaylov, Marie Girodengo, Sara Guerrisi, Pranetha Baskaran, Laura Kleckner, Helen R. Flynn, J. Mark Skehel, Suzanne Claxton, Adrian Harwood, Matthew White, Deepak P. Srivastava, Sila K. Ultanir, Joseph M. Bateman

**Affiliations:** Kinases and Brain Development Lab, The Francis Crick Institute, 1 Midland Road, London NW1 1AT, UK; Department of Basic and Clinical Neuroscience, King’s College London, 5 Cutcombe Road, London SE5 9RX, UK; Neuroscience & Mental Health Innovation Institute, Cardiff University, Maindy Rd, Cardiff CF24 4HQ, UK; Proteomics Science Technology Platform, The Francis Crick Institute, 1 Midland Road, London NW1 1AT, UK; MRC Centre for Neurodevelopmental Disorders, King’s College London, London, UK; Department of Translational Genomic, Institute of Mental Health and Neuroscience, Maastricht University, Universiteitssingle 50, 6200 MD Maastricht, NL

**Keywords:** proteomics, phosphoproteomics, mTOR, TSC, neuron, splicing, PhosphoTargets

## Abstract

Mechanistic target of rapamycin complex 1 (mTORC1) regulates diverse cellular processes through its kinase activity and is essential for nervous system development. Although mTORC1 hyperactivation causes numerous neurodevelopmental disorders, its downstream targets in the developing nervous system remain poorly defined. Here, by profiling thousands of proteins and tens of thousands of phosphosites during the neurogenic phase in mouse and human models, we systematically define the proteomic and phosphoproteomic landscape downstream of mTORC1 during neurodevelopment. To facilitate community access, we developed PhosphoTargets, an interactive platform comprising protein and phosphoprotein expression data from mTORC1 models and patient tissue. We demonstrate the utility of this resource by identifying the deubiquitylating enzyme USP9X as a novel mTORC1 substrate and showing that mTORC1-dependent phosphorylation of USP9X regulates human neural progenitor cell proliferation. Furthermore, integration with gene expression datasets reveals a new role for mTORC1 in the regulation of mRNA splicing. Together, these data provide a comprehensive resource for investigating mTORC1 signalling mechanisms and reveal novel processes through which mTORC1 regulates neurodevelopment.

## Introduction

mTOR (mechanistic target of rapamycin) is a large, highly conserved serine/threonine kinase that regulates critical cellular processes including cell growth, autophagy, lipid synthesis, glucose metabolism, cytoskeletal dynamics and cell survival (*1, 2*). mTOR was discovered in *Saccharomyces cerevisiae* with the identification of the genes *TOR1* and *TOR2,* mutations in which conferred resistance to rapamycin (*3, 4*). Mammalian mTOR was discovered through analysis of rapamycin-interacting proteins across diverse mammalian cell types and identified as a homolog of TOR1/2 (*5–8*).

mTOR acts as the catalytic core of two protein complexes: mTOR complex 1 (mTORC1) and mTOR complex 2 (mTORC2). mTORC1 is a key regulator of neurogenesis in both invertebrates and mammals (*9, 10*). The neurogenic role of mTOR was first identified through a genetic screen for genes causing defects in mouse forebrain development, which showed that loss of mTOR prevented formation of the telencephalic vesicles (*11*). The mTOR pathway was subsequently shown to play key roles in neurogenesis in *Drosophila* (*12–16*). Since these pioneering studies, mTOR signalling has been shown to regulate many aspects of early brain development in rodent models and humans, including neuronal progenitor genesis, proliferation, differentiation and migration (*10, 17–19*).

Hyperactivation of mTORC1 signalling during neurodevelopment results in a group of rare neurological disorders known as mTORopathies. mTORopathies share two major symptoms: malformations of cortical development and intractable epilepsy (*20, 21*). Tuberous sclerosis complex (TSC) is the most common mTORopathy and causes neuropsychiatric conditions, including infantile spasms, intractable epilepsy, autism and cognitive disability in more than 90% of patients (*22, 23*).

Understanding of the regulation of nervous system development by mTORC1 is limited by the paucity of knowledge of its targets in neurodevelopment. Phosphoproteomic resources in mammalian cell lines exist (*24–26*), but there is no equivalent for the developing nervous system. This limits the ability to interrogate mTOR-related neurodevelopmental disorders (NDDs), design targeted therapies, or connect mTORopathies such as TSC to broader neurodevelopmental disorder networks. To address this, we used quantitative proteomics and phosphoproteomics to identify the targets of mTORC1 in mouse and human models of neurodevelopment. We identify 32 novel targets of mTORC1 in the embryonic murine brain, with functions including chromatin and transcriptional regulation, autophagy and RNA metabolism. In human neural progenitor cells (NPCs) we identify 64 novel targets of mTORC1 and reveal protein networks with functions including mRNA splicing, many of which are associated with NDDs. We validate this phosphoproteomic resource by showing that the novel target USP9X is directly phosphorylated by mTOR and mTORC1-dependent phosphorylation of USP9X regulates the proliferation of human NPCs. Finally, consistent with changes in phosphorylation of splicing factors, we integrate transcriptomic and proteomic data which revealed that mTORC1 activation causes largescale perturbation of mRNA splicing in human NPCs. Overall, our findings demonstrate the potential of this proteomic and phosphoproteomic resource for the discovery of novel mTORC1-dependent cellular and molecular processes in neurodevelopment.

## Results

### Creation of a resource for the interrogation and discovery of novel mTORC1 regulated processes in neurodevelopment

Our strategy to survey the mTORC1 dependent proteome and phosphoproteome in neurodevelopment centred around the use of Tsc1 mutant mouse and TSC1 mutant human cellular models. In mice, we used conditional deletion of *Tsc1* in neural progenitors from embryonic day (E) 10.5 using *Nestin-Cre* and analysed brain tissue at E16.5. As a human model, we generated *TSC1* mutant human induced pluripotent stem cells (hiPSCs) and differentiated them to NPCs. These stages in development were selected as representing high levels of neurogenesis together with diverse progenitor and differentiated cell populations. We used Tsc1 mutant mice or TSC1 mutant NPCs compared to controls, and rapamycin treated Tsc1 mutant mice or rapamycin treated TSC1 mutant NPCs, to capture both mTORC1 activation and inhibition. To maximise coverage, we performed parallel experiments using trypsin and chymotrypsin digests followed by Tandem Mass Tag (TMT) liquid chromatography-tandem mass spectrometry (LC-MS/MS) based proteomics and phosphoproteomics. The data from these 16 LC-MS/MS experiments were made available on PhosphoTargets (https://phosphotargets.com) for easy interrogation and analysis.

### Analysis of the proteome and phosphoproteome during mTORC1 activation and inhibition in the developing mouse brain

We first used western blot analysis of *Nestin-Cre;Tsc1^fl/fl^*(Tsc1 cKO) E16.5 brain tissue and confirmed almost complete loss of TSC1 expression and a strong reduction in TSC2 expression levels, consistent with the requirement for TSC1 to stabilise TSC2 and prevent its ubiquitin-mediated degradation (Figure 1A; Supplementary figure S1A-D) (*27–31*). As expected, phosphorylation of mTORC1-dependent RPS6 S240/244 and 4E-BP T37/46, relative to their total protein levels, were increased in the Tsc1 cKO brain, evidence of mTORC1 hyperactivation (Supplementary figure S1A, B, E, F). mTOR S2448, which is phosphorylated by p70S6 kinase (*32*), was unchanged relative to total mTOR in the Tsc1 cKO brain (Supplementary figure 1A, B, G). Western blot analysis of rapamycin treated Tsc1 cKO embryos (Figure 1B) confirmed almost complete loss of RPS6 S240/244 and 4E-BP T37/46 phosphorylation and a significant decrease in phosphorylation of mTOR S2448 (Supplementary figure S2A-E). Phosphorylation of the mTORC2 target PKCα S657 (*33*) was unchanged in Tsc1 cKO embryonic brain tissue compared to controls or with rapamycin treatment (Supplementary figures S1A, B, H and S2A, B, F).

**Figure 1.**
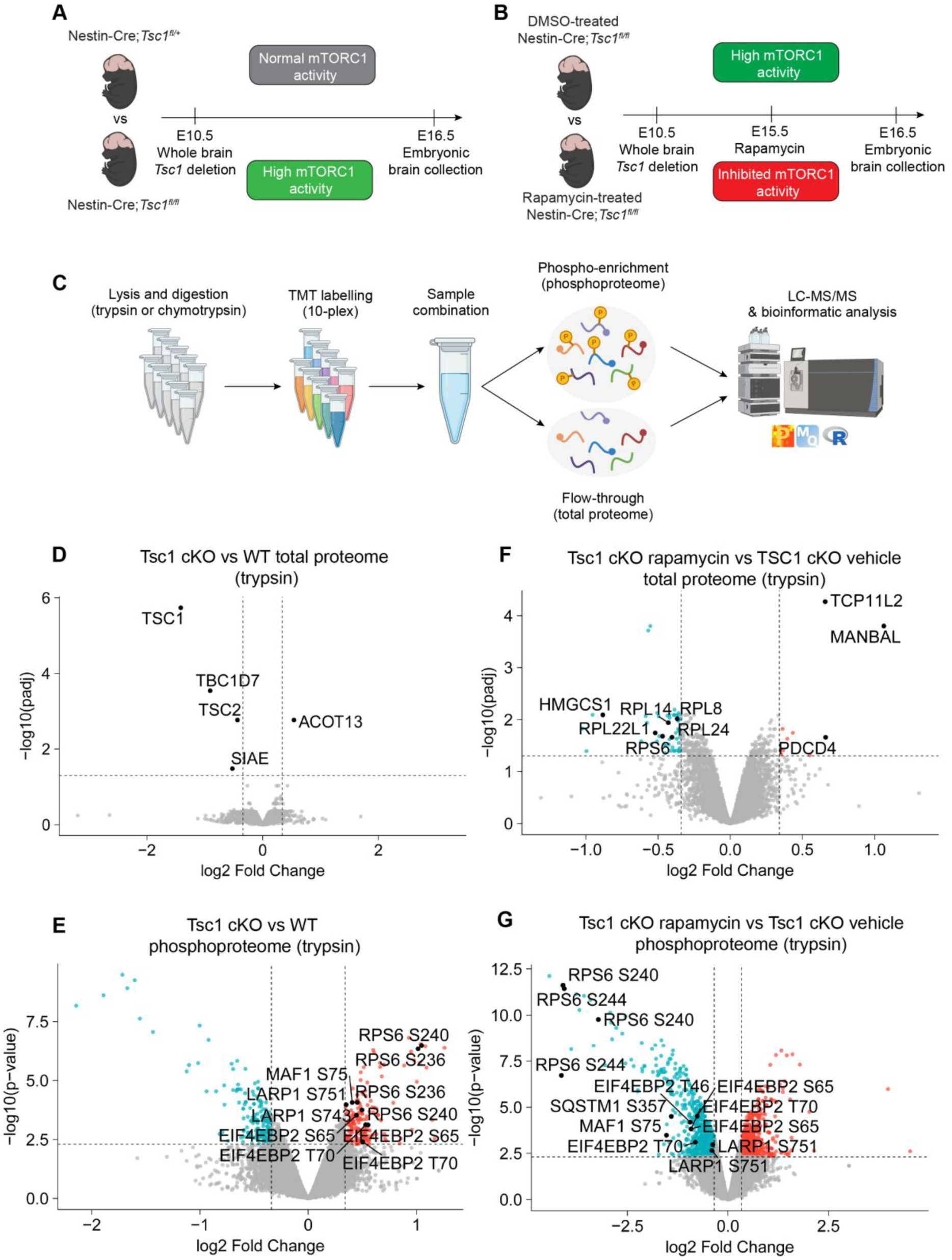
Characterisation of the mouse neurodevelopmental mTORC1 dependent phosphoproteome. (A) Schematic of the Tsc1 cKO mouse model versus control experiment used to collect E16.5 brain tissue for proteomics and phosphoproteomics. (B) Schematic of the rapamycin treated versus vehicle treated Tsc1 cKO mouse model experiment used to collect E16.5 brain tissue for proteomics and phosphoproteomics. (C) Schematic of the TMT labelling and proteomic/phosphoproteomic pipeline. (D, E) Volcano plots of the proteomic (D) and phosphoproteomic (E) data from E16.5 Tsc1 cKO mice versus control brain tissue (trypsin digest). Established mTORC1 targets are labelled in (E). Control n=5 brains, Tsc1 cKO n=4 brains. (F, G) Volcano plots of the proteomic (F) and phosphoproteomic (G) data from E16.5 Tsc1 cKO rapamycin treated mice versus Tsc1 cKO vehicle treated mice brain tissue (trypsin digest). Established mTORC1 targets are labelled in (G). Vehicle treated Tsc1 cKO n=5 brains, rapamycin treated Tsc1 cKO n=5 brains. Proteins or phosphopeptides with significantly increased expression coloured red and significantly decreased expression coloured blue.

Having confirmed mTORC1 perturbation in the Tsc1 cKO mouse model, we next performed quantitative proteomics and phosphoproteomics of whole brain tissue (Figure 1A-C). First we compared E16.5 Tsc1 cKO mice embryos to littermate controls and identified 40,276 phosphopeptides, corresponding to 8,736 proteins with trypsin, and 16,882 phosphopeptides, corresponding to 5,997 proteins with chymotrypsin (Supplementary tables S1-S4). Only 5 proteins had significantly altered expression in the Tsc1 cKO brain (Figure 1D, Supplementary figure S1I, Supplementary tables S1, S3). Consistent with the western blot analysis, TSC1 and TSC2 expression were strongly decreased, along with the third member of the TSC complex TBC1D7 (Figure 1D; Supplementary figure S1I; Supplementary tables S1, S3). In addition, Acyl-coenzyme A thioesterase 13 (ACOT13) expression was significantly increased and Sialate O-acetylesterase (SIAE) expression was significantly decreased (Figure 1D; Supplementary figure S1I; Supplementary tables S1, S3).

Phosphoproteomic data from Tsc1 cKO mice showed a significant increase in phosphorylation of 275 phosphopeptides, corresponding to 155 proteins, including RPS6 and the established mTORC1 substrates EIF4EBP2, MAF1 and LARP1 (*34*) (Figure 1E; Supplementary tables S2, S4; Supplementary figure S1J). Gene ontology (GO) and Reactome analysis of these 155 proteins confirmed strong enrichment for mTORC1 signalling (Supplementary figure S1K, L).

Quantitative proteomics and phosphoproteomics of rapamycin treated compared to vehicle treated Tsc1 cKO embryonic brain tissue identified 33,574 phosphopeptides, corresponding to 8,009 proteins, with trypsin, and 14,283 phosphopeptides, corresponding to 6,123 proteins with chymotrypsin (Supplementary tables S5-S8). Rapamycin treatment of Tsc1 cKO mice significantly changed the expression of 83 proteins including reducing the expression of multiple ribosomal proteins, such as the mTORC1 target RPS6, consistent with the role of mTORC1 as a regulator of protein translation (Figure 1F; Supplementary figure S2G; Supplementary tables S5, S7) (*34*). Rapamycin mediated mTORC1 inhibition in the embryonic brain resulted in reduced phosphorylation of 1010 phosphopeptides, corresponding to 512 proteins, including RPS6 and established mTORC1 substrates EIF4EBP2, MAF1, LARP1, SQSTM1 (P62) (*34*) (Figure 1G; Supplementary figure S2H; Supplementary tables S6, S8). GO analysis of these proteins highlighted regulation of gene expression, metabolic processes and translation (Supplementary figure S2I).

### Novel mTORC1 targets in the developing brain have diverse functions

To identify high confidence mTORC1 targets we next used a pipeline to select phosphosites that were significantly increased in Tsc1 cKO mice versus control *and* significantly decreased by Tsc1 cKO mice rapamycin treatment (Supplementary figure S3). This pipeline resulted in selection of 81 phosphosites corresponding to 44 proteins. Of these 44 proteins, 12 were previously reported as direct substrates of mTORC1 or indirectly regulated by mTORC1 via S6 kinase (Figure 1E, G; Supplementary figures S1J, S2H; Supplementary tables S9 and S10) (*34, 35*). Interestingly, 11 (of 31) phosphosites in known mTORC1 or S6K substrate proteins have not been previously reported (DNAJC2 T48, S49; RPS6 T241, S242, S246; S6K1 S398, T399; SQSTM1 S357; SRPK2 S490, S491, T492) (Figure 1E, G; Supplementary figures S1J, S2H; Supplementary table S9).

The remaining 50 (of 81) phosphosites in 32 (of 44) proteins were not in previously established mTORC1 or S6K substrates. These 32 proteins are thus novel high-confidence mTORC1 targets (potentially as direct substrates or indirectly regulated) in neurodevelopment (Figure 2A, B; Table 1). The total levels of these proteins were unchanged (Supplementary tables S1, S3, S5, S7), indicating that these proteins are regulated by mTORC1 at the level of phosphorylation. The novel mTORC1 target proteins have a range of functions in chromatin/transcriptional regulation, autophagy/protein turnover, cytoskeleton regulation, RNA metabolism, cell division and translation (Figure 2C; Supplementary table S11). Identification of this inventory of high confidence novel mTORC1-dependent phosphosites unlocks the potential to interrogate their role in neurodevelopment.

**Figure 2.**
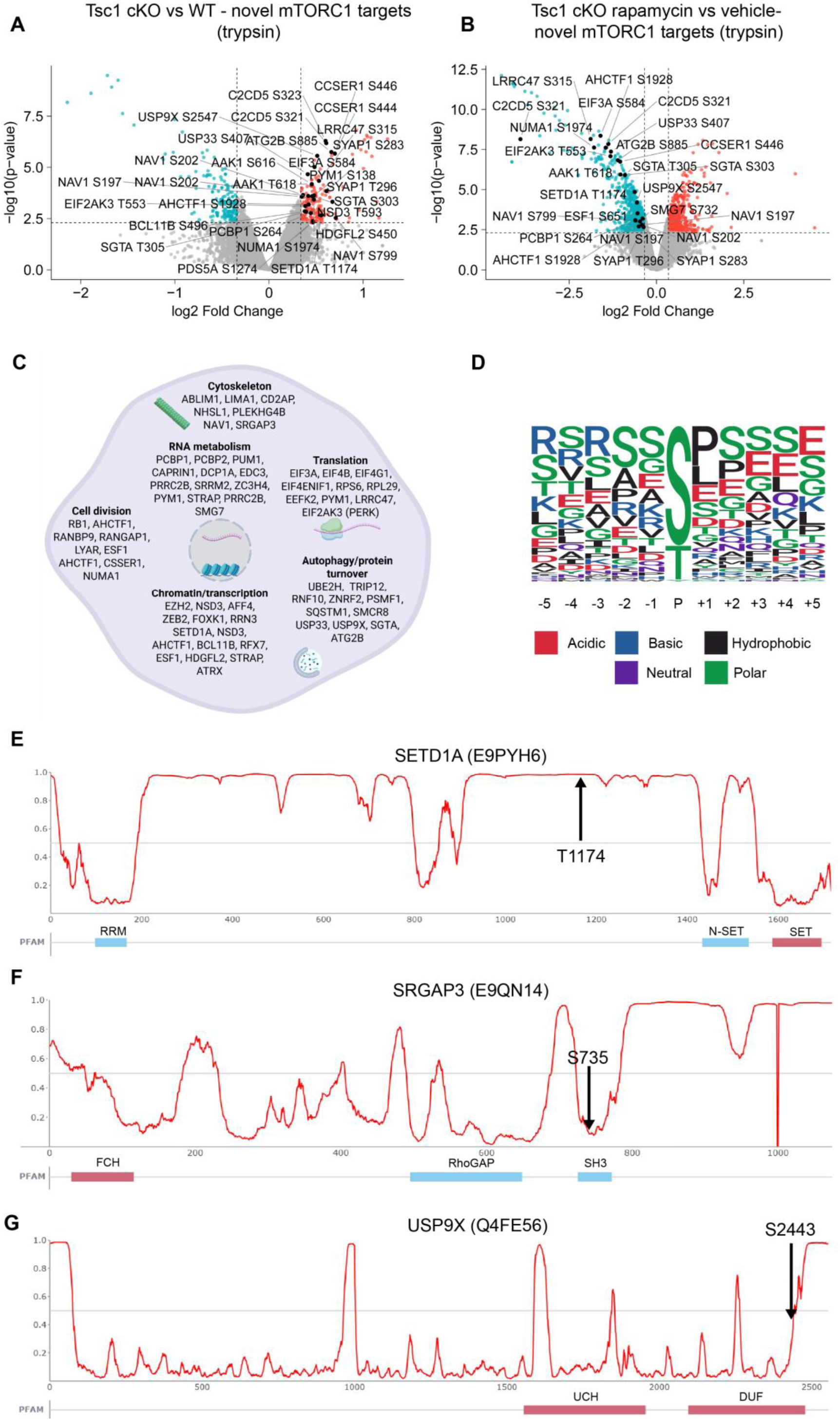
Novel mTORC1 targets in mouse neurodevelopment. (A) Volcano plot of phosphoproteomic data from E16.5 Tsc1 cKO mice versus control brain tissue showing novel mTORC1 targets. Phosphopeptides with significantly increased expression coloured red and significantly decreased expression coloured blue. Control n=5 brains, Tsc1 cKO n=4 brains. (B) Volcano plot of phosphoproteomic data from E16.5 Tsc1 cKO rapamycin treated mice versus Tsc1 cKO vehicle treated mice brain tissue showing novel mTORC1 targets. Phosphopeptides with significantly increased expression coloured red and significantly decreased expression coloured blue. Vehicle treated Tsc1 cKO n=5 brains, rapamycin treated Tsc1 cKO n=5 brains. (C) Diagram illustrating the cellular and molecular functions of novel mTORC1 targets identified in the developing mouse brain and human NPCs. (D) Consensus motif of the mTORC1 target phosphosites identified in the developing mouse brain. (E) Disorder plot of SETD1A showing the position of mTORC1 target phosphosite T1174 in an IDR. (F) Disorder plot of SRGAP3 showing the position of mTORC1 target phoshosite S735 within the SH3 domain. (G) Disorder plot of USP9X showing the position of mTORC1 target phoshosite S2443 within the DUF domain.

**Table 1.**
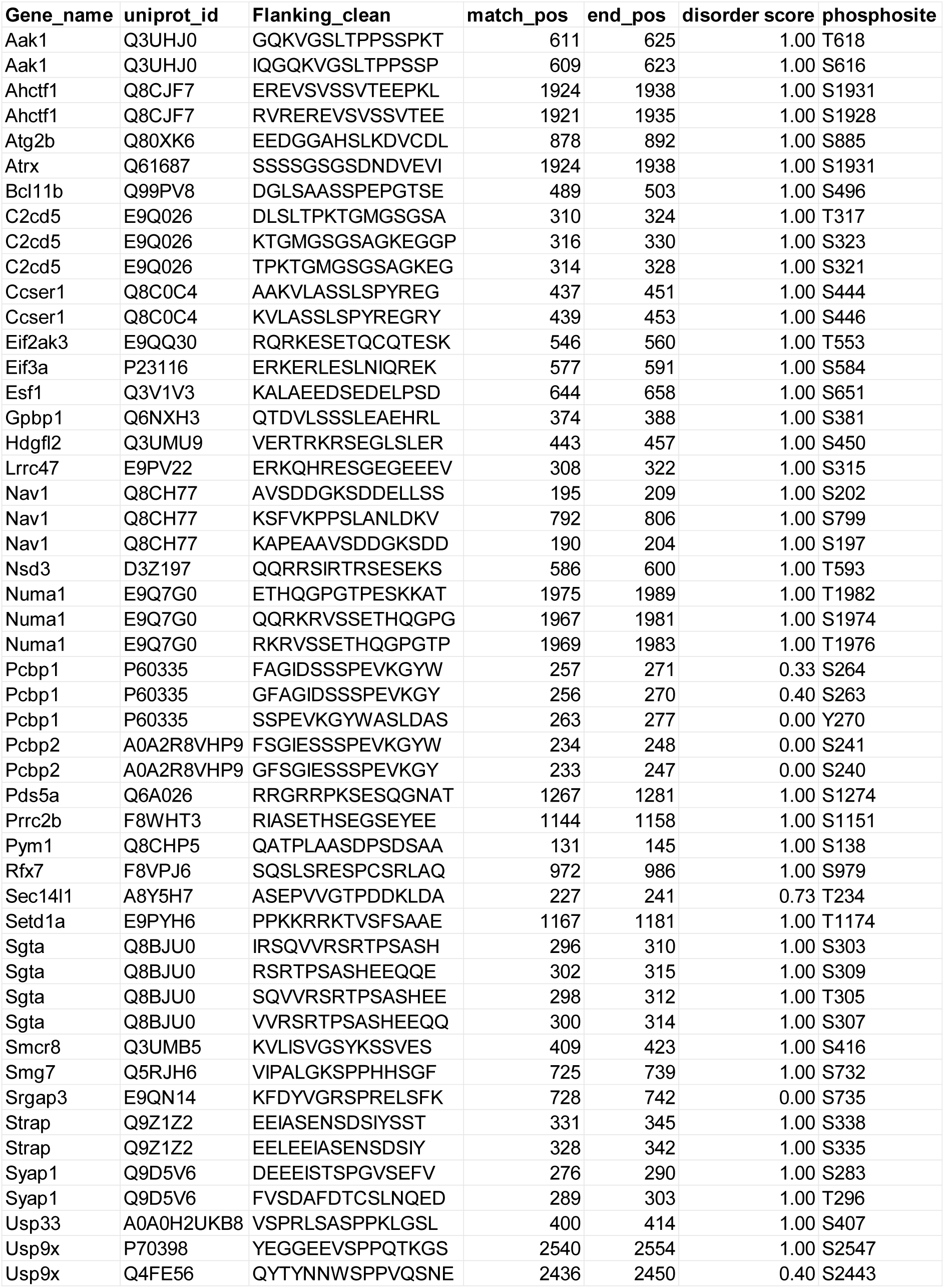
Novel mTORC1 target phosphosites identified in the TSC mouse model. Disorder probability scores for each phosphosite also shown.

| Gene_name | uniprot_id | Flanking_clean | match_pos | end_pos | disorder score | phosphosite |
|---|---|---|---|---|---|---|
| Aak1 | Q3UHJ0 | GQKVGSLTPPSSPKT | 611 | 625 | 1.00 | T618 |
| Aak1 | Q3UHJ0 | IQGQKVGSLTPPSSP | 609 | 623 | 1.00 | S616 |
| Ahctf1 | Q8CJF7 | EREVSVSSVTEEPKL | 1924 | 1938 | 1.00 | S1931 |
| Ahctf1 | Q8CJF7 | RVREREVSVSSVTEE | 1921 | 1935 | 1.00 | S1928 |
| Atg2b | Q80XK6 | EEDGGAHSLKDVCDL | 878 | 892 | 1.00 | S885 |
| Atrx | Q61687 | SSSSGSGSDNDVEVI | 1924 | 1938 | 1.00 | S1931 |
| Bcl11b | Q99PV8 | DGLSAASSPEPGTSE | 489 | 503 | 1.00 | S496 |
| C2cd5 | E9Q026 | DLSLTPKTGMGSGSA | 310 | 324 | 1.00 | T317 |
| C2cd5 | E9Q026 | KTGMGSGSAGKEGGP | 316 | 330 | 1.00 | S323 |
| C2cd5 | E9Q026 | TPKTGMGSGSAGKEG | 314 | 328 | 1.00 | S321 |
| Ccser1 | Q8C0C4 | AAKVLASSLSPYREG | 437 | 451 | 1.00 | S444 |
| Ccser1 | Q8C0C4 | KVLASSLSPYREGRY | 439 | 453 | 1.00 | S446 |
| Eif2ak3 | E9QQ30 | RQRKESETQCQTESK | 546 | 560 | 1.00 | T553 |
| Eif3a | P23116 | ERKERLESLNIQREK | 577 | 591 | 1.00 | S584 |
| Esf1 | Q3V1V3 | KALAEEDSEDELPSD | 644 | 658 | 1.00 | S651 |
| Gpbp1 | Q6NXH3 | QTDVLSSSLEAEHRL | 374 | 388 | 1.00 | S381 |
| Hdgfl2 | Q3UMU9 | VERTRKRSEGLSLER | 443 | 457 | 1.00 | S450 |
| Lrrc47 | E9PV22 | ERKQHRESGEGEEEV | 308 | 322 | 1.00 | S315 |
| Nav1 | Q8CH77 | AVSDDGKSDDELLSS | 195 | 209 | 1.00 | S202 |
| Nav1 | Q8CH77 | KSFVKPPSLANLDKV | 792 | 806 | 1.00 | S799 |
| Nav1 | Q8CH77 | KAPEAAVSDDGKSDD | 190 | 204 | 1.00 | S197 |
| Nsd3 | D3Z197 | QQRRSIRTRSESEKS | 586 | 600 | 1.00 | T593 |
| Numa1 | E9Q7G0 | ETHQGPGTPESKKAT | 1975 | 1989 | 1.00 | T1982 |
| Numa1 | E9Q7G0 | QQRKRVSSETHQGPG | 1967 | 1981 | 1.00 | S1974 |
| Numa1 | E9Q7G0 | RKRVSSETHQGPGTP | 1969 | 1983 | 1.00 | T1976 |
| Pcbp1 | P60335 | FAGIDSSSPEVKGYW | 257 | 271 | 0.33 | S264 |
| Pcbp1 | P60335 | GFAGIDSSSPEVKGY | 256 | 270 | 0.40 | S263 |
| Pcbp1 | P60335 | SSPEVKGYWASLDAS | 263 | 277 | 0.00 | Y270 |
| Pcbp2 | A0A2R8VHP9 | FSGIESSSPEVKGYW | 234 | 248 | 0.00 | S241 |
| Pcbp2 | A0A2R8VHP9 | GFSGIESSSPEVKGY | 233 | 247 | 0.00 | S240 |
| Pds5a | Q6A026 | RRGRRPKSESQGNAT | 1267 | 1281 | 1.00 | S1274 |
| Prrc2b | F8WHT3 | RIASETHSEGSEYEE | 1144 | 1158 | 1.00 | S1151 |
| Pym1 | Q8CHP5 | QATPLAASDPSDSAA | 131 | 145 | 1.00 | S138 |
| Rfx7 | F8VPJ6 | SQSLSRESPCSRLAQ | 972 | 986 | 1.00 | S979 |
| Sec14l1 | A8Y5H7 | ASEPVVGTPDDKLDA | 227 | 241 | 0.73 | T234 |
| Setd1a | E9PYH6 | PPKKRRKTVSFSAAE | 1167 | 1181 | 1.00 | T1174 |
| Sgta | Q8BJU0 | IRSQVVRSRTPSASH | 296 | 310 | 1.00 | S303 |
| Sgta | Q8BJU0 | RSRTPSASHEEQQE | 302 | 315 | 1.00 | S309 |
| Sgta | Q8BJU0 | SQVVRSRTPSASHEE | 298 | 312 | 1.00 | T305 |
| Sgta | Q8BJU0 | VVRSRTPSASHEEQQ | 300 | 314 | 1.00 | S307 |
| Smcr8 | Q3UMB5 | KVLISVGSYKSSVES | 409 | 423 | 1.00 | S416 |
| Smg7 | Q5RJH6 | VIPALGKSPPHHSGF | 725 | 739 | 1.00 | S732 |
| Srgap3 | E9QN14 | KFDYVGRSPRELSFK | 728 | 742 | 0.00 | S735 |
| Strap | Q9Z1Z2 | EEIASENSDSIYSST | 331 | 345 | 1.00 | S338 |
| Strap | Q9Z1Z2 | EELEEIASENSDSIY | 328 | 342 | 1.00 | S335 |
| Syap1 | Q9D5V6 | DEEEISTSPGVSEFV | 276 | 290 | 1.00 | S283 |
| Syap1 | Q9D5V6 | FVSDAFDTCSLNQED | 289 | 303 | 1.00 | T296 |
| Usp33 | A0A0H2UKB8 | VSPRLSASPPKLGSL | 400 | 414 | 1.00 | S407 |
| Usp9x | P70398 | YEGGEEVSPPQTKGS | 2540 | 2554 | 1.00 | S2547 |
| Usp9x | Q4FE56 | QYTYNNWSPPVQSNE | 2436 | 2450 | 0.40 | S2443 |

### mTORC1 target phosphosites are predominantly located within intrinsically disordered regions (IDRs)

Using the identified mTORC1 targets we aimed to develop a further understanding of the mTORC1-dependent phosphorylation landscape in neurodevelopment. mTOR has a strong bias towards serine as the phosphoacceptor residue (77% serine, 23% threonine), and a bias towards proline at the +1 position with glutamic acid, phenylalanine, tyrosine and glutamine also favoured at +1 (*24, 34, 36*). Analysis of mouse embryonic brain mTORC1 target phosphosites showed that serine was the dominant phosphoacceptor residue (77%), with proline most common (26%) at the +1 position and some enrichment for leucine, glutamic acid, glutamine and tyrosine (Figure 2D). Thus, the consensus phosphorylation motif in the mTORC1 targets identified in the mouse TSC model is consistent with the previously established mTORC1 substrate consensus motif.

We next investigated the wider context surrounding the mTORC1 target phosphosites identified in the mouse TSC model. Phosphorylated residues are most frequently found within intrinsically disordered regions (IDRs), flexible regions lacking a conserved three-dimensional structure (*37, 38*). We determined the disorder score within the 14 residues surrounding the 81 mTORC1 target phosphosites. This analysis showed that the majority of phosphosites, such as T1174 in SETD1A, a histone methyltransferase that regulates excitatory synaptic transmission (*39*), were in IDRs (Figure 2E; Table 1; Supplementary table S10). Importantly, using the disorder score, we also identified phosphosites that are close to or within structured conserved domains. These include S735 in the C-terminal SH3 domain of SRGAP3, a Rho GTPase-activating protein that regulates neuronal differentiation (*39*), and S2443 in the C-terminal domain of unknown function (DUF) of USP9X, a deubiquitylating enzyme (DUB) with a variety of roles in neurodevelopment (*40*) (Figure 2F, G; Table 1; Supplementary table S10). Phosphorylation by mTORC1 may therefore modify the function of novel mTORC1 targets such as SRGAP3 and USP9X via these conserved domains.

### Proteomic and phosphoproteomic analysis of mTORC1 activation and inhibition in a cellular model of human neurodevelopment

Mouse neurodevelopmental models capture the complexity of brain development but can be limited by species specific differences. Therefore, to model mTORC1 activation in a cellular model of human neurodevelopment we generated TSC1 KO hiPSCs containing a deletion in *TSC1* exon 17 (Figure 3A, B; Supplementary figure S4) (*41*). We differentiated TSC1 KO and isogenic control hiPSCs into cortical NPCs using directed differentiation via dual SMAD inhibition (*42*), which was confirmed by expression of the pluripotency marker OCT4 and the NPC markers PAX6, TUBB3 and Nestin at day 20 (Figure 3C; Supplementary figure S5). Western blot analysis confirmed the absence of TSC1 and reduced TSC2 expression in TSC1 KO NPCs (Figure 3D, G, H). TSC1 KO NPCs also had significantly increased expression of RPS6 S240/244, S6K1 T390 and a trend towards increased 4E-BP T37/46, while rapamycin treatment abrogated the increase in RPS6 S240/244 and S6K1 T390 phosphorylation but did not affect 4E-BP T37/46 (Figure 3E, F, I-K), which can be insensitive to rapamycin in cultured cells (*43, 44*). Phosphorylation of the mTORC2 target PKCα S657 was unchanged in TSC1 KO NPCs (Figure 3F, L).

**Figure 3.**
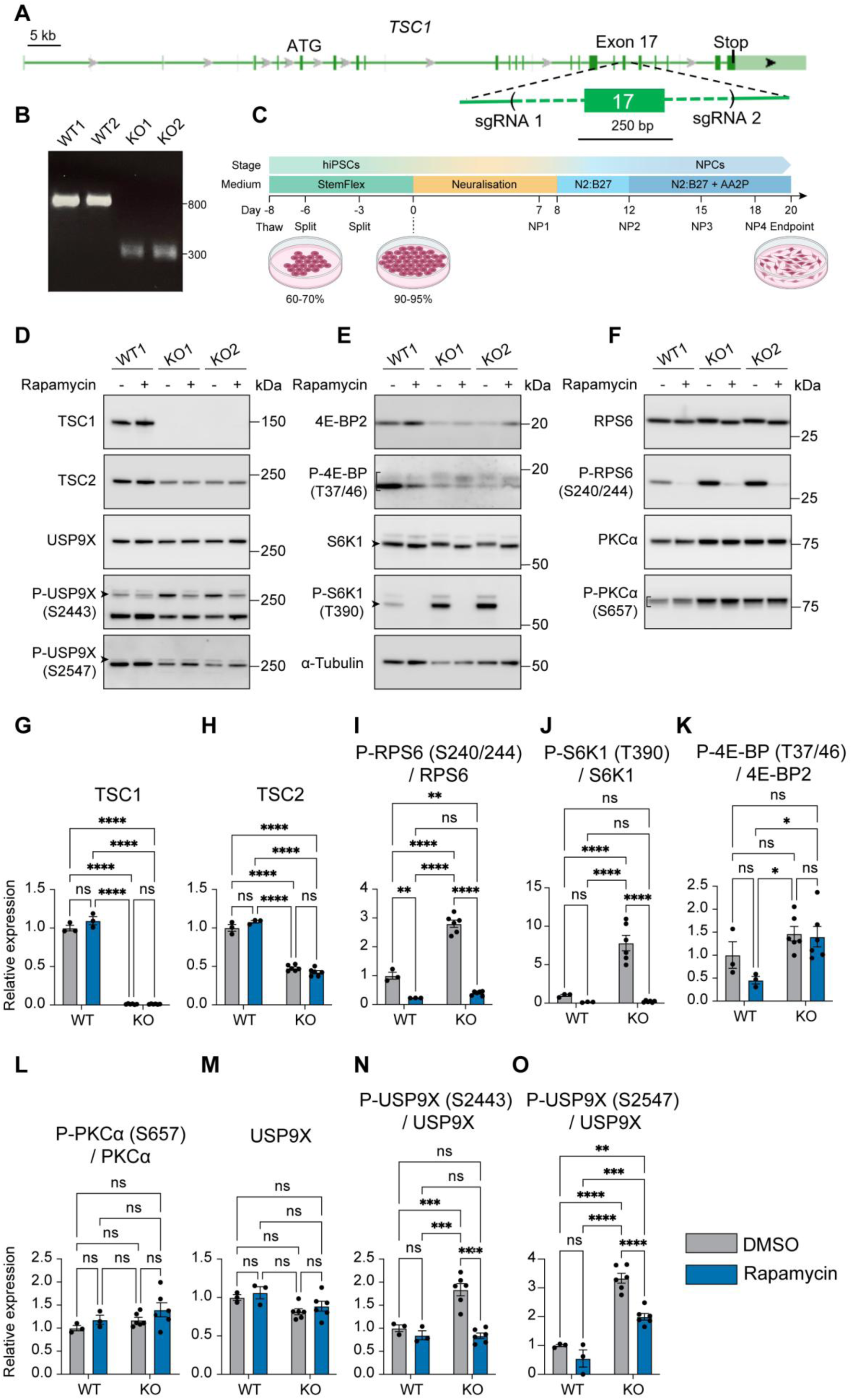
mTORC1 activation in a human cellular model of neurodevelopment. (A) The *TSC1* gene region showing the location of the gRNAs used to delete exon 17 in hiPSCs. (B) PCR analysis of control and TSC1 KO iPSCs to confirm deletion of exon 17. The 800bp band present in wild-type (WT) controls is absent in TSC1 KO mutant hiPSCs. (C) Schematic of the NPC differentiation protocol. NP: neural passage. (D) Western blot analysis of control and TSC1 KO NPCs treated with vehicle or rapamycin shows phosphorylation of USP9X S2443 and S2547 are mTORC1 activity dependent. (E, F) Western blot analysis of P-4E-BP T37/T46 and P-S6K1 T390 (E), P-RPS6 S240/S244 and P-PKCα S657 (F) in control and TSC1 KO NPCs treated with vehicle or rapamycin. (G, H) Quantification of TSC1 and TSC2 expression levels in control and TSC1 KO NPCs treated with vehicle or rapamycin. (I-L) Quantification of P-4E-BP T37/T46 (I), P-S6K1 T390 (J), P-RPS6 S240/S244 (K) and P-PKCα S657 (L) levels in control and TSC1 KO NPCs treated with vehicle or rapamycin. (M-O) Quantification of total USP9X (M), P-USP9X S2443 (N) and P-USP9X S2547 (O) levels in control and TSC1 KO NPCs treated with vehicle or rapamycin. WT n=3, TSC1 KO n=3; vehicle treated TSC1 KO n=6, rapamycin treated TSC1 KO n=6. Data are represented as mean ± SEM and were analysed using the students t test. ns not significant, * p < 0.05, **p < 0.01, ***p < 0.001****, p < 0.0001.

Quantitative proteomics of TSC1 KO NPCs compared to controls (Supplementary figure S6A) showed that the expression of a large number of proteins (3297) was significantly changed in (Figure 4A; Supplementary tables S12, S13). GO analysis showed that proteins with increased expression in TSC1 KO NPCs were most enriched for GO terms involving the cytoskeleton and the KEGG pathway term ribosome, while proteins with significantly decreased expression were enriched for GO terms involving chromatin organisation and RNA metabolic processes (Supplementary figure S6C-E). Moreover, 59 large and small ribosomal subunit proteins had significantly increased expression in TSC1 KO NPCs (Supplementary tables S12, S13), while none were decreased, reflecting the role of mTORC1 in ribosome biogenesis and protein synthesis.

**Figure 4.**
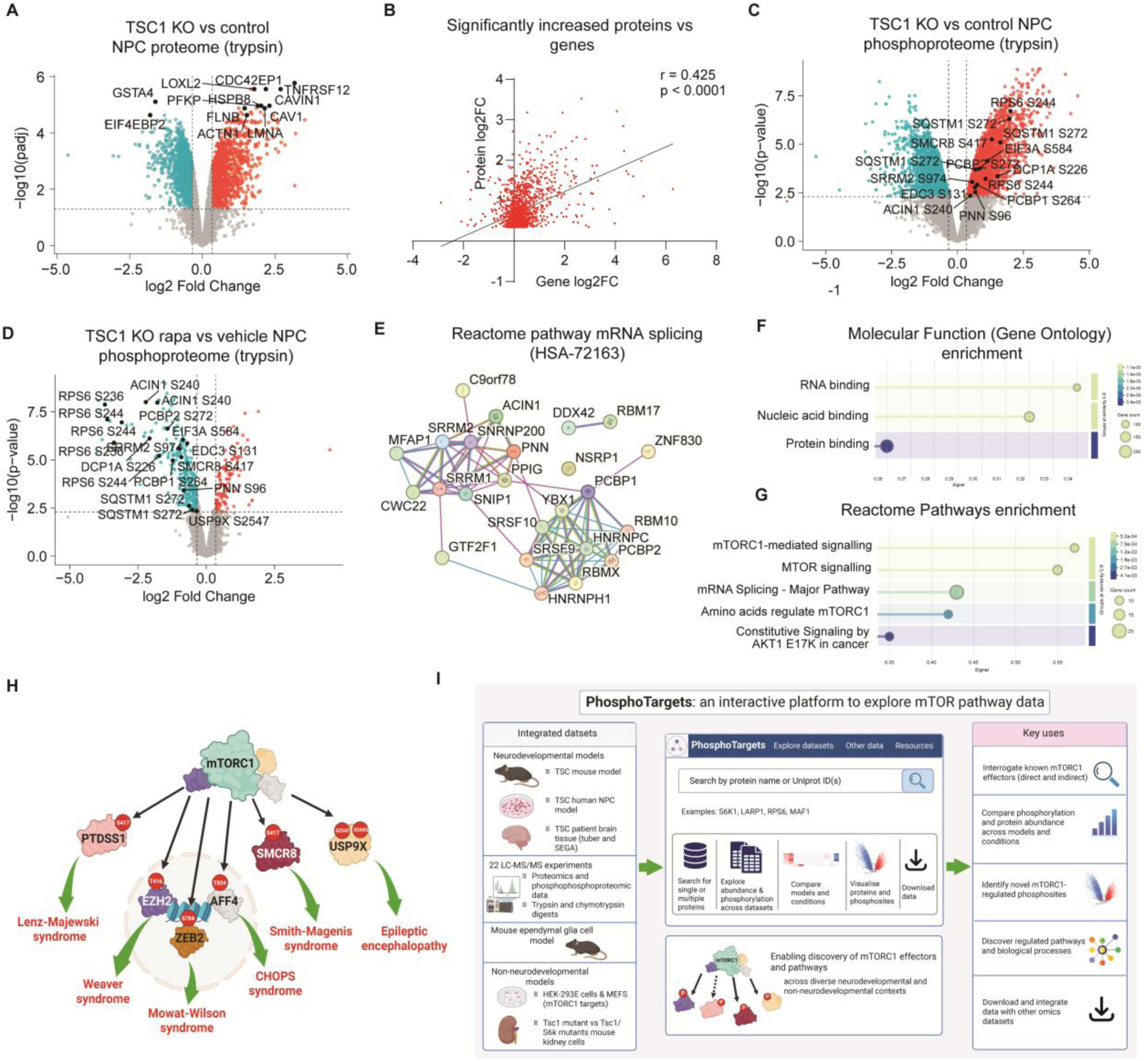
Identification of novel mTORC1 targets in human neurodevelopment. (A) Volcano plot representing proteomic data from control versus TSC1 KO NPCs (trypsin digest). Control n=3 biological replicates, TSC1 KO n=5 biological replicates. (B) Pearson correlation of significantly increased proteins (trypsin digest) in control versus TSC1 KO NPCs with gene expression. For snRNA-seq Control n=4 biological replicates, TSC1 KO n=4 biological replicates. (C) Volcano plot representing phosphoproteomic data from control versus TSC1 KO NPCs (trypsin digest). Known and novel mTORC1 targets are labelled. Control n=3, TSC1 KO n=5. (D) Volcano plot representing proteomic data from TSC1 KO NPCs treated with vehicle or rapamycin (trypsin digest). Known and novel mTORC1 targets are labelled. Vehicle treated TSC1 KO n=5 biological replicates, rapamycin treated TSC1 KO n=5 biological replicates. (E) Proteins representing phosphoproteins significantly enriched for the Reactome pathway mRNA splicing (HSA-72163) among the phosphoproteins significantly decreased in the TSC1 KO NPCs treated with rapamycin. (F, G) GO molecular function terms (F) and Reactome pathways (G) significantly enriched in phosphoproteins with decreased expression in rapamycin treated versus vehicle treated TSC1 KO NPCs. (H) Schematic showing examples of mTORC1 targets in human NPCs associated with NDDs. (I) Schematic of PhosphoTargets interactive resource. For volcano plots proteins or phosphopeptides with significantly increased expression coloured red and significantly decreased expression coloured blue.

To investigate how much of the proteomic changes in TSC1 KO NPCs were due to transcriptional regulation we performed RNA-sequencing (RNA-seq) of TSC1 KO and control NPCs (Supplementary table S20). The expression of mRNAs and the proteins encoded by these transcripts in TSC1 KO NPCs showed a highly significant positive correlation (r = 0.37, p<0.0001). Moreover, levels of proteins that had significantly increased expression correlated very strongly with their cognate mRNA expression in TSC1 KO NPCs (r = 0.425, p<0.001, Figure 4B). These data indicate that the largescale alterations in protein expression in TSC1 KO NPCs originate predominantly at the level of transcription.

4179 phosphopeptides, corresponding to 1794 proteins, were significantly increased in TSC1 KO NPCs (Figure 4C; Supplementary tables S14, S15). The results of GO analysis of these proteins were similar to GO analysis of the proteins with significantly increased expression (Supplementary figure S6C, F). Moreover, 30% of the proteins corresponding to significantly increased phosphopeptides in the TSC1 KO NPCs were also significantly increased in the proteomics dataset (Supplementary figure S6G). Thus, a minority proportion of the mTORC1 phosphosites identified in analysis of TSC1 KO versus control NPCs are potentially increased due to higher protein expression.

By contrast to TSC1 KO versus control NPCs, rapamycin treatment of TSC1 KO NPCs (Supplementary figure S6B) did not significantly alter expression at the protein level (Supplementary tables S16, S17). However, rapamycin treatment of TSC1 KO NPCs caused a significant decrease in a large number of phosphopeptides (695), corresponding to 323 proteins (Figure 4D; Supplementary tables S18, S19). These proteins were significantly enriched for GO terms RNA binding and nucleic acid binding and for Reactome pathways mTORC1-mediated signalling and mRNA splicing (Figure 4E-G). Thus, during acute mTORC1 inhibition largescale signalling changes in human NPCs occur at the level of protein phosphorylation. Moreover, these data show that splicing factors are a major target of mTORC1 in human NPCs.

### Novel mTORC1 targets are conserved in neurodevelopment

To select high confidence mTORC1 targets in the human TSC model we focused on phosphosites that were increased in TSC1 KO versus control NPCs *and* decreased in rapamycin treated TSC1 KO NPCs. These criteria selected 142 phosphosites, representing 79 proteins (Supplementary table S21). Six of these proteins are known direct mTORC1 substrates, either with previously identified phosphosites (PRAS40/AKT1S1 S212; ACIN1 S240; LARP1 S766, S774, S849, S851, S868, T1071; LPIN S106; PRKAA1 S356; ZNRF2 S145) or novel phosphosites (EEF2K S74, SQSTM1 S266, T269, S272, S328, S332) (Supplementary table S21) (*34*). Nine other proteins are known indirect mTORC1 targets via S6K (EIF4B, CAD, DNAJC2, EEF2K, EIF4G1, PRKAA2, RICTOR, RPS6) or AMPK (C18ORF25) (Supplementary table S21) (*35, 45*). Similar to the mTORC1 targets identified in mice, human NPC mTORC1 target phosphosites had a strong preference for serine as the phosphoacceptor and proline at the +1 position and were highly enriched within IDRs (Supplementary figure S6H; Supplementary table S21). Moreover, a small minority of phosphosites were within or close to conserved domains, such as S216 located between adjacent LIM domains in the axon pathfinding actin-binding protein ABLIM1 (*46*) (Supplementary figure S6I; Supplementary table S21), and may thus modify domain function.

Of the 32 novel mTORC1 target proteins identified in the TSC mouse model 9 proteins (AHCTF1, EIF3A, HDGFL2, NSD3, PCBP1, PCBP2, SGTA, SMCR8, USP9X) and 7 phosphosites (EIF3A S584; HDGFL2 S454; PCBP1 S263, S264; PCBP2 S272; SMCR8 S417, USP9X S2547) were also mTORC1 targets in the human TSC NPC model. Thus, even though the phosphoproteomic analyses were performed from remarkably different TSC models, mouse embryonic brain and human NPCs, we were able to identify conserved mTORC1 targets in mouse and human neurodevelopment.

### Novel mTORC1 targets are associated with human neurodevelopmental disorders

We next investigated the potential for the phosphoproteomic resource to connect mTOR signalling to broader NDD networks. Mutations in genes that activate mTORC1 cause a group of rare NDDs known as mTORopathies, characterised by malformations of cortical development and epilepsy (*19, 47, 48*). mTORopathies share features such as epilepsy with other rare genetic NDDs. We therefore hypothesised that the molecular mechanisms underlying mTORopathies such as TSC might connect to the proteins mutated in other rare NDDs. Interestingly, of the 32 novel mTORC1 targets identified in the mouse TSC model 12 are associated with rare NDDs, including Smith-Magenis syndrome (SMRC8), Intellectual disability with autism spectrum disorder (PCBP1) and epileptic encephalopathy (USP9X) (Table 2). Of the 64 novel mTORC1 targets identified in the TSC human NPC model 13 are associated with rare NDDs including epileptic encephalopathy (USP9X), Smith-Magenis syndrome (SMRC8), CHOPS syndrome (AFF4), Weaver syndrome (EZH2), Mowat Wilson syndrome (ZEB2) and Lenz-Majewski syndrome (PTDSS1) (Figure 4H; Table 3). These disorders share features such as brain malformations, epilepsy and autism spectrum disorder with mTORopathies, suggesting that there may be common molecular mechanisms across these rare NDDs that are coordinated by mTORC1 during neurodevelopment (Figure 4H).

**Table 2.** Novel mTORC1 targets identified in the TSC mouse model are associated with rare NDDs.

| **Gene** | **Cellular functions** | **NDD** | **Reference** |
|---|---|---|---|
| *EIF3A* | Translation initiation | Cardiovascular, craniofacial, and neurodevelopmental disorder | (*86*) |
| *ATRX* | Transcriptional regulation and chromatin remodeling | ATR-X syndrome (Alpha-thalassemia X-linked intellectual disability | (*87*) |
| *SMCR8* | Autophagy | Smith-Magenis syndrome | (*88*) |
| *SRGAP3* | Cytoskeletal reorganisation | ID, ASD, schizophrenia risk | (*89*) |
| *USP9X* | Deubiquitination, cell cycle, cell survival, cytoskeleton dynamics | X-linked intellectual disability | (*54*) |
| *BCL11B* | Transcription regulation | NDDs with immuno-deficiency, Cranyostosis | (*90*) |
| *RFX7* | Transcription regulation | ID, ASD, behavioural abnormalities | (*90*) |
| *SETD1A* | Transcription regulation, epigenetic regulation, DNA repair | Developmental delay, intellectual disability, schizophrenia risk, speech delay, and behavioural phenotypes | (*91*) |
| *PCBP1* | Transcription regulation, translation regulation | Intellectual disability with autism spectrum disorder | (*92*) |
| *PCBP2* | Transcription regulation, translation regulation | NDD associated with motor and language delay | (*93*) |
| *PDS5A* | Epigenetic regulation, DNA repair, DNA replication | Cohesino-pathies (developmental delay and congenital anomalies) | (*94*) |
| *SYNGAP1* | Promotes mTORC2-mediated phosphorylation of AKT1 | SYNGAP1-related disorder | (*95*) |

**Table 3.** Novel mTORC1 targets identified in the human TSC NPC model are associated with rare NDDs.

| Gene | Cellular functions | NDD | Reference |
|---|---|---|---|
| *AFF4* | Transcriptional regulation | CHOPS syndrome: ID, hypotonia, craniofacial anomalies, growth delay | (*96*) |
| *EZH2* | Histone methyltransferase | Weaver syndrome: overgrowth, ID, macrocephaly | (*97*) |
| *ZEB2* | Transcriptional repressor | Mowat-Wilson syndrome: severe ID, epilepsy, Hirschsprung disease | (*98*) |
| *TRIP12* | E3 ligase | Syndromic ID, ASD, hypotonia, epilepsy | (*99*) |
| *PUM1* | RNA-binding protein | PUM1-associated developmental disability, ataxia, and seizures | (*100*) |
| *PTDSS1* | Phosphatidylserine synthase, located in the mitochondria-associated endoplasmic reticulum membrane | Lenz-Majewski syndrome: hyperostotic dwarfism, ID, craniofacial anomalies | (*101*) |
| *CAD* | Carbamoyl-phosphate synthetase, de novo synthesis of pyrimidine nucleotides | Developmental and epileptic encephalopathy, treatable metabolic defect | (*102*) |
| *SLC20A2* | Type III sodium-dependent phosphate transporter 2 | Familial idiopathic basal ganglia calcification | (*103*) |
| *RB1* | Tumour suppressor, inhibits G1/S transition | ID and facial dysmorphic features | (*104*) |
| *CAPRIN1* | RNA-binding protein, regulates<br>mRNA transport, translation and stability | ID, speech delay, ASD | (*105*) |
| *SMCR8* | Autophagy | Smith-Magenis syndrome | (*88*) |
| *PI4K2A* | PI4-kinase, located on the trans-Golgi network and endosomes | Epilepsy, developmental delay | (*106*) |
| *USP9X* | Deubiquitination, cell cycle, cell survival, cytoskeleton dynamics | X-linked intellectual disability | (*54*) |

### An interactive resource for interrogation of mTOR signalling pathway mechanisms in cell and animal models and in human disease

The proteomic and phosphoproteomic data in this study and our recent analysis of TSC patient brain tissue (*49*) represent 22 individual LC-MS/MS experiments. To aid in the analyses of these data by the scientific community we created an interactive platform: PhosphoTargets (https://phosphotargets.com). Using PhosphoTargets, data from the TSC mouse model and TSC human NPC model can be interrogated by searching for single or multiple protein names or Uniprot ID (Figure 4I). Proteomic and phosphoproteomic data from TSC patient tuber and subependymal giant cell astrocytoma (SEGA) brain tissue can be interrogated alongside the mouse and NPC data. We also included recent phosphoproteomic data from rapamycin treated mouse ependymal glial cells (*50*) (Figure 4I). For comparison with non-neurodevelopmental models we have included phosphoproteomic identification of mTORC1 targets in HEK-293E cells and MEFs (*24*), as well as Tsc1 mutant versus Tsc1/S6k mutant mouse kidney cell phosphoproteomic data (*35*) (Figure 4I). This interactive platform can be utilised for analysing changes in known mTORC1 direct and indirect effectors in different neurodevelopmental and non-neurodevelopmental contexts. PhosphoTargets can further be used to identify novel mTORC1-regulated phosphosites as potential key effectors and, more broadly, to discover novel cellular processes regulated by mTORC1.

### USP9X is a novel conserved mTORC1 substrate in neurodevelopment

To validate our mTORC1 phosphoproteomic resource and the selection of novel mTORC1 targets and demonstrate their utility, we focused on the highly conserved DUB USP9X. USP9X was identified in the mouse and human TSC models as a novel mTORC1 target (Figure 2A, B; Figure 4D; Table 1; Supplementary table S21). USP9X has been shown to physically interact with the mTORC1 component Raptor (*51*). USP9X is also essential for early embryogenesis and has key roles in neurodevelopment (*40*). Loss-of-function mutations in USP9X cause neurodevelopmental disorders including epileptic encephalopathy and X-linked recessive nonsyndromic intellectual developmental disorder-99, associated with intellectual disability, epilepsy and autism (*52–54*).

Our phosphoproteomic analysis identified USP9X S2443 and S2547 as mTORC1 targets in the TSC mouse model and USP9X S2547 in the human NPC TSC model (Figure 5A; Table 1; Supplementary table S21). Interestingly, phosphorylation of S2547 regulates the DUB activity of USP9X (*55*). We generated phospho-specific antibodies against these residues and validated them using USP9X mutants containing alanine substitutions at S2443 and S2547 (Supplementary figure S7). These phospho-specific antibodies were then used to confirm the TSC mouse model phosphoproteomic data by western blot analysis of Tsc1 cKO versus control and Tsc1 cKO rapamycin treated versus vehicle treated mouse brain tissue (Figure 5B-I).

**Figure 5.**
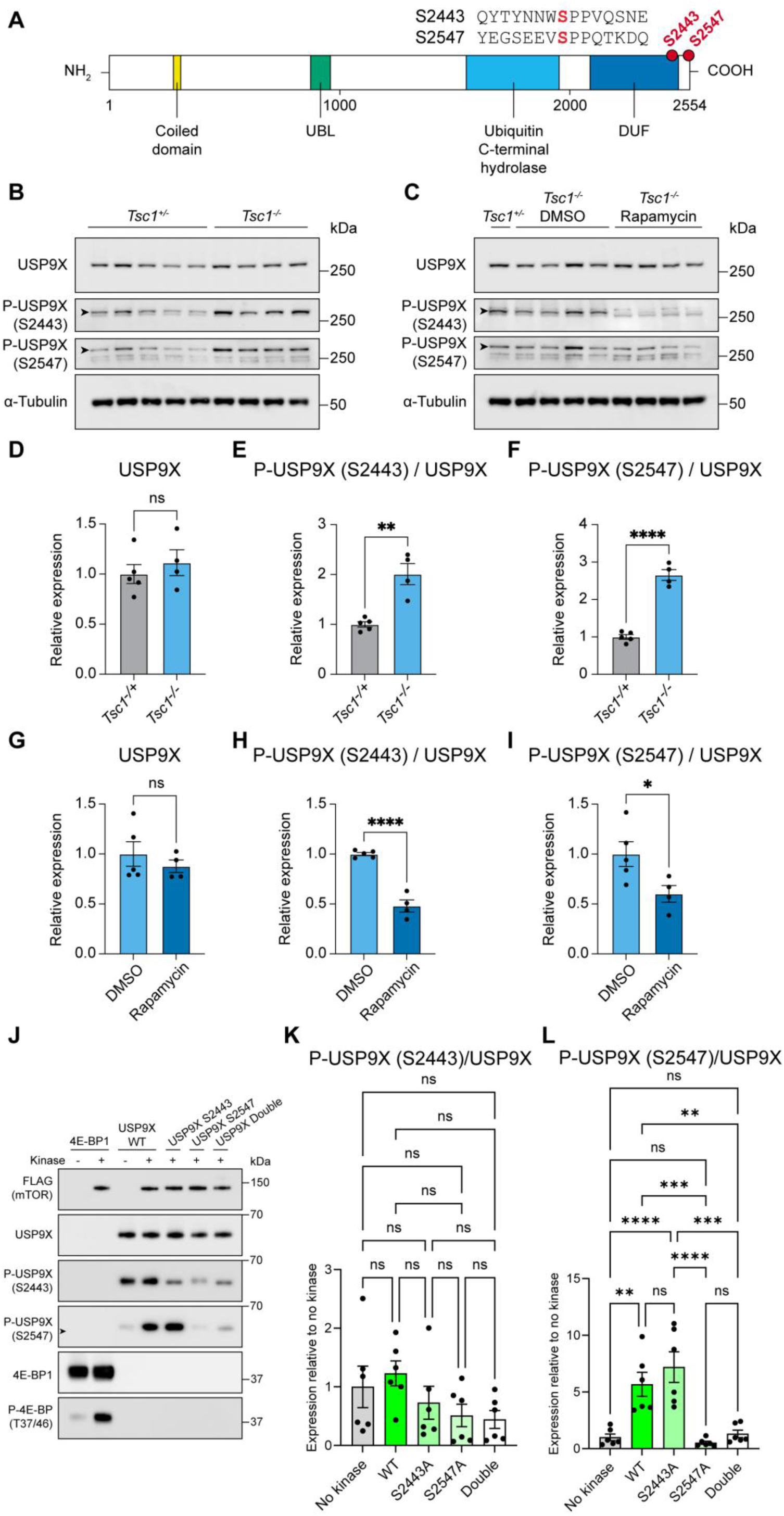
USP9X is a novel target of mTORC1 during mouse neurodevelopment. (A) Schematic of the USP9X domain structure showing the mTORC1-dependent phosphosites identified by phosphoproteomics. UBL: ubiquitin-like domain; DUF: domain of unknown function. Sequence surrounding phosphosites (in red) is shown above. (B, C) Western blot analysis using phospho-specific antibodies shows phosphorylation of USP9X S2443 and S2547 are mTORC1-dependent in E16.5 brain tissue from the Tsc1 cKO versus control (B) and Tsc1 cKO rapamycin treated versus vehicle mouse models (C). (D-F) Quantification of USP9X S2443 phosphorylation in the E16.5 brain tissue from the Tsc1 cKO versus control mouse model. Control n=5 brains, Tsc1 cKO n=4 brains. (G-I) Quantification of USP9X S2443 phosphorylation in the E16.5 brain tissue from the Tsc1 cKO rapamycin treated versus vehicle mouse model. Vehicle treated Tsc1 cKO n=5 brains, rapamycin treated Tsc1 cKO n=4 brains. (J) Western blot analysis of mTOR kinase assays using phospho-specific antibodies against 4E-BP1 T37/46, USP9X S2443 and USP9X S2547 showing S2547 is directly phosphorylated by mTOR. (K, L) Quantification of USP9X S2443 (K) and USP9X S2547 (L) phosphorylation by mTOR. n=6 kinase assays for all conditions. Data are represented as mean ± SEM and were analysed using the students t test. ns not significant, * p < 0.05, **p < 0.01, ***p < 0.001, ****p < 0.0001.

Consistent with the human NPC phosphoproteomics, phosphorylation of USP9X S2547 was significantly increased in TSC1 KO NPCs and decreased by rapamycin treatment (Figure 3D, O). Despite non-significant changes in USP9X S2443 by phosphoproteomics, western blot analysis showed that phosphorylation of S2443 was significantly increased in TSC1 KO NPCs and significantly decreased by rapamycin treatment (Figure 3D, N). These data show that mTORC1-dependent phosphosites in USP9X identified in the TSC mouse model embryonic brain are conserved in a human TSC NPC model.

Both USP9X S2443 and S2547 have a proline at the +1 position (Figure 5A), typical of mTORC1 substrates, and do not have an arginine at the -3 or -5 position, typical of AGC kinase (such as S6K) substrates ((*56*), suggesting these residues may be directly phosphorylated by mTORC1. To test this, we expressed and purified recombinant N-terminal truncated wild-type USP9X, single S2443A and S2547A mutants, as well as double S2443A/S2547A USP9X mutant, and performed *in vitro* kinase assays using recombinant mTOR with recombinant 4E-BP1 as a positive control substrate. As expected, probing with a phospho-specific antibody for 4E-BP1 T37/46 showed 4E-BP1 was strongly phosphorylated only in the presence mTOR (Figure 5J). Although USP9X S2443 phosphorylation was reduced in the mutants this was not significant and S2443 phosphorylation was detected even in the absence of mTOR (Figure 5J, K). However, USP9X S2547 phosphorylation was almost completely dependent on mTOR and was dramatically reduced in the S2547A and double S2443A/S2547A mutants (Figure 5J, L). These *in vitro* data show that USP9X S2547 is a direct substrate of mTOR and that the phosphoproteomic resource can be used to identify novel mTORC1 substrates.

### Loss of phosphorylation of USP9X at S2443 and S2547 causes increased proliferation in human NPCs

Depending on the context, USP9X can promote or inhibit cell proliferation and regulate self-renewal of NPCs (*57–60*). To test whether phosphorylation of USP9X at S2443 and S2547 is functionally important in NPCs we generated serine to alanine double mutant USP9X S2443A/S2547A hiPSCs (Supplementary figure S4B). Genomic sequencing showed the presence of both mutations and western blot analysis confirmed loss of S2443 and S2547 phosphorylation in NPCs differentiated from two independent USP9X S2443A/S2547A hiPSC lines (Figure 6A-C and Supplementary figure S8). USP9X physically interacts with Raptor and maintains Raptor protein levels (*51, 58*). However, Raptor levels and phosphorylation of RPS6 were unchanged in USP9X S2443A/S2547A NPCs (Figure 6C; Supplementary figure S9), indicating that loss of USP9X phosphorylation does not affect upstream mTORC1 activity.

**Figure 6.**
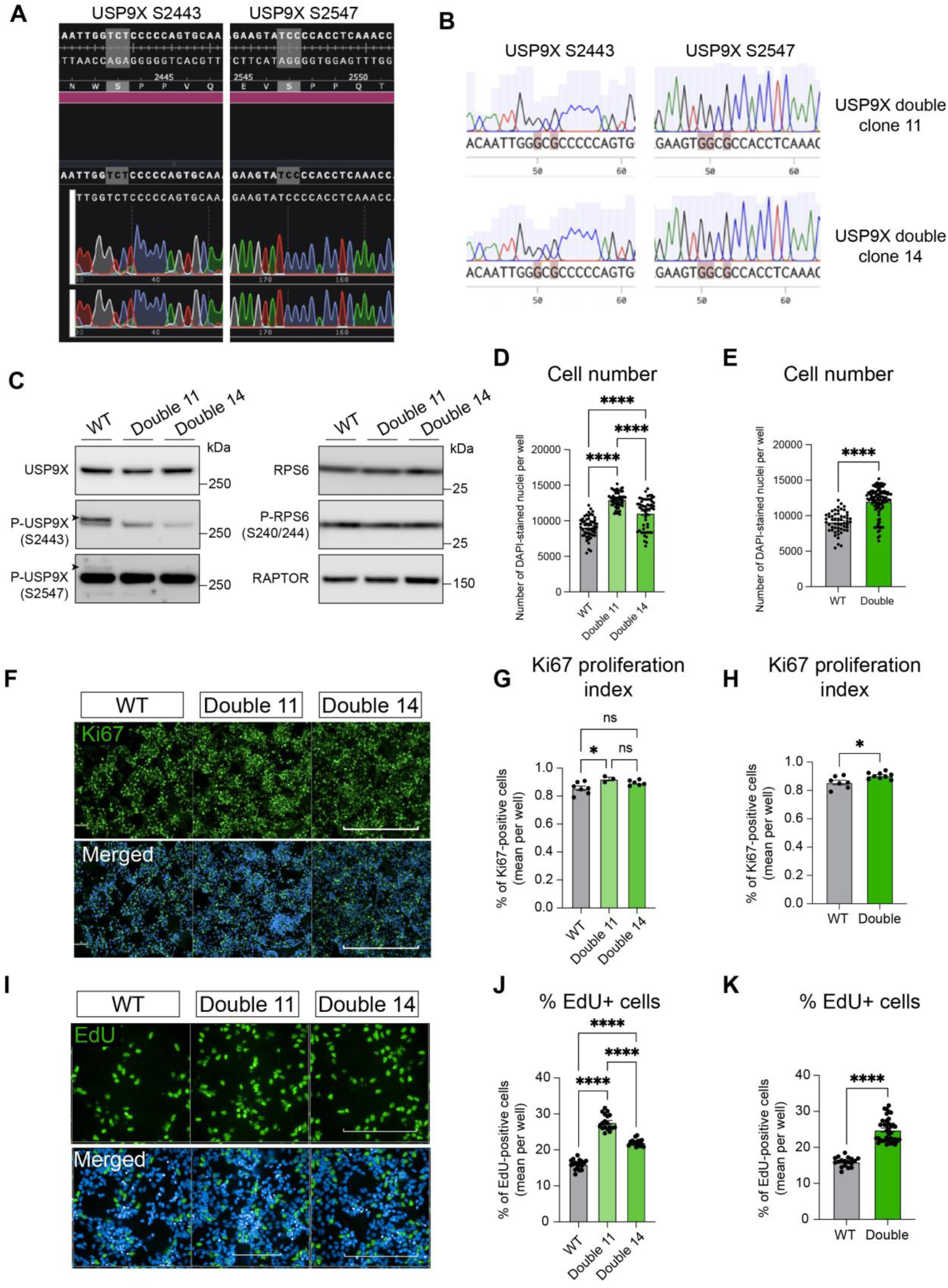
Inhibition of USP9X S2443/S2547 phosphorylation increases proliferation in human NPCs. (A) USP9X genomic DNA sequence around USP9X S2443 and S2547 encoding residues. (B) Sanger sequencing of USP9X iPSC phosphomutant lines double 11 and double 14 showing the presence of point mutations converting serine 2443 and serine 2547 to alanine encoding residues. (C) Western blot analysis of USP9X iPSC phosphomutant lines double 11 and double 14 showing loss of S2443 and S2547 phosphorylation on left hand panel, P-RPS6 S240/2444 and RAPTOR expression on right hand panel. (D, E) Quantification of cell number in control (WT), individual USP9X phosphomutant NPC (double 11 and double 14) lines (D) and both lines combined (E) after growth for 48 hours. n=54 wells per condition. (F) Immunostaining for Ki67 in control (WT) and USP9X phosphomutant NPC (double 11 and double 14) lines. Scale bar: 400µm. (G, H) Quantification of Ki67 expressing cells in control (WT), individual USP9X phosphomutant NPC (double 11 and double 14) lines (G) and both lines combined (H). WT n=7 wells, Double 11 n=3 wells, Double 14 n=6 wells. (I) EdU incorporaton in control (WT) and USP9X phosphomutant NPC (double 11 and double 14) lines. Scale bar: 200µm. (J, K) Quantification of EdU positive cells in control (WT), individual USP9X phosphomutant NPC (double 11 and double 14) lines (J) and both lines combined (K). n=18 wells per condition. Data are represented as mean ± SEM and were analysed using one-way ANOVA or the students t test. ns not significant, *p < 0.05, ****p < 0.0001.

To test whether loss of USP9X phosphorylation at S2443 and S2547 affects proliferation, we plated NPCs at neural passage 4 (NP4) and quantified cell number after 48 hours. USP9X S2443A/S2547A NPC numbers were significantly increased compared to controls after 48 hours (Figure 6D, E). Moreover, USP9X S2443A/S2547A mutant NPCs showed a small but significant increase in Ki67 proliferation index and a 55% increase in EdU incorporation (Figure 6F-K). Therefore, preventing phosphorylation of USP9X mTORC1 target phosphosites S2443 and S2547 was required for regulation of proliferation in NPCs. These data thus validate the phosphoproteomic resource as a means to identify novel and functionally important mTORC1-dependent phosphosites such as USP9X S2443 and S2547.

### Novel mTORC1 targets in human neurodevelopment form interacting networks with functions including regulation of RNA metabolism

As a more global approach, we next aimed to determine the utility of the proteomic and phosphoproteomic data resources to identify novel cellular processes regulated by mTORC1 in neurodevelopment. STRING protein-protein interaction analysis of the 79 mTORC1 targets from the human TSC NPC model identified several networks with shared functions. The largest network consisted of proteins involved in translational regulation, including ribosomal subunits (RPS6, RPL29), translation initiation factors (EIF4B, EIF3A, EIG4G1), the translational GTPase HBS1L, the ribosomal processing factors RRP1B and HBS1L, the RNA binding protein CAPRIN1 and the pre-rRNA processing factor LYAR (Figure 7A). A second network centred around the Ragulator subunit LAMTOR interacting with the mTORC1 subunit AKT1S1 (PRAS40), the vacuolar ATPase ATP6V1H, and the guanine nucleotide exchange factor autophagy regulator SMCR8 (Figure 7B). A third network consisted of proteins regulating mRNA splicing, including the splicing-dependent multiprotein exon junction complex (EJC) component ACIN1, the spliceosome component SRRM2 and the auxiliary EJC component PNN (Figure 7C). Moreover, PCBP1 and PCBP2, proteins with multiple functions in regulation of gene expression, mRNA stability, translation and mRNA splicing, as well as interactors EDC3 and DCP1A that regulate mRNA decapping to control mRNA degradation were also mTORC1 targets in the TSC NPC model (Figure 7C-E; Supplementary table S21). Using the phosphoproteomic data resource we therefore identified novel mTORC1 targets involved in translational regulation and mTORC1 signalling/autophagy in human NPCs. Moreover, our data suggest a novel role for mTORC1 in regulating mRNA splicing during human neurodevelopment.

**Figure 7.**
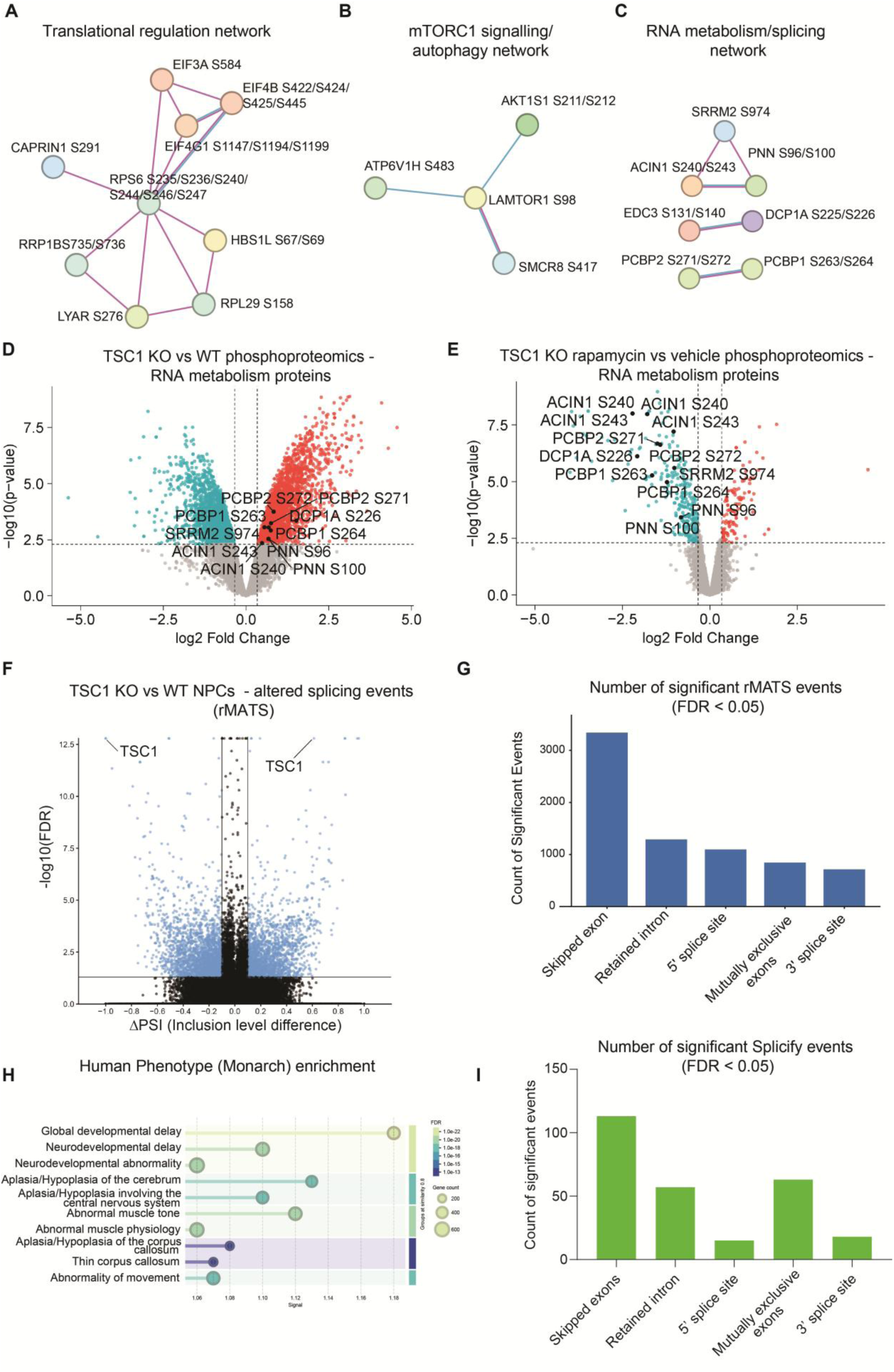
mTORC1 activation perturbs mRNA splicing in human NPCs. (A-C) Novel mTORC1 target protein interaction networks identified in the human TSC model. (D) Volcano plot representing phosphoproteomic data from control versus TSC1 KO NPCs (trypsin digest). Novel RNA metabolism mTORC1 targets are labelled. Control n=3, TSC1 KO n=5. (E) Volcano plot representing proteomic data from TSC1 KO NPCs treated with vehicle or rapamycin (trypsin digest). Novel RNA metabolism mTORC1 targets are labelled. Vehicle treated TSC1 KO n=5 biological replicates, rapamycin treated TSC1 KO n=5 biological replicates. (F) Volcano plot showing significantly different splicing events (blue dots) identified by rMATS in TSC1 KO versus control NPCs. (G) Number of significant rMATS splicing events of each type in TSC1 KO versus control NPCs. Control n=4, TSC1 KO n=4. (H) Human phenotype (Monarch) analysis of genes with altered splicing in TSC1 KO versus control NPCs. (I) Number of significant Splicify splicing events of each type in TSC1 KO versus control NPCs.

### mTORC1 activation perturbs mRNA splicing in human NPCs

In the mouse TSC model four proteins involved in RNA metabolism were identified as novel mTORC1 targets, while a network of novel mTORC1 targets regulating mRNA splicing was identified in the human TSC NPC model (Figure 2C, Figure 4E-G, Figure 7C-E, Supplementary tables S11, S21). Together these data suggest a novel role for mTORC1 in regulation of mRNA splicing during neurodevelopment. To test this possibility we analysed our TSC1 KO NPC RNA-seq data using Replicate Multivariate Analysis of Transcript Splicing (rMATS), a splicing analysis pipeline that detects differences in specific types of splice event including alternative 3’ or 5’ splice site, mutually exclusive exons, retained introns and skipped exons (*61*). Deletion of exon 17 in the TSC1 KO NPCs (Figure 3A) should alter splicing. rMATS identified *TSC1* as the gene with the most significantly altered splicing and several thousand significant splicing events with a meaningful ΔPSI, with skipped exons the most common splicing alteration in TSC1 KO NPCs (Figure 7F, G; Supplementary table S22). IsoformSwitchAnalyzeR analysis (*62*) also confirmed increased usage of a TSC1 isoform truncated at exon 17 that is sensitive nonsense mediated decay (Supplementary figure S10A). 3232 genes had at least one altered splicing event in TSC1 KO NPCs and GO molecular function analysis of these genes showed the strongest enrichment for RNA binding, while Reactome pathways analysis showed strongest enrichment for metabolism of RNA (Supplementary figure S10B, C; Supplementary table S22).

Our recent analysis of TSC patient SEGA brain tissue also showed increased phosphorylation of RNA splicing proteins and largescale changes in mRNA splicing (*49*). Around a third of the genes whose splicing were altered in SEGAs also had altered in TSC1 KO NPCs. Interestingly, human phenotype (Monarch) analysis of the genes whose splicing was affected in TSC1 KO NPCs showed highly significant enrichment for genes associated with global developmental and neurodevelopmental delay/abnormality, aplasia/hypoplasia of the cerebrum and thin corpus callosum (Figure 7H). These data are consistent with perturbed mRNA splicing contributing to the aberrant neurodevelopmental effects of mTORC1 activation.

We next used the Splicify pipeline (*63*) to determine whether there was evidence of consistent splicing alterations at both the RNA and protein level in TSC1 KO NPCs. Splicify integrates RNA-seq data and LC-MS/MS data to identify protein isoforms resulting from differential splicing between two conditions. By integrating the TSC1 KO NPC RNA-seq and proteomic data Splicify identified 266 significant splicing events at the RNA level with consistent significant changes at the protein level in TSC1 KO NPCs (alternative 3’ or 5’ splice site, mutually exclusive exons, retained introns and skipped exons) (Supplementary table S23). Similar to the rMATS analysis at the RNA level (Figure 7G), skipped exons were the most common splicing alteration at the protein level in TSC1 KO NPCs (Figure 7I). These data demonstrate how the proteomic resource can be integrated with other omics datasets to identify novel roles for mTORC1 signalling in human NPCs.

Overall, by revealing mTORC1 downstream effectors in mouse brain and human cellular neurodevelopmental models our study exemplifies how these resources can be utilised, ranging from identification of new single substrate phosphorylation to global effects on mRNA splicing. Moreover, PhosphoTargets provides a platform for easy analysis of mTORC1 signalling in neurodevelopmental models and human disease, enabling hypothesis generation or testing of novel roles for mTORC1 signalling.

## Discussion

Although mTORC1 has dozens of direct substrates and indirect targets (*24–26, 34*), our current knowledge of mTORC1 in nervous system development is based on analysis of a small number of established effectors (*64, 65*). The research community therefore lacks crucial knowledge of the proteomic and phosphoproteomic landscape of mTORC1 signalling in neurodevelopment. The resources we provide here address this knowledge gap by providing deep proteomic and phosphoproteomic profiling of mouse and human neurodevelopmental models of mTORC1 activation and inhibition. These resources have the potential to unlock a new phase in mTORC1 neurodevelopmental biology, mTORopathy discovery and translational research.

To aid researchers in interrogating our proteomic and phosphoproteomics data we developed the PhosphoTargets platform. Data from each of the mouse and human NPC analyses can be viewed using interactive volcano plots or tables and ranked according to log fold change and adjusted p value or p value. Individual or multiple proteins of interest can be searched and the mouse and human NPC data can be viewed simultaneously to determine conservation. Also included are the proteomic and phosphoproteomic data from our recent study of TSC patient tuber and SEGA tissue (*49*). In order to compare our findings with previously published, non-neuronal models we have included data from rapamycin or Torin 1 treated HEK293E cells and Tsc2 mutant MEFs (*24*), as well as from rapamycin treated mouse ependymal glial cells and Tsc1/S6k mutant mouse kidney cells (*35, 50*). This platform will help to unlock the potential of our proteomic and phosphoproteomic resources for the community to elucidate new mTORC1-regulated molecular and cellular processes and identify novel biomarkers and therapeutic targets of mTORC1 signalling in NDDs.

mTORC1 signalling is required for all major cortical neurodevelopmental cellular processes including neuroepithelial and radial glial cell (RGC) symmetric division, asymmetric division of RGCs, migration of intermediate progenitor cells and neuroblasts, as well as maturation of neuroblasts to fully differentiated neurons (*10*). We selected specific timepoints in mouse and human neurodevelopmental models to capture mTORC1 targets in a range of progenitor and mature cell types in the developing brain. Mouse E16.5 is a period of intense neurogenesis and cortical expansion, equivalent to gestational weeks 12-14 in humans (*66*). Human NPCs are highly proliferative and express key developmental transcription factors, including PAX6, the neural stem/progenitor marker Nestin, and TUBB3, essential for axonal growth. Our NPC dataset may be particularly suitable for identifying novel mTORC1 targets that regulate NPC proliferation, as we demonstrated for USP9X.

We also identified novel mTORC1 targets that have spatiotemporal specific expression including the transcription factor BCL11B (also known as CTIP2), which is expressed in GABAergic cortical layer 5 interneurons, and ZEB2, which is expressed in interneurons in the medial ganglionic eminence (*67, 68*). These cell type specific mTORC1 targets cannot be identified in non-neurodevelopmental models and further demonstrate the potential of this resource in the context of the developing nervous system.

Advantages to our approach were the use of both proteomics and phosphoproteomics, the increased depth of coverage of phosphosites identified by using trypsin and chymotrypsin protease digests in parallel and the robust criteria for defining mTORC1 targets. The majority of phosphosites identified in the parallel trypsin/chymotrypsin LC-MS/MS experiments were mutually exclusive, greatly expanding the phosphoproteomic coverage compared to a single protease. Further evidence of this deep coverage are the novel mTORC1-dependent phosphosites we identified in well studied mTORC1 substrates, such as S6K1 and indirect targets such as RPS6. Moreover, we defined high confidence mTORC1 targets as phosphosites that were significantly increased by mTORC1 activation *and* significantly decreased by mTORC1 inhibition. This does not mean however that phosphosites that only respond to either mTORC1 activation or inhibition are not valid, especially when considering proteins or phosphosites that are conserved between mouse and human. For example, using our strict criteria phosphorylation of the zinc finger transcriptional repressor ZEB2, which has multiple functions in neurodevelopment and is mutated in Mowat–Wilson syndrome (*69*), at S784 was identified as an mTORC1 target in the human TSC NPC model (Figure 2C, Figure 4H, Supplementary table S21) but not in the TSC mouse model. However, ZEB2 phosphorylation was significantly increased at S760 in Tsc1 cKO versus control mice and significantly decreased at S754, S756 and S758 in Tsc1 cKO rapamycin treated mice (Supplementary tables S2, S4, S6, S8). Therefore, the criteria for considering whether any given protein is a potential mTORC1 target should be carefully considered and can be expanded or refined beyond the definition used here, which is easy to do using PhosphoTargets.

Our approach is limited by the use of whole brain tissue and a mixed population of cell types in the NPCs. Thus, proteins expressed only in rare cell types may not be detected. Moreover, ubiquitously expressed but low-abundance proteins might not be detected by data-dependent acquisition mass spectrometry, which is intensity-biased. In future, analysis of specific brain regions, or isolation of individual NPC cell types, could be used to focus on mTORC1 targets in specific cell types. In addition, data-independent acquisition proteomic methods might further expand the list of detected proteins and phosphosites since they are not biased towards more abundant proteins.

We identified a novel network of mTORC1 targets in mouse and human TSC models centred around RNA metabolism and mRNA splicing. Until recently, mis-regulation of mRNA splicing had not been observed in TSC cell or animal models or in TSC patients. Our recent analysis of TSC patient SEGA tissue showed increased phosphorylation of hnRNPs and spliceosome proteins as well as largescale changes in mRNA splicing (*49*). Regulation of splicing thus appears as a newly discovered function of mTORC1 in a human neurodevelopmental model of TSC and TSC patient brain tissue. Remarkably, in both SEGAs and NPCs, genes whose splicing is mis-regulated were enriched for genes encoding RNA binding proteins, as seen in other contexts with hnRNPs (*70, 71*). Moreover, the genes whose splicing are altered in SEGAs converge with cancer-associated splicing programmes and the genes we identified in the TSC human NPC model are strongly associated with perturbed neurodevelopment, suggesting that altered splicing contributes to the neurological manifestations in TSC.

Of the 142 mTORC1 target phosphosites identified in the human TSC model (Supplementary table S21), 37 are also significantly increased in SEGA tissue (*49*). As well as established mTORC1 substrates such as ACIN1 S240/S243 and LARP1 S766/S774, many of the phosphosites identified as mTORC1 targets in both human NPCs and SEGA tissue are in novel mTORC1 targets such as EDC3 S131, FOXK1 S441/S445, PCBP1 S264 and PNN S96/S100 (Supplementary table S21). These common phosphosites demonstrate that the targets of mTORC1 identified in human TSC1 KO NPCs are also relevant in the brain of TSC patients, and also the power of human disease and cell model phosphoproteomic resources to identify conserved phosphosites.

An exciting potential of the phosphoproteomic resource is to reveal the intersection between mTORC1 signalling and mTORopathies with other NDDs. The novel mTORC1 targets identified in both mouse and human models fall into several functional groups, including chromatin/transcriptional regulation, translational control, autophagy/protein turnover, cytoskeletal regulation, cell division and RNA metabolism (Figure 2C). These functions reflect the range of molecular processes necessary for neurogenesis and greatly expand knowledge of the repertoire of mTOR signalling in neurodevelopment. Variants in the genes encoding many of the novel mTORC1 targets we identified are associated with rare NDDs, suggesting a network of key disease associated neurogenic factors orchestrated by mTORC1 (Figure 4H). Most of these NDDs are caused by missense or heterozygous mutations where some functional protein remains. Targeting mTORC1 signalling to tune the activity or expression of these mTORC1 effectors may therefore have therapeutic potential in these rare NDDs.

## Materials and Methods

### Tsc1 cKO mice

Mouse studies were carried out in accordance with UK Home Office regulations and the UK Animals (Scientific Procedures) Act of 1986 (ASPA) under a UK Home Office licence (PPL 70/8719) and approved by the King’s College London Ethical Review Committee. All mice were housed in an animal facility of King’s College London on an alternating 12-hour light-dark cycle with free access to food and water.

*Nestin-Cre* mice (obtained from Jackson, stock number 003771) and *Tsc1^fl/fl^* mice (obtained from Jackson, stock number 005680) were maintained in a C57BL/6 background. Genotyping was performed on tails excised from embryos, following the recommended Jackson protocols for these stocks. Primers used for genotyping are shown in Supplementary table S24.

Brains were collected from E16.5 *Nestin-Cre;Tsc1^fl/+^*or *Nestin-Cre;Tsc1^fl/fl^* pups and dissected in half. Where rapamycin treatment was performed, pregnant dams were injected intraperitoneally with 1mg/kg rapamycin (Fluorochem Ltd) or an equivalent volume of vehicle (dimethyl sulfoxide (DMSO)) at E15.5, embryos were removed at E16.5, and embryonic brains were dissected. Hemispheres were flash frozen in liquid nitrogen and kept at -70°C before being lysed in urea buffer (8M urea, 50mM 4-(2-hydroxyethyl)-1-piperazineethanesulfonic acid (HEPES) pH 8.2, 10mM glycerol-2-phosphate, 50mM sodium fluoride, 5mM sodium pyrophosphate, 1mM ethylenediaminetetraacetic acid (EDTA), 1mM ethylene glycol-bis(β-aminoethyl ether) (EGTA), 1mM sodium vanadate, 1mM dithiothreitol (DTT), 1X cOmplete EDTA-free protease inhibitor cocktail (11836170001; Roche), 1X phosphatase inhibitor cocktail 3 (P044; Sigma-Aldrich), 500nM okadaic acid). Lysates were then kept at -80°C. For western blot, lysates were supplemented with 2X Pierce lithium dodecyl sulfate (LDS) Sample Buffer (NP008; ThermoFisher Scientific) containing 0.2M DTT and heated at 70°C for 10mins for denaturation before being stored at -20°C.

### Mass spectrometry

LC-MS/MS was used to analyse the total proteome and the phosphoproteome of mouse brain hemispheres and NPCs. After lysis in urea buffer, protein concentrations were determined using bicinchoninic acid (BCA) assay (Pierce™ BCA Protein Assay Kit, 23227; ThermoFisher Scientific), following the manufacturer’s instructions, and adjusted using urea buffer. 200µg protein per sample were used for embryonic mouse brains, and 50µg protein per sample were used for NPCs. Sample preparation, data acquisition and mass spectrometry analysis were carried out as previously described in (*72*) with some modifications.

### Protein reduction, alkylation & digestion

To break disulfide bonds, proteins were treated with 10mM DTT and incubated at 56°C for 1h. Afterward, samples were alkylated by adding 20mM iodoacetamide and kept in the dark at room temperature (RT) for 30mins. The reaction was quenched with an additional 20mM DTT, and the solution was diluted with 50mM HEPES (pH 8.5) to lower the urea concentration to less than 2M for trypsin digest and less than 1M for chymotrypsin digest. Proteins were then digested either overnight at 37°C with trypsin (1:20 enzyme:protein per sample; MS grade, 90058, ThermoFisher Scientific), or overnight at 25°C with chymotrypsin (1:20 enzyme:protein, Sequencing grade, V1061, Promega) with the presence of 10mM CaCl_2_.

### Sample cleanup and TMT labelling

After digestion, peptides were acidified with 1% trifluoroacetic acid (TFA) and purified using C18 columns (Nest Group BioPureSPN MACRO; Proto 300 C18; Part# HMM S18V). The samples were then freeze-dried in a vacuum concentrator. Peptides were then labelled with TMT 10-plex Isobaric Label Reagent Set (0.8mg per tag, 90110, ThermoFisher Scientific; LOT# UK288606 for mouse samples and WD313468 for NPC samples) according to the manufacturer’s instructions. Labelling efficiency and mixing accuracy were checked using an Orbitrap Eclipse Tribrid mass spectrometer with a 60-minute high-energy collision dissociation (HCD) MS2 fragmentation method. Labelling efficiency was greater than 99%.

### Phosphopeptide enrichment and fractionation

Labelled peptides were quenched with hydroxylamine, pooled together, and partially freeze-dried. They were then acidified to pH 2.0, cleaned up using C18 Sep Pak 1cc Vac columns (100mg bed volume; Waters), and dried. A final mixing check was performed using LC-MS/MS with a 240-minute HCD MS2 fragmentation method. The peptide mixture was then subjected to high-select SMOAC to capture phosphopeptides. It was first passed through a high-select TiO2 phospho-enrichment column (A32993; ThermoFisher Scientific), following the manufacturer’s protocol. Flow-through and wash fractions were combined, dried and subsequently used for Fe-NTA phospho-enrichment (A32992; ThermoFisher Scientific). One-tenth of the combined flow-through and wash fractions from this enrichment was used for total proteome analysis. The eluates from SMOAC were freeze-dried, solubilised and pooled together. Total proteome and phosphoproteome samples (except for NPCs) were subjected to high-pH reversed-phase fractionation (84868; ThermoFisher Scientific), dried and resolubilised in 0.1% trifluoroacetic acid, prior to LC-MS/MS.

### Data acquisition, processing and analysis

Data were acquired in data-dependent acquisition mode, with 180-minute HCD MS2 and 180-minute multistage activation synchronous precursor selection MS3 fragmentation methods (*73*), using an Orbitrap Eclipse Tribrid mass spectrometer (ThermoFisher Scientific). Acquired raw LC-MS/MS data from both MS2 and MS3 were co-processed using MaxQuant versions 1.6.2.10 (mouse) and 2.7.5.0 (NPCs) against either the expanded UniProt mouse (UP000000589, March 2021) or human (UP000005640, March 2023) reference proteome. For phosphoproteome, phosphorylation at STY was set as variable modification. Minimum peptide length of 6 and FDR 0.01 were used for the processing. Processed data were then analysed using an R-coding script tailored to isobaric labelling mass spectrometry. The script was generated as a hybrid, using the backbone and differential gene expression analysis of the ProteoViz package (*74*) as a general script workflow, and borrowing the normalisation script from the Proteus package (*75*). Briefly, the “proteinGroups.txt” and “Phospho (STY)Sites.txt” tables were read into matrices and filtered for “reverse” hits, “potential contaminant” and proteins “only identified by site.” Data were then normalised using CONSTANd (*76*), log2-transformed and differentially analysed using linear models for microarray data (limma). For all proteomic analyses, significance was defined as proteins with an adjusted p (padj) <0.05 and log2 fold change (log2FC) >0.34 or <-0.34. For all phosphoproteomic analyses, significance was defined as phosphopeptides with p<0.005 and log2FC >0.34 or <-0.34 as these criteria were optimal for inclusion of known mTORC1-regulated phosphosites across all 8 experiments. Volcano plots were generated using Graphpad Prism or ggrepel (a ggplot2 extension) as part of the tidyverse in R.

### Disorder score analysis

To calculate disorder score AIUPred v2.1 (*77*) from the github repository doszilab/AIUPred (May 2026) was used. Briefly, after local installation of AIUPred, standard disorder analyses were first executed with the command line “aiupred -i NAME.fasta -o results.tsv” where NAME.fasta were the same FASTA files used for the peptide matching in the MaxQuant processing of raw mass spectrometry data. The output file contains the disorder score for each amino acid in each protein entry. Next, the flanking sequence column from the MaxQuant output present in the limma analysis tables consisting of the phosphorylated residue and the 7 amino acids on each side was matched to the AIUPred output for the respective proteins using R/RStudio. Scores above 0.5 were considered intrinsically disordered regions (IDRs). Overall flanking sequence disorder score was calculated as the proportion of IDR amino acids in the flanking sequence.

### Reactome and gene ontology (GO) analyses

Reactome and GO analyses were performed using the STRING database V12.0 (*78*). For each experiment, all detected proteins or proteins corresponding to all detected phosphopeptides were used as the statistical background.

### Site-directed mutagenesis and cloning

Site-directed mutagenesis was used to generate serine-to-alanine phosphomutant constructs. Each polymerase chain reaction (PCR) contained 10µL 2X Platinum SuperFi II Master Mix (Invitrogen), 1µL forward primer (10µM stock), 1µL reverse primer (10µM stock), 1µL DNA template (10ng/µL stock) and ddH2O to a total volume of 20µL. Phosphorylation, intramolecular ligation/circularization, and template DNA removal was performed using a kinase, ligase and DpnI (KLD) enzyme mix (M0554, NEB) following the manufacturer’s protocol. DNA was transformed into Stable Competent *E. coli* (NEB), and colonies were grown at 30°C with kanamycin (15µg/mL). Plasmid purification was performed using either a QIAprep Spin Miniprep Kit (27106; QIAGEN), using the manufacturer’s instruction for a centrifuge-based method, or a ZymoPUREII™ Plasmid Maxiprep kit (D4203-1; Zymo Research), using the manufacturer’s instructions for a vacuum-based method. The mutations were confirmed using Sanger sequencing. Primers and plasmid details are shown in Supplementary table S24.

Gateway cloning was performed to generate N-terminal 6xHis/GST-tagged USP9X (aa2245-end) constructs suitable for use in bacterial protein purification. Primers were designed to insert attB fragments flanking the PCR product of interest. Each PCR contained 50µL 2X Platinum SuperFi II Master Mix (Invitrogen), 5µL forward primer (10µM stock), 5µL reverse primer (10µM stock), 5µL DNA template (10ng/µL stock) and ddH_2_O to a total volume of 100µL. The PCR programme used was the same as the one described above. The attB-flanked PCR product was then transferred into the expression clone carrying attP sites using a Gateway™ BP Clonase™ II Enzyme Mix kit (Invitrogen). Each BP reaction contained 1-5µL eluted attB-flanked DNA (50-100ng), 1.3µL pDONR^TM^207 (150ng/µL), 3µL 5X BP Clonase^™^ reaction buffer, and Tris-EDTA (TE) buffer (10mM Tris-HCl, 1mM EDTA pH 8) up to 15µL. Samples were vortexed and incubated overnight at 25°C. 5µL of BP reaction was used to assess reaction efficiency: 0.5µL proteinase K were added and the mix was incubated 10mins at 37°C. 1µL DNA was then transformed into 50µL 5-alpha competent *E. coli* (NEB) and grown at 37°C with gentamycin (10µg/mL). 3-5 colonies were sent for sequencing to confirm reaction efficiency. The remaining volume of BP reaction was combined with a destination vector carrying attR sites using a Gateway^™^ LR Clonase^™^ II Plus Enzyme Mix kit (Invitrogen). Each LR reaction contained: 10µL BP reaction, 2µL pDest-565 (150ng/µL) and 3µL LR Clonase^™^ Plus Enzyme Mix. Samples were vortexed and incubated at 25°C for several hours. 2µL of proteinase K were added, followed by 10mins of incubation at 37°C. 1µL DNA was transformed into 50µL 5-alpha competent *E. coli* (NEB) and grown at 37°C with ampicillin (100µg/mL). Constructs were confirmed using whole plasmid sequencing.

### HEK293T cell culture and transfection

HEK293T cells were kept in T75 flasks in Dulbecco’s Modified Eagle’s Medium/Nutrient Mixture F12 Ham (DMEM/F12) (D8437; Sigma Aldrich), with 10% foetal bovine serum (FBS) and 1% penicillin/streptomycin. Cells were passaged every 3-4 days in Trypsin-EDTA (25200056; Gibco) until they reached confluency. Passaging was done using either ratio splitting (1:10) or cell counting (1 million cells (1mL) in 14mL fresh medium) with a Countess™ Automated Cell Counter (Invitrogen). For plating on 12- and 24-well NUNC™ plates (ThermoFisher Scientific), confluent cells from a T75 flask were counted and plated at 100,000 (100µL) cells per well in 900µL fresh medium or at 50,000 (50µL) cells per well in 450µL fresh medium, respectively. For plating on 6-well NUNC™ plates (ThermoFisher Scientific), confluent cells were resuspended in 10mL of medium and 200µL was added to each well in 1.8mL fresh medium.

Transfections were carried out 24hrs after plating onto 6-, 12- or 24-well plates using X-tremeGENE™ 9 DNA Transfection reagent (6365809001; Roche) and Opti-MEM (31985062; Gibco), according to the manufacturer’s instructions.

48hrs after transfection (or 72hrs after plating), cells were treated with either 100nM rapamycin (SM83; Cell Guidance Systems), 100nM Torin1 (4247/10; Biotechne), or an equivalent volume of vehicle (DMSO) for 1h. Treatment was immediately followed by cell lysis.

### Generation of USP9X phosphomutant hiPSCs

Two point-mutations were introduced into the *USP9X* in KOLF2.1J hiPSCs (*79*). The transcript of interest for USP9X corresponds to ENST00000378308.7 (USP9X-202), chromosome X, based on Genome Reference Consortium Human Build 38 (GRCh38/hg38). The first point mutation was c.7327T>G giving rise to p.Ser2443Ala, and the second was c.7639T>G giving rise to pSer2547Ala. The optimal design for both target sites used two Cas12a single guide RNAs (sgRNAs): sgRNA_114 and sgRNA_115. The presence of the desired point mutations within the sgRNA sequence was sufficient to prevent re-engagement and cutting after editing, eliminating the need for silent mutations. Silent mutations (c.7329T>G for p.Ser2443Ala, c.7638A>G and c.7641C>G for pSer2547Ala) were introduced to generate a Narl restriction enzyme (RE) site for screening of clones containing the desired point mutations. The hiPSCs were simultaneously nucleofected with both sgRNAs and single-stranded oligodeoxynucleotides (ssODNs).

sgRNA/Cas12a mixes were made as follows: 2.5µL of sgRNA_114 (100µM stock in IDTE (10mM Tris, 0.1mM EDTA)) or sgRNA_115 (100µM stock in IDTE), 5µL Opti-MEM, 2µL polyglutamic acid (100mg/mL), and 2µL hifiCas12a (10µg/mL stock). They were mixed up and down 5 times and incubated 15mins at 37°C in a beads bath. 2.5µL of USP9X_2443_ssODN (100µM stock with sgRNA_114) or USP9X_2547_ssODN (100µM stock with sgRNA_115) and 1µL of Cas12a electroporation enhancer (IDT) were then added, and the solutions mixed up and down.

KOLF2.1J hiPSCs cultured on 6-well NUNC™ plates (ThermoFisher Scientific) pre-treated with Rho-associated coiled-coil kinase (ROCK) inhibitor (ROCKi) were rinsed with pre-warmed EDTA-based Versene Solution (15040033; Gibco) and incubated with Accutase for 2mins. Accutase was removed whilst the cells were still adherent, and cells were triturated with StemFlex™ medium (A3349401; Gibco) supplemented with 1X RevitaCell™ Supplement (Gibco; A2644501). Cells were pelleted by centrifugation at 1000rpm for 4mins. Pellets were washed with ∼40µL of Primary Cell P3 buffer (Lonza). Excess medium was removed, and pellets were resuspended in 85µL P3 buffer. All 85µL of cell suspension were added to the tubes containing sgRNA/Cas12a and mixed up and down 5 times before being transferred to a cuvette. Nucleofection was performed using program CA-137 on an Amaxa 4D nucleofector (Lonza) using a 96-well format kit (V4XP-3032; Lonza). StemFlex™ medium supplemented with homology-directed repair (HDR) Enhancer v2 (IDT) was immediately added to the tubes, and the entire contents were transferred onto new 6-well plates. Medium was changed at the end of the day to StemFlex™ with 1X RevitaCell™ and HDR Enhancer v2. The next day, medium was changed to StemFlex™ only. The following day, some of the cells were plated at clonal density and the remainder of the bulk nucleofected cells were frozen. Clones were picked after ∼12 days of expansion.

48 clones were initially picked, expanded and screened. Screening was performed by PCR of the target site with the primers shown in Supplementary table S24, followed by Sanger sequencing. The double S2443A/S2547A point mutation clones 11 and 14 were further expanded and cryopreserved. KaryoStat+ analysis (Fisher Scientific) confirmed that the karyotype was normal in control and USP9X mutant hiPSC lines and the absence of off-target mutations in *USP9Y* was confirmed by Sanger sequencing. Short tandem repeat (STR) profiling was used to confirm KOLF2.1J cell line identity. PluriTest (ThermoFisher) and immunostaining were used to confirm pluripotency of control and USP9X mutant hiPSCs.

### Generation of TSC1 KO hiPSCs

To generate TSC1 KO hiPSCs, the Cas9 IBJ4 cell line was used, originally from Lieber Institute for Brain Development, MD, USA. The Cas9 cell line carries the Cas9 in a AAVS1 locus under a tetracycline inducible promoter. The cells were cultured as feeder-free cultures in Essential 8*TM* Medium (Thermo Fisher, #A1517001) and on Matrigel (Corning, #354234). For the nuclease transfection of the stem cells, the Lipofectamine CRISPRMAX Cas9 Transfection kit (Thermo Fisher, CMAX00008) was used, following the manufacturer’s protocol. The TSC1 RNA guides for the transfection originate from the Bateup Group (*41*): Guide 1: ACTATGTGTCTCCCCCGTGA; Guide 2: AGACATGAATGGTGACACCT. The cells were treated for 48 h altogether with doxycycline (when cells reached ∼70% confluency), starting with 24 h prior the transfection for the induction of Cas 9 expression in the cells. The Lipofection took place one day after passaging the stem cells, leading to the Lipofection of many smaller colonies and therefore an increased surface of cells. The transfection was repeated the next day to increase the success rate of the lipofection. The transfected cells were subcloned and DNA extracted for sequencing. For genome sequencing, the Oxford Nanopore MinION sequencing platform was used, which required an additional M13 tag attached to the PCR primers, to ensure a binding spot for the sequencing barcodes to ensure the possibility to differentiate between the subclones (Supplementary table S24). Barcoded PCR products were pooled together and purified using the Wizard® SV Gel and PCR Clean-Up System (Promega, #A9281). 1μg of purified DNA was used for sequencing using the 1D2 sequencing kit (Oxford Nanopore, #SQK-LSK308). The MinION sequencing was run with the MinKNOW programme until around 400,000 reads were acquired. A custom written script was used to identify if deletions were present in the barcoded samples. To confirm the presence of deletions, the barcoded reads were checked on Integrative Genomics Viewer. The TSC1 KO iPSC cell line contains a 511bp deletion in Exon17 of the *TSC1* gene. STR profiling was used to confirm IBJ4 cell line identity. PluriTest (ThermoFisher) and immunostaining were used to confirm pluripotency of control and TSC1 KO hiPSCs.

### hiPSC culture

hiPSCs were incubated in normoxic conditions (5% CO_2_, 20% O_2_) at 37°C and maintained in StemFlex™ medium on 6-well NUNC™ plates (ThermoFisher Scientific) coated with Geltrex™ (A1413302; Gibco) diluted 1:100 in Knockout™ DMEM (10829018; Gibco). Cells were passaged at a 1:6 ratio upon reaching 60-70% confluency. During passage, cells were rinsed with and incubated in EDTA-based Versene Solution (15040066; Gibco) for 5mins, then replated in new GeltrexTM-coated plates.

### Directed differentiation of hiPSCs into NPCs

The hiPSC lines were differentiated into forebrain cortical NPCs by dual SMAD inhibition following a protocol adapted from (*80*). Briefly, hiPSCs were passaged onto GeltrexTM-coated 6-well NUNC™ plates at a 2:3 ratio and maintained under normoxic conditions for 48hrs until they approached ∼80% confluence. For USP9X cell lines, another passage was performed at a 2:1 ratio, resulting in cells reaching 100% confluence the following day. For TSC1 KO cell lines, cells were grown a further 24hrs until they reached 90-95% confluence.

Directed differentiation was then initiated by changing StemFlex™ medium to neuralisation medium containing N2:B27 (N2 medium and B27 medium at a 1:1 ratio) supplemented with 100nM LDN193189 (SML0559; Sigma-Aldrich) and 10µM SB431542 (S4317; Sigma Aldrich) for dual SMAD inhibition. N2 medium consisted of DMEM/F12 (D8437; Sigma Aldrich), supplemented with 1X GlutaMAX™ Supplement (35050061; Gibco) and 1X N-2 Supplement (15410294; Gibco). B27 medium consisted of Neurobasal™ medium (21103049; Gibco), 1X GlutaMAX™ Supplement and 1X B-27™ Supplement Minus Vitamin A (12587001; Gibco). The neuralised cells were incubated under normoxic conditions at 37°C. Neuralisation medium was replenished every 24hrs from day 0 to day 7. At the end of the neuralisation period, on day 8, neuralisation medium was replaced with N2:B27 without inhibitors, which was replenished daily until day 12. On day 12, N2:B27 was replaced with N2:B27 containing 200µM L-ascorbic acid 2-phosphate (AA2P) (A8960; Sigma-Aldrich), which was replenished daily from this day onwards.

The cells underwent four neural passages (NPs): on day 7 (NP1, 1:1 ratio), day 12 (NP2, 1:1 ratio), day 15 (NP3, 1:1 ratio) and day 18 (NP4, 1:2 ratio). NP1 and NP2 were performed, briefly, as follows: cells were washed with RT Hanks’ Balanced Salt Solution (HBSS) without calcium and magnesium (14170088; Gibco) and incubated for 5mins at 37°C with StemPro™ Accutase™ Cell Dissociation Reagent (A1110501; Gibco). The cells were then collected with the Accutase by pipetting a maximum of 5 times with a p1000 micropipette to avoid breaking down cell clusters. The cells were mixed with RT DMEM/F12 at a 1:4 ratio and centrifuged at 200g for 2mins to separate the cells and Accutase. Cells were gently (to avoid breaking down cell clusters) resuspended in the appropriate medium (neuralisation medium during NP1, N2:B27 with AA2P during NP2) supplemented with 10µM ROCKi (SCM075; Sigma-Aldrich) to enhance cell survival. Cells were then plated on new GeltrexTM-coated 6-well plates at a 1:1 ratio. ROCKi was removed 18-24hrs after each passage and replaced with appropriate medium (N2:B27 without inhibitors after the first passage, N2:B27 with AA2P after the second passage). The same process was followed for NP3 and NP4, with a few modifications. Collection of cells with Accutase was done by pipetting 20-30 times with a p1000 micropipette to obtain single cells. Similarly, resuspension in medium (N2:B27 with AA2P) was performed vigorously to obtain single cells. Finally, final plating took place during NP4 and required different types of plates and splitting ratios (described below) depending on upcoming experiments.

Successful differentiation of hiPSCs into NPCs was validated by immunostaining for the pluripotency marker OCT4 and the NPC markers paired box protein Pax-6 (PAX6), βIII-tubulin (TUBB3) and Nestin, as described in (*80*) and (*81*). A biological replicate was defined as a culture that has been passaged independently for at least 3 passages prior to sample fixation or collection.

For rapamycin treatment, two days after NP4 cells were treated with 100nM rapamycin (SM83; Cell Guidance Systems) or an equivalent volume of vehicle (DMSO) for 5hrs. Treatment was delivered via half medium change and interrupted by either cell fixation or lysis.

For 5-ethynyl-2’-deoxyuridine (EdU) labelling, two days after NP4 cells were labelled with EdU using a Click-iT™ EdU Cell Proliferation Kit for Imaging (C10337; Invitrogen) following the manufacturer’s instructions. EdU labelling solution was delivered via half medium change and cells were incubated in their usual growth conditions 4hrs before cell fixation.

### RNA extraction

During NP4, NPCs were seeded 1:1 to a new GeltrexTM-coated 6-well NUNC™ plate. To extract RNA, on DIV21, cells were lysed with 1mL TRIzol™ Reagent (15596026; ThermoFisher Scientific) per each well of a 6-well plate for 5 minutes at room temperature, pipetting vigorously to ensure complete lysis. Then, 200 µL of chloroform was added to each tube and incubated for 3 minutes at room temperature, prior to spinning at 10.000 RPM for 5 minutes at 4°C. The aqueous phase was then transferred to another tube, and 500 µL of isopropanol was added to each tube, along with 1µL of RNA Grade Glycogen (R0551; ThermoFisher Scientific) to then incubate for 15 minutes at room temperature prior to spinning at 13.000 RPM for 15 minutes at 4°C. After discarding the supernatant, RNA pellet was washed with 1mL of 80% ethanol, spinning at 13.000 RPM for 5 minutes at 4°C. After the wash, pellet was allowed to air dry for around 20 minutes prior resuspension in RNA-free water. To enhance RNA quality, we then performed sodium-acetate precipitation, adding 0.1 volumes of sodium-acetate and 3 volumes of 100% ethanol and placed it at -80°C overnight. The day after, RNA has been spun down at 13.000 RPM for 15 minutes at 4°C. Supernatant was discarded and the RNA pellet was washed twice with 1mL of 80% ethanol spinning 13.000 RPM for 5 minutes at 4°C. RNA pellet was then resuspended in RNA-free water and measured using NanoDrop™ 2000.

### RNA Sequencing

Total RNA was extracted from n=4 independent differentiations of NPCs. Messenger RNA was purified from total RNA using poly-T oligo-attached magnetic beads. After fragmentation, the first strand cDNA was synthesized using random hexamer primers followed by the second strand cDNA synthesis. The library was ready after end repair, A-tailing, adapter ligation, size selection, amplification, and purification. The library was checked with Qubit and real-time PCR for quantification and bioanalyzer for size distribution detection. Sequencing was performed using Illumina NovaSeq X Plus Series with a PE150 paired-end strategy. Raw data of fastq format were firstly processed through fastp software. HISAT2 was used to align the paired-end clean reads to the reference genome hg38. featureCounts was used to count the reads numbers mapped to each gene and then the FPKM of each gene was calculated based on the length of the gene and reads count mapped to this gene.

### Splicing analysis

The .fastq files were processed using the nf-core framework nf-core/rnaseq v3.14.0 pipeline (*82*) with nextflow v25.10.4. Files have been aligned to GRCh38 for basecalling, and GENCODE v49 was used for annotation. For the alignment, the STAR method was performed, followed by Salmon quantification. Built-in QC was assessed at this stage. Differential expression analysis was performed using DESeq2 pipeline (*83*).

Splicing analysis was performed using nf-core/rnasplice v1.0.4 with rMATS (*82*) enabled to identify splicing events defined as SE (Skipped Exon), A3SS (Alternative 3’ Splice Site), A5SS (Alternative 5’ Splice Site), MXE (Mutually exclusive exons) and RI (Retained Intron). Events detected were then combined and filtered for FDR<0.05 and ΔPSI > 0.10. The genes resulting in significant splicing events were used for GO analysis.

### Proteogenomic analysis

Proteogenomic analysis was carried out using the github repository NKI-TGO/SPLICIFY (*63*) and python 2.7. Briefly, paired-end raw RNAseq files were trimmed removing low quality reads and Illumina adapters using trimmomatic v0.33 with the following parameters: phred = 33, illuminaclip = data/adapters/TruSeq3-PE.fa:2:30:10, leading = 20, trailing = 20, crop = 150, avgqual = 20, slidingwindow = 4:20, minlen = 150. Reads were aligned to the human genome using STAR aligner. Firstly, star_index was generated using the genome FASTA “GRCh38.primary_assembly.genome.fa” and the genome annotation “gencode.v50.primary_assembly.annotation.gtf” followed by the alignment step. Transcript splicing analysis was carried out using rMATS v3.2.5 in an unpaired mode. rMATS results were converted to fasta sequences using the matsToFasta step with the following parameters: comparisonName = group1vsgroup2, input = JCEC, # input = JunctionCountsOnly, fdr = 0.05, outputBed = PathToBED.bed, ifMXE = yes. The proteomic part of SPLICIFY uses the MaxQuant output tables “peptides.txt” and “evidence.txt” along with the matsToFasta outputs. This part is carried out in three steps starting with the “extraction of variant peptides from the evidence file” through “adding RNA splice variant information to the peptide table” to “running differential peptide expression” using *limma* and imputation for missing values.

### GO analysis

GO analysis was performed using the STRING database V12.0 (*78*). All detected proteins or proteins represented by phosphopeptides were used as the statistical background.

### Kinase assay

Plasmids containing desired inserts were transformed in BL21(DE3) competent *E. coli* (72035; Sigma-Aldrich). Colonies were then inoculated in 50mL terrific broth (TB) medium supplied with ampicillin (100µg/mL) and grown overnight shaking at 37°C. 750mL TB with antibiotic was inoculated with overnight culture to an optical density OD600 = 0.05. Cells were grown at 37°C shaking until reaching OD600 = 0.7 followed. Induction of protein expression was obtained via the addition of 300µL isopropyl β-D-1-thiogalactopyranoside (IPTG). Cells were then incubated shaking for 3hrs at 37°C.

Induced overnight bacterial culture was centrifuged at 4,000g for 15mins at 4°C. 0.5g cell pellet was then resuspended in 1mL lysis buffer for GST-tagged proteins (150mM NaCl, 50mM HEPES, 0.1mM EDTA, 0.5% Triton X-100, 10% glycerol, 1mM DTT, 1X cOmplete EDTA-free protease inhibitor cocktail, 2mM phenylmethanesulfonyl fluoride (PMSF)). Bacterial pellets were sonicated on ice 5×5s at 10s intervals. Lysates were put on a rotating wheel 10mins at 4°C before being centrifugated at 4°C, 21,000g. The supernatant was centrifuged 5mins at 4°C, 21,000g. The new supernatant was then incubated for 1h on a rotating wheel at 4°C with 50µL (slurry) beads (Glutathione Sepharose™ 4B, 17075601; Cytiva), which had been pre-washed in lysis buffer. Columns were washed once in lysis buffer, twice in high salt buffer (500mM NaCl, 50mM HEPES, 0.1mM EDTA, 1mM DTT, 1X cOmplete EDTA-free protease inhibitor cocktail, 0.2mM PMSF), twice in low salt buffer (150mM NaCl, 50mM HEPES, 0.1mM EDTA, 1mM DTT, 1X cOmplete EDTA-free protease inhibitor cocktail, 0.2mM PMSF), and eluted in 50µL elution buffer (150mM NaCl, 50mM HEPES pH 7.4-7.5, 40mM reduced L-Glutathione).

Purity of each protein fragment was assessed by Coomassie staining on NuPAGE 4-12% Bis-Tris polyacrylamide gels. Gels were rinsed in ddH_2_O, incubated in InstantBlue® Coomassie Protein Stain (ISB1L) (ab119211; Abcam) overnight at RT on a shaker, and rinsed 5×2mins in ddH2O. Gel imaging was performed via epi-illumination using an Amersham Imager 600 RGB or Amersham ImageQuant™ 800 GxP.

Kinase assays were conducted in kinase buffer (25mM HEPES pH 7.2, 50mM NaCl, 0.01% Tween® 20, 10mM MnCl2, 1mM EDTA) with a human recombinant truncated (aa1362-end) FLAG-tagged mTOR kinase (14-770; Sigma-Aldrich). Target proteins were purified 6xHis-GST-tagged truncated (aa2245-end) versions of USP9X (WT, S2443A, S2547A and Double). A GFP-tagged 4E-BP1 (gift from Professor Andrew Tee from Cardiff University) was used as a control target protein.

Kinase reactions contained 35µL kinase buffer, 30nM kinase, 400nM target protein, 10mM adenosine-5’-(3-thio)-triphosphate (ATPγS) and ddH_2_O up to 60µL. Reactions occurred over 30mins at 30°C in a shaking incubator. Half of the reaction volume was kept at -20°C for later use, while the other half was alkylated 1h at RT by addition of 20mM EDTA and 6mM p-Nitrobenzyl mesylate (PNBM) to allow the detection of thiophosphate esters (THP). In preparation for western blots, reactions were supplemented with 4X Pierce LDS Sample Buffer containing 0.4M DTT and heated at 70°C for 10mins for denaturation before being stored at -20°C.

### Western blot analysis

HEK293T cells were lysed directly in 2X Pierce LDS Sample Buffer containing 0.2M DTT, followed by sonication on ice 5×5s at 10s intervals and denaturation at 70°C for 10mins. Lysates were then stored at -20°C.

HiPSCs and NPCs were lysed in RIPA Lysis and Extraction Buffer (ThermoFisher), 1X cOmplete EDTA-free protease inhibitor cocktail (Roche), 1X phosphatase inhibitor cocktail (Sigma). Protein concentrations were estimated using BCA assay and adjusted. Lysates were then supplemented with 2X Pierce LDS Sample Buffer containing 0.2M DTT, heated at 70°C for 10mins for denaturation, and stored at -20°C.

Samples were run on NuPAGE 4-12% or 8% Bis-Tris polyacrylamide gels (ThermoFisher Scientific) and transferred to a PVDF membrane (Millipore) using wet transfer. Membranes were blocked in 10% non-fat milk in Tris-buffered saline with 0.1% Tween® 20 detergent (TBST) for 30mins. They were then incubated with primary antibodies in 5% non-fat milk or 5% BSA in TBST, either overnight at 4°C or for 3hrs at RT. Horseradish peroxidase (HRP)-conjugated secondary antibody incubation was carried out at RT for 1h. For western blot detection, the membranes were incubated with enhanced chemiluminescence (ECL) (Amersham ECL Prime Western Blotting Detection Reagent, RPN2236; Cytiva) and visualised with a chemiluminescence digital imaging system (Amersham Imager 600 RGB or Amersham ImageQuant™ 800 GxP). Image analysis was performed using the ImageQuant™ IQTL 11.0 software. Image contrast and brightness adjustments for publication in this thesis were performed using Fiji (*84*). For quantification, total protein levels were normalised against α-tubulin and phosphorylation levels were normalised to the corresponding total protein levels. Primary and secondary antibodies are shown in Supplementary table S25.

### Immunocytochemistry (ICC)

HiPSCs during NP4 cells were counted using a NucleoCounter® NC-200™ (Chemometec), and plated on Falcon® 96-well Black/Clear Flat Bottom TC-treated Imaging Microplates (353219; Corning) at a density of 80,000 cells per well. ROCKi-containing medium was replaced with N2:B27 with AA2P 18-24hrs later, and the cells were left to recover for another 24hrs before fixation.

HiPSCs or NPCs were plated on coverslips or 96-well plates were fixed with 4% paraformaldehyde (PFA) for 10mins at RT. They were then washed in Dulbecco’s phosphate-buffered saline (DPBS), followed by blocking 30mins at RT in blocking buffer (2-10% goat serum and 0.01% Triton X-100 in DPBS). Plates and coverslips were either used immediately for ICC or kept at 4°C in phosphoate-buffered saline (PBS) with 0.02% sodium azide (PBSA). Primary antibody incubation was performed overnight at 4°C in blocking buffer. The next day, cells were washed and incubated in secondary fluorescent antibodies in blocking buffer 1h at RT. Final washes were performed, during which cells were stained with 0.5µg/mL 4′,6-diamidino-2-phenylindole (DAPI) (D3571; Invitrogen). Plates were kept at 4°C in PBSA. For coverslips, mounting was done using Fluoromount-G™ (Invitrogen) and slides were stored in the dark at 4°C after drying. Primary and secondary antibodies are shown in Supplementary table S25.

### Image acquisition and analysis

Images of coverslips were acquired by confocal microscopy using a VT-iSIM (Olympus). Images of 96-well plates were acquired by confocal microscopy using an Operetta CLS™ High-Content Analysis System (PerkinElmer) and analysed using the Harmony™ High-Content Imaging and Analysis software. Image contrast and brightness adjustments were performed using Fiji (*84*).

## Statistical analysis

Statistical analysis was performed using the GraphPad Prism 9 software (Dotmatics), unless stated otherwise, with the statistical tests indicated in the figure legends. Statistical significance was observed as follows: ns, not significant; *, P<0.05; **, P<0.01; ***, P<0.001; and ****, P<0.0001.

## Data availability

Supplementary tables can be accessed by reviewers through Dryad using this link: http://datadryad.org/share/LINK_NOT_FOR_PUBLICATION/V-OB3NNIv9M-jEGr6XrEjuj-9pBktow1THfRAMWCT14

The mass spectrometry proteomics data have been deposited to the ProteomeXchange Consortium via the PRIDE partner repository (*85*) with the dataset identifier PXD081781. Reviewers can access the deposited files on the PRIDE website using the same accession identifier with the token: 29JF3XRvYm5v. RNA-seq data have been deposited in GEO, accession GSE336501, reviewer token ozcziuqctpkzrqn.

## Acknowledgements

We are grateful to Ian Tully for the genome analysis script and Nick Klopp for advice on the disorder analysis.

## Author contributions

Conceptualisation: SKU,

## JMB Proteomics: HF, MS, SRM

In vivo studies: PB

In vitro studies: SC, SRM, MG, LK, SG

Computational analysis: MW

Data analysis: SRM, MG, SG, JMB

Writing: JMB

Review and editing: All authors

Supervision: SKU, DS, AH, JMB

Funding acquisition: SKU, DS, AH, JMB

## Supplementary figures

**Supplementary figure S1.**
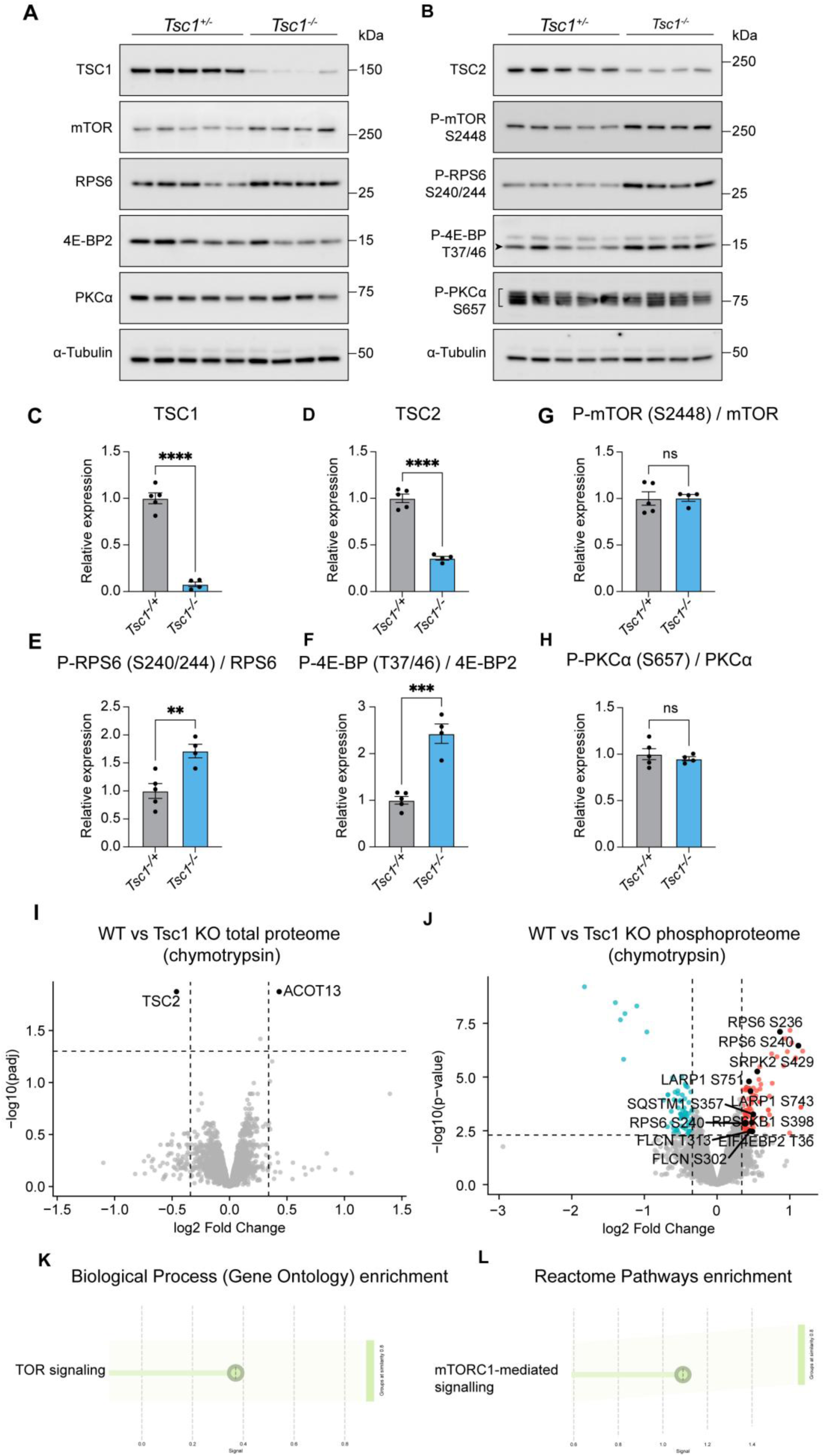
Validation of the Tsc1 cKO mouse model. (A, B) Western blot analysis of E16.5 brain tissue from the Tsc1 cKO versus control mice. (C, D) Quantification of TSC1 (C) and TSC2 (D) expression levels in E16.5 brain tissue from the Tsc1 cKO versus control mice. (E-H) Quantification of P-RPS6 S240/S244 (E), P-4E-BP T37/T46 (F), P-mTOR S2448 (G), P-PKCα S657 (H) levels in E16.5 brain tissue from the Tsc1 cKO versus control mice. (I, J) Volcano plots of the proteomic (I) and phosphoproteomic (J) data from E16.5 Tsc1 cKO mice versus control E16.5 brain tissue (chymotrypsin digest). Proteins or phosphopeptides with significantly increased expression coloured red and significantly decreased expression coloured blue. Established mTORC1 targets are labelled. (K, L) GO biological process (K) and Reactome (L) analysis of proteins corresponding to phosphopeptides with significantly increased in Tsc1 cKO versus control mice. Control n=5 brains, Tsc1 cKO n=4 brains. Data are represented as mean ± SEM and were analysed using the students t test. ns not significant, **p < 0.01, ***p<0.001, ****p < 0.0001.

**Supplementary figure S2.**
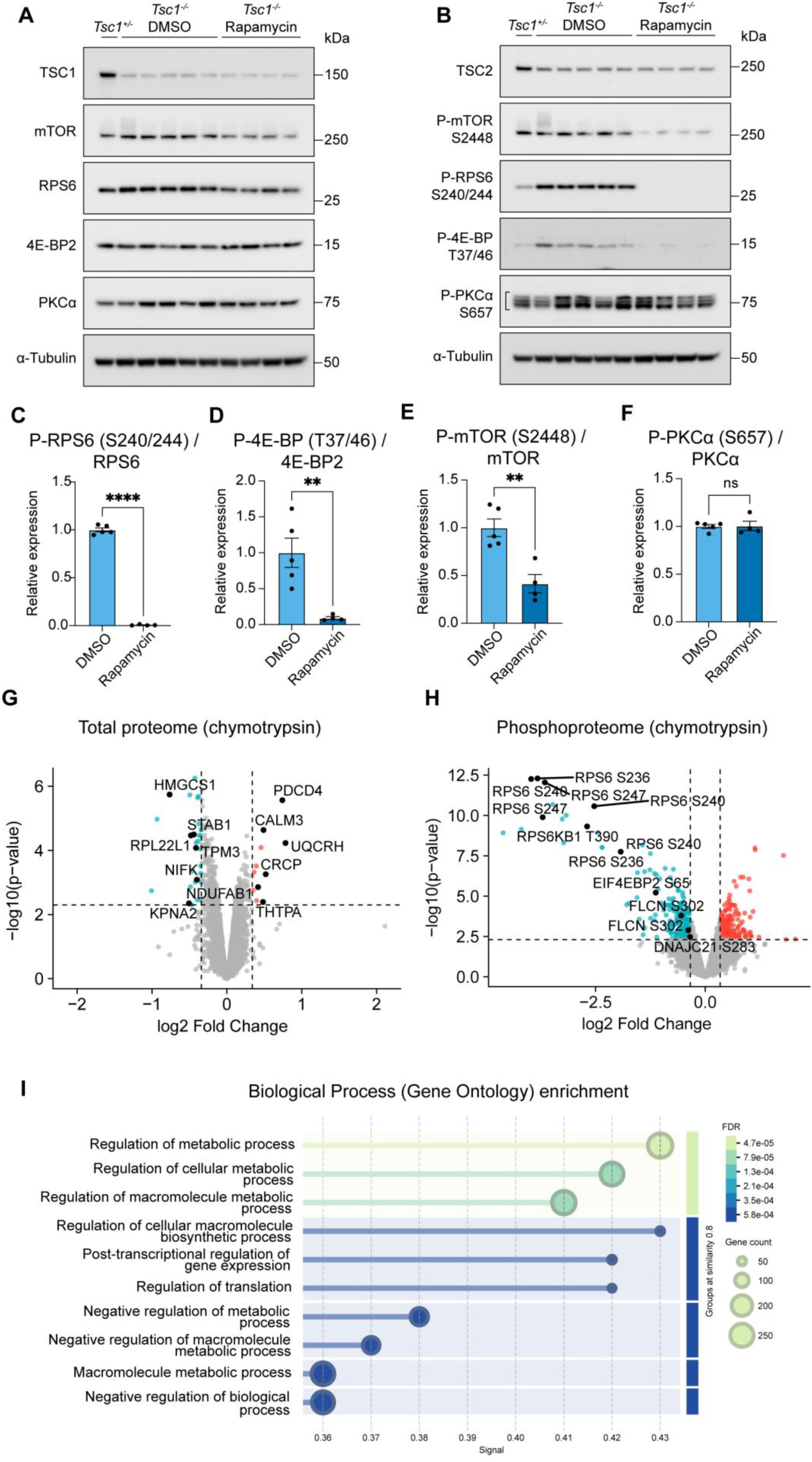
Validation of mTORC1 inhibition by rapamycin treatment in Tsc1 cKO mice. (A, B) Western blot analysis of E16.5 brain tissue from Tsc1 cKO vehicle treated or rapamycin treated mice. (C, D) Quantification of TSC1 (C) and TSC2 (D) expression levels in E16.5 brain tissue from Tsc1 cKO vehicle treated versus rapamycin treated mice. (E-H) Quantification of P-RPS6 S240/S244 (E), P-4E-BP T37/T46 (F), P-mTOR S2448 (G), P-PKCα S657 (H) levels in E16.5 brain tissue from Tsc1 cKO vehicle treated versus rapamycin treated mice. (I, J) Volcano plots of the proteomic (I) and phosphoproteomic (J) data from Tsc1 cKO vehicle treated versus rapamycin treated mice (chymotrypsin digest). Established mTORC1 targets are labelled in (J). Proteins or phosphopeptides with significantly increased expression coloured red and significantly decreased expression coloured blue. (K) GO biological process analysis of proteins corresponding to phosphopeptides with significantly decreased expression in Tsc1 cKO vehicle treated versus rapamycin treated mice. Vehicle treated Tsc1 cKO n=5 brains, rapamycin treated Tsc1 cKO n=5 brains. Data are represented as mean ± SEM and were analysed using the students t test. ns not significant, *p<0.05, **p < 0.01, ****p < 0.0001.

**Supplementary figure S3.**
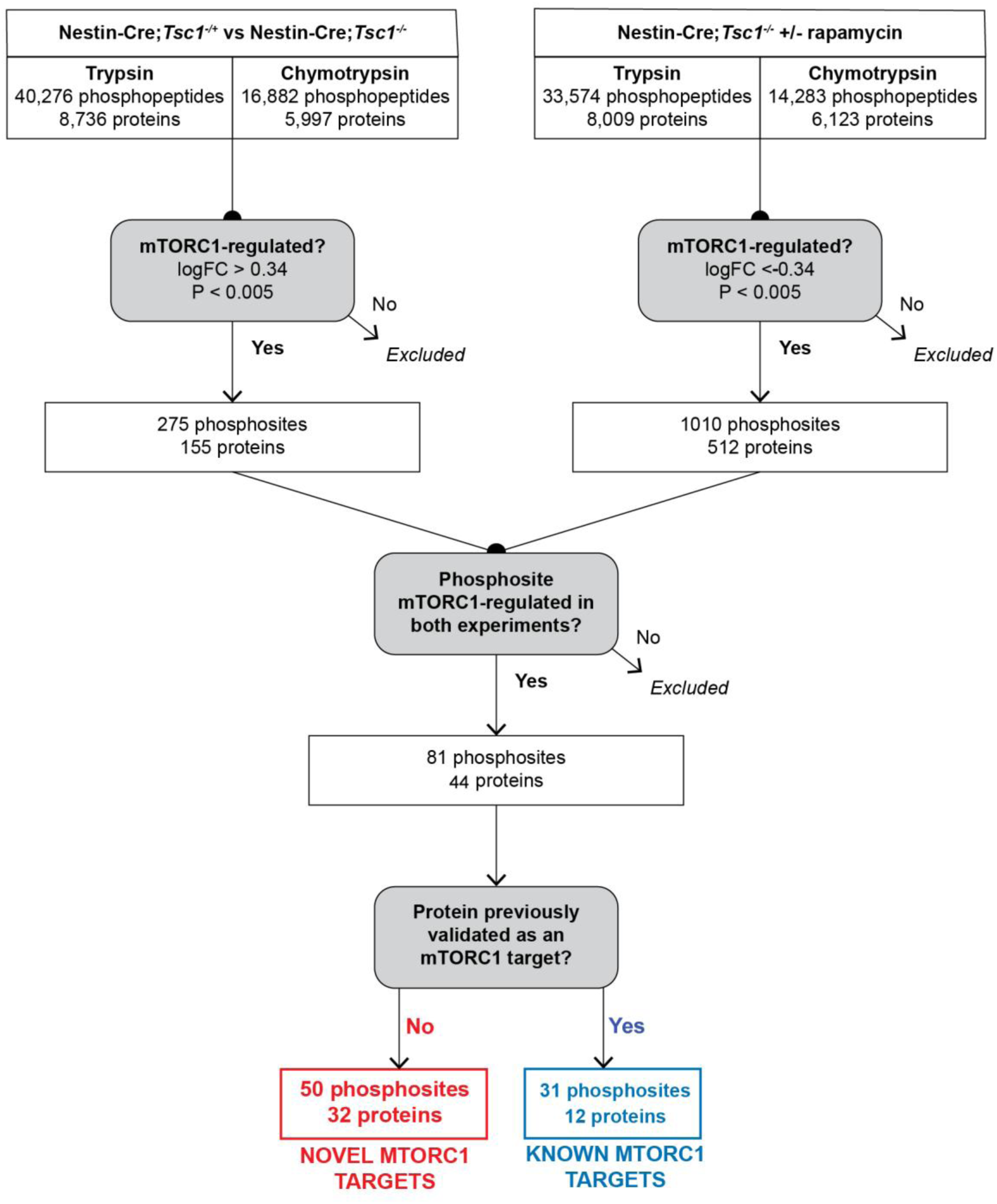
**Pipeline used to select novel mTORC1 target phosphosites in mouse neurodevelopment.**

**Supplementary figure S4.**
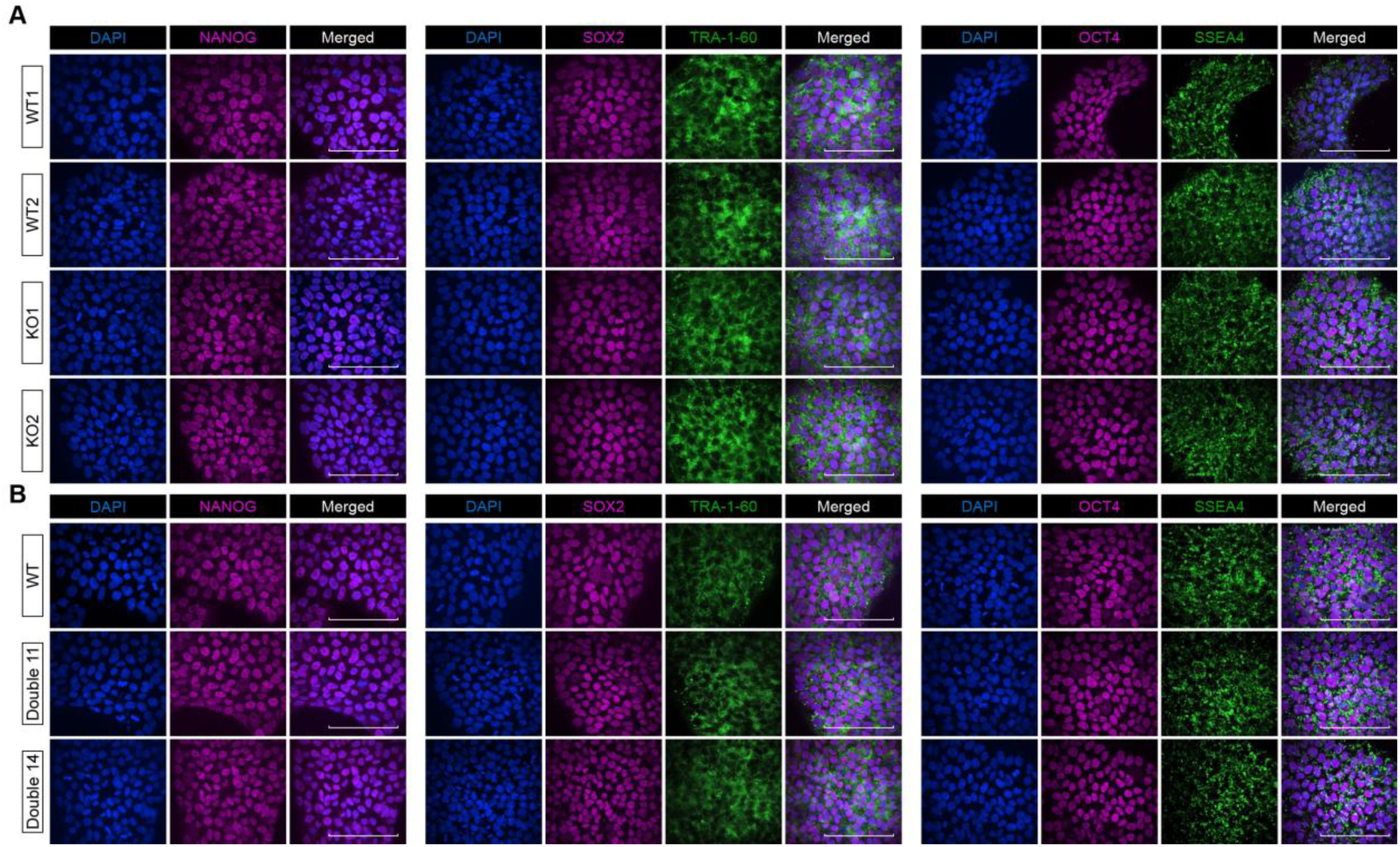
Validation of TSC1 KO and USP9X phosphomutant iPSCs. (A) Immunostaining for Nanog, SOX2, TRA-1-60, OCT4 and SSEA4 in control (WT1 and WT2) and TSC1 KO (KO1 and KO2) iPSC lines. (B) Immunostaining for Nanog, SOX2, TRA-1-60, OCT4 and SSEA4 in control (WT) and USP9X phosphomutant (double 11 and double 14) iPSC lines. Scale bar: 100µm.

**Supplementary figure S5.**
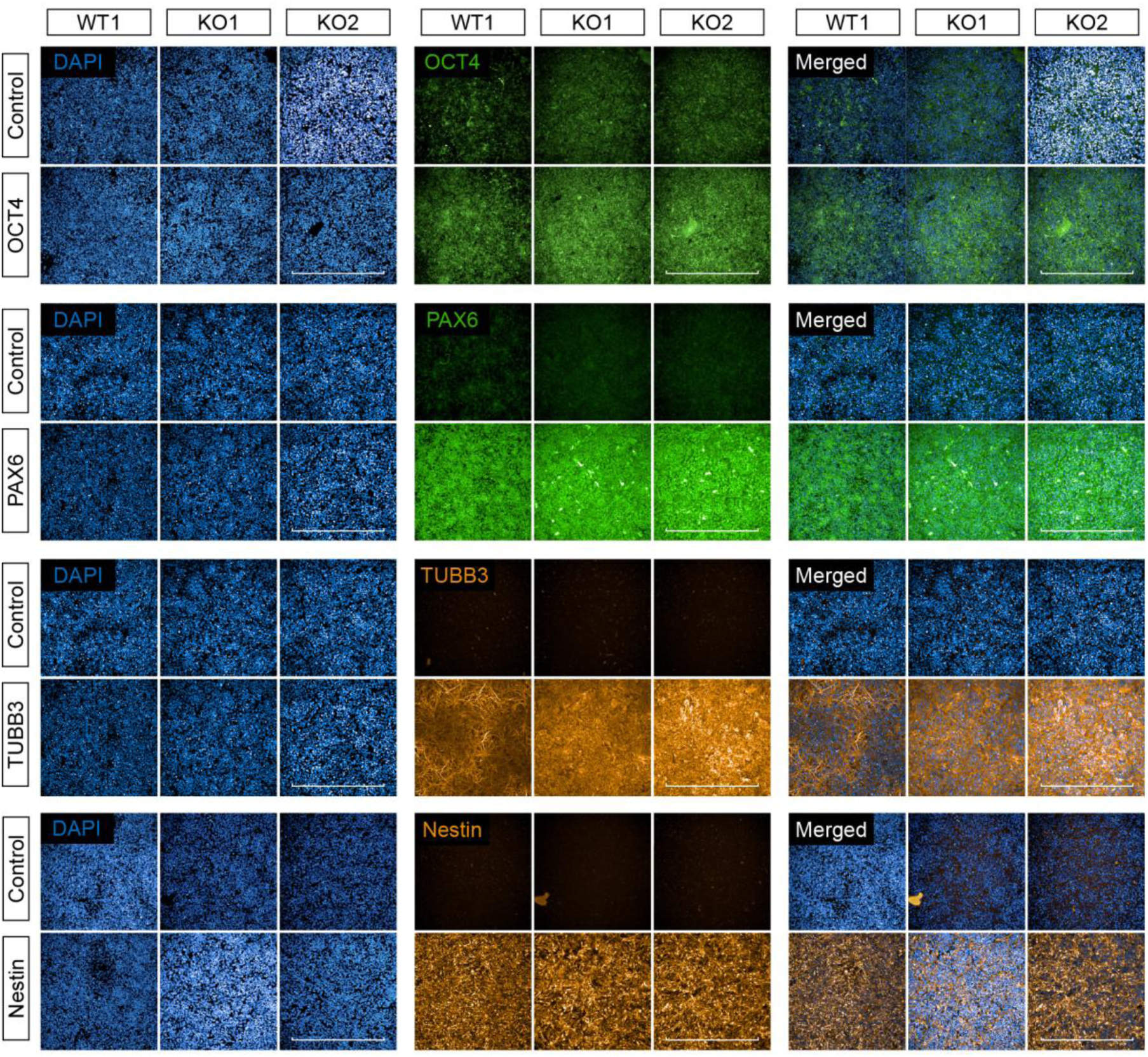
Validation of TSC1 KO NPCs. OCT4, PAX6, TUBB3 and Nestin immunostaining, or negative control staining without the primary antibody, in control (WT1) and TSC1 KO (KO1 and KO2) NPCs. Scale bar: 500µm.

**Supplementary figure S6.**
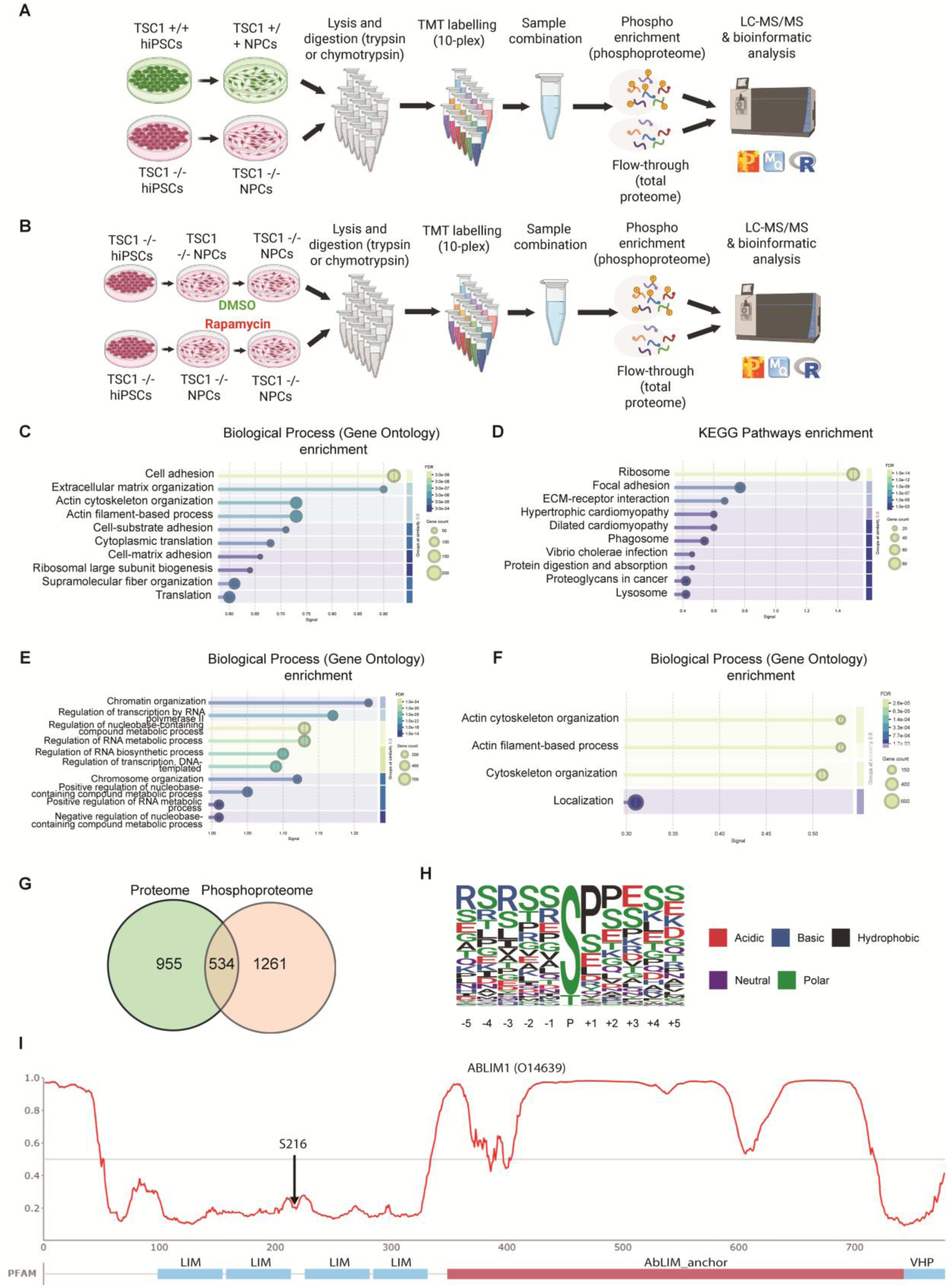
GO and KEGG pathway analysis of proteomic and phosphoproteomic changes in TSC1 KO NPCs. (A) Schematic showing differentiation and LC-MS/MS pipeline for identification of mTORC1 targets in control versus TSC1 KO NPCs. (B) Differentiation and LC-MS/MS pipeline for identification of mTORC1 targets in TSC1 KO NPCs treated with vehicle or rapamycin. (C, D) GO biological process terms (C) and KEGG pathways (D) significantly enriched in proteins with increased expression in TSC1 KO versus control NPCs. (E) GO biological process terms significantly enriched in proteins with decreased expression in TSC1 KO versus control NPCs. (F) GO biological process terms significantly enriched in phosphoproteins with increased expression in TSC1 KO versus control NPCs. (G) Overlap between proteins and phosphoproteins with significantly increased expression in TSC1 KO versus control NPCs. (H) Consensus motif of the mTORC1 target phosphosites identified in human NPCs. (I) Disorder plot of ABLIM1 showing the position of mTORC1 target phosphosite S216 between adjacent LIM domains.

**Supplementary figure S7.**
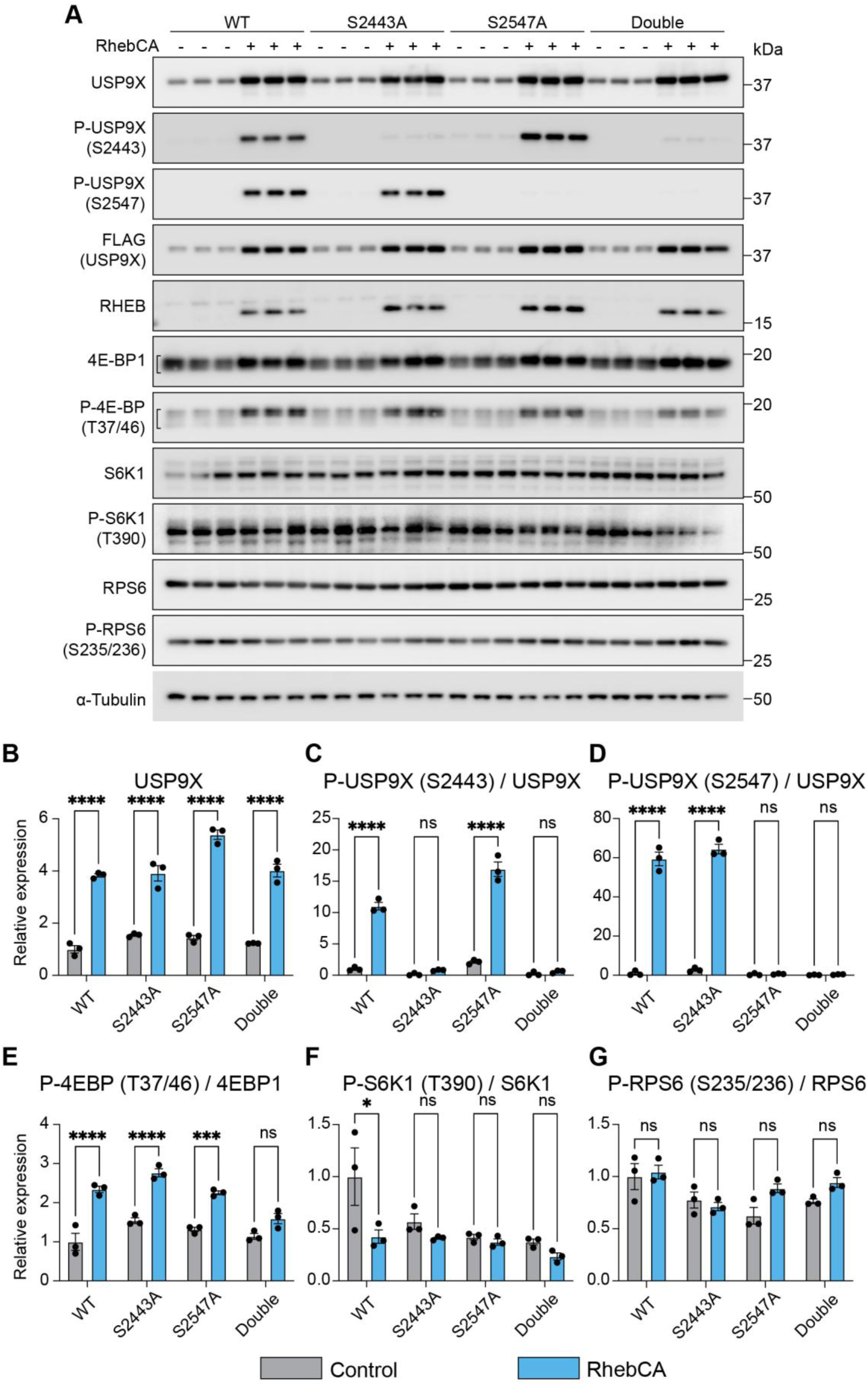
Validation of USP9X S2443 and S2547 phospho-specific antibodies. (A) Western blot analysis of HEK293 cells transfected with a wild-type, S2443A, S2547A or double mutant 309 amino acid fragment (2245-end) of USP9X, together with or without RhebCA to activate mTORC1. (B-D) Quantification of total (B), S2443 (C) and S2547 (D) USP9X expression levels. (E-G) Quantification of P-4E-BP T37/T46 (E), P-S6K T390 (F) and P-RPS6 S240/S244 (G) expression levels. n=3 biological replicates for all conditions. Data are represented as mean ± SEM and were analysed using the students t test. ns not significant, *p < 0.05, ***p < 0.001, ****p < 0.0001.

**Supplementary figure S8.**
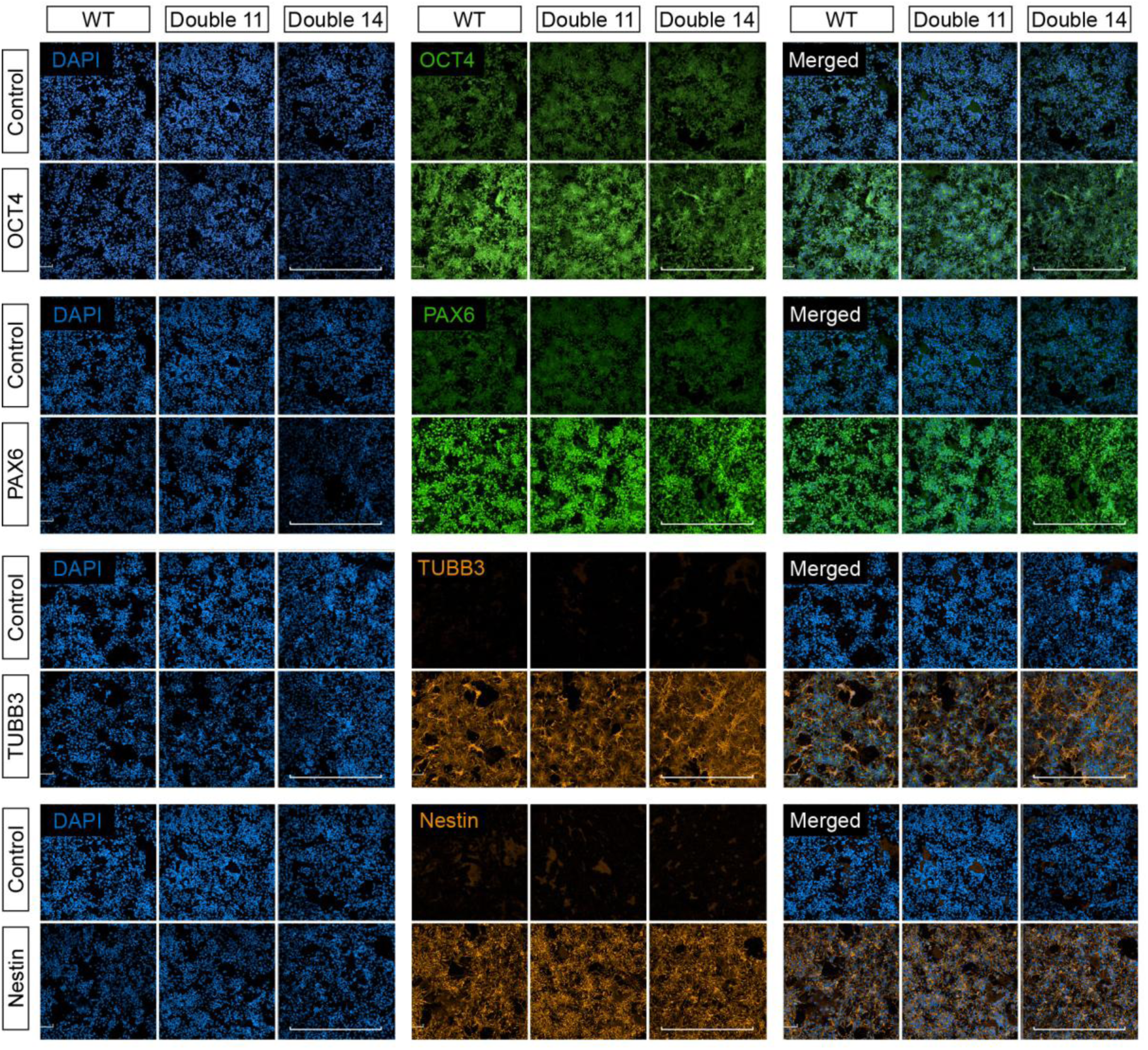
Validation of USP9X phosphomutant NPCs. OCT4, PAX6, TUBB3 and Nestin immunostaining, or negative control staining without the primary antibody, in control (WT) and USP9X phosphomutant (double 11 and double 14) NPCs. Scale bar: 500µm.

**Supplementary figure S9.**
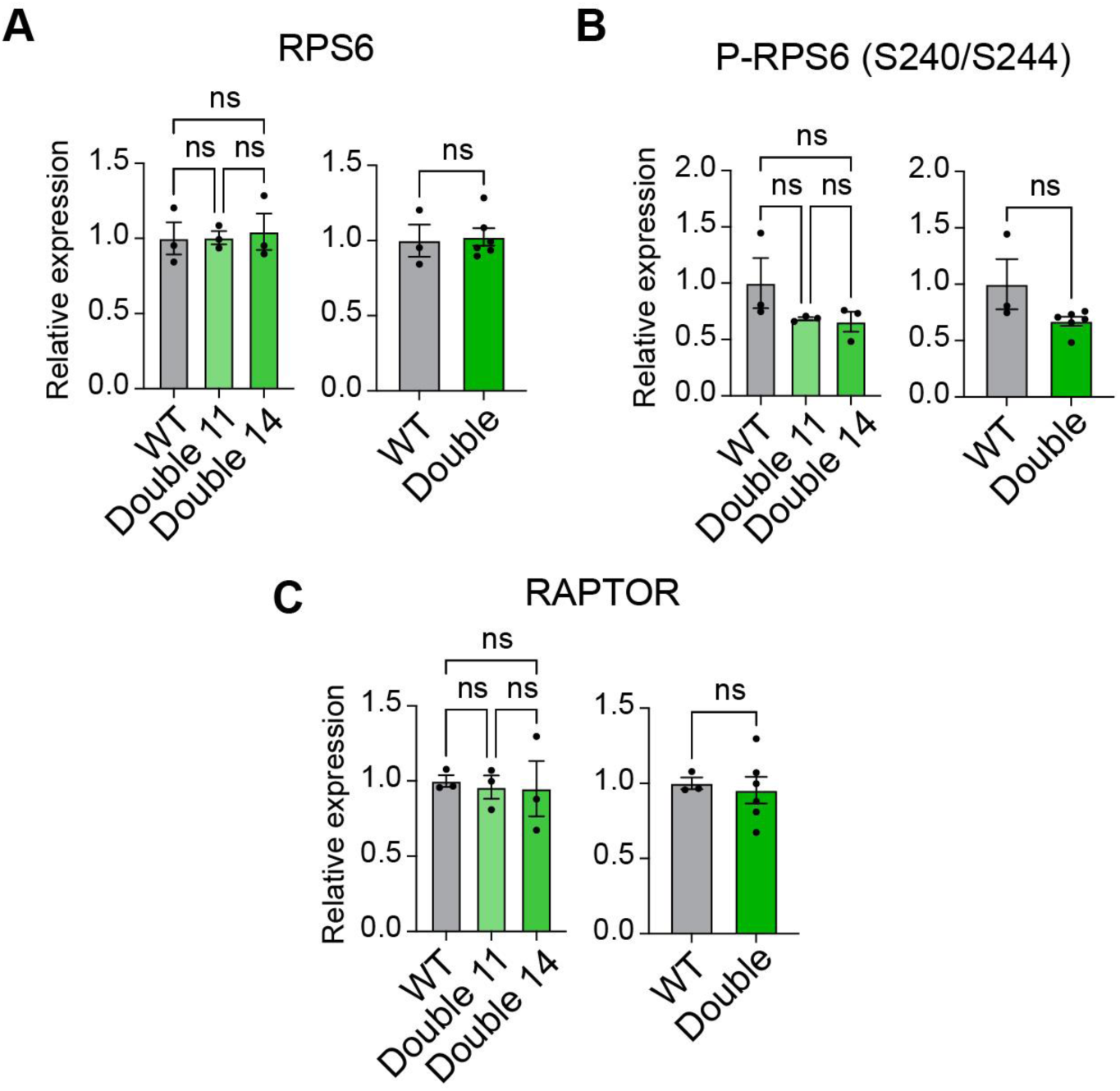
Inhibition of USP9X S2443/S2547 phosphorylation does not affect P-RPS6 or RAPTOR expression levels in NPCs. (A-C) Quantification of total RPS6 (A), P-RPS6 S240/2444 (B) and RAPTOR (C) expression in control (WT) and USP9X hiPSC phosphomutant lines double 11 and double 14. n=3 biological replicates for each condition. Data are represented as mean ± SEM and were analysed using one-way ANOVA. ns not significant.

**Supplementary figure S10.**
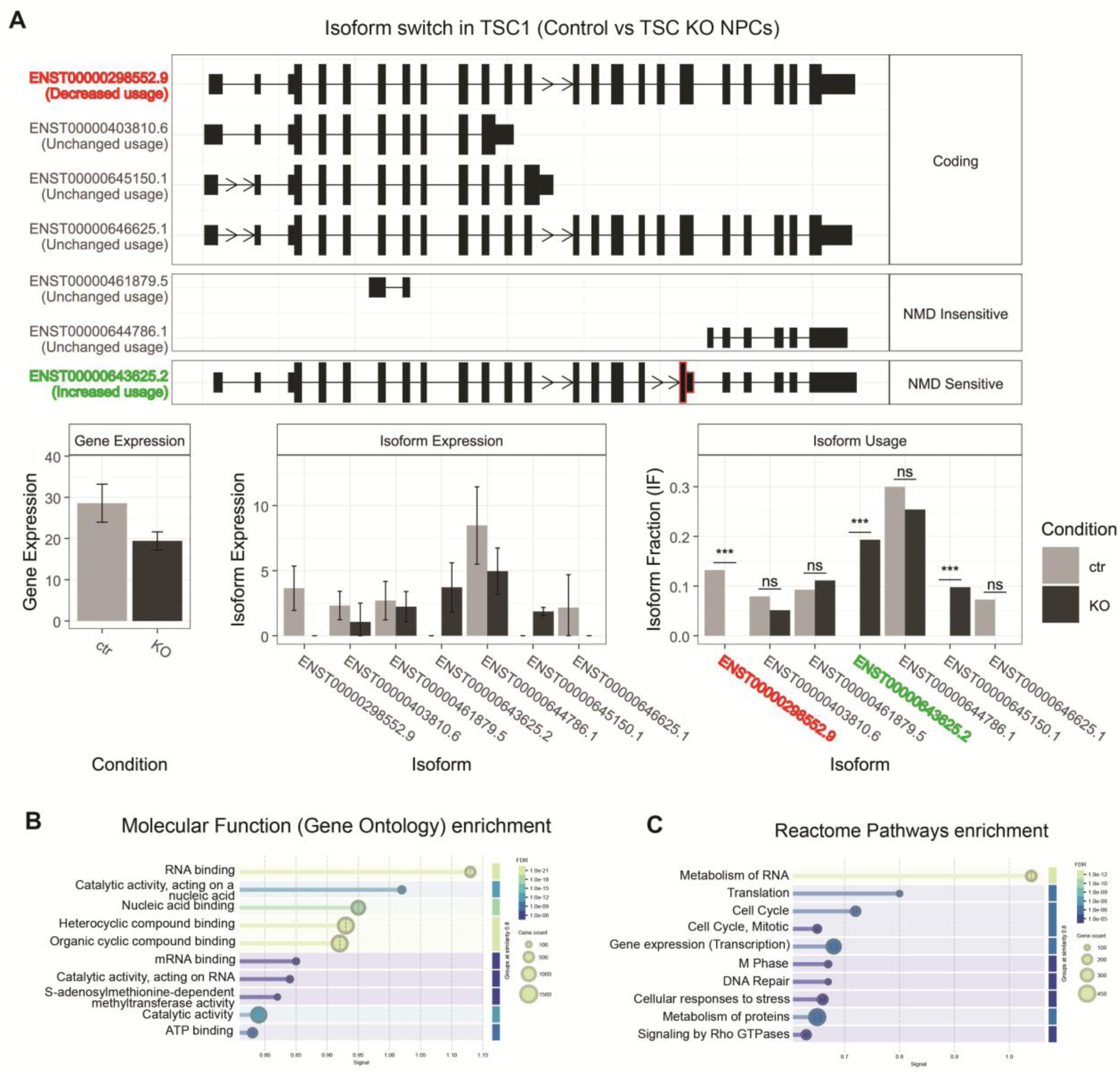
(A) IsoformSwitchAnalyzeR analysis *TSC1* splicing in TSC1 KO versus control NPCs. (B, C) GO molecular function (B) and Reactome pathway (C) analysis of genes with altered splicing in TSC1 KO versus control NPCs. Data are represented as mean ± SEM and were analysed using the students t test. ***p < 0.001

## Supplementary tables

Supplementary tables can be accessed by reviewers through Dryad using this link:

http://datadryad.org/share/LINK_NOT_FOR_PUBLICATION/V-OB3NNIv9M-jEGr6XrEjuj-9pBktow1THfRAMWCT14

Supplementary table S1. Proteomic analysis of E16.5 brain tissue from control versus Tsc1 cKO mice brain (trypsin).

Supplementary table S2. Phosphoproteomic analysis of E16.5 brain tissue from control versus Tsc1 cKO mice brain (trypsin).

Supplementary table S3. Proteomic analysis of E16.5 brain tissue from control versus Tsc1 cKO mice brain (chymotrypsin).

Supplementary table S4. Phosphoproteomic analysis of E16.5 brain tissue from control versus Tsc1 cKO mice brain (chymotrypsin).

Supplementary table S5. Proteomic analysis of E16.5 brain tissue from rapamycin treated versus vehicle treated Tsc1 cKO mice (trypsin).

Supplementary table S6. Phosphoproteomic analysis of E16.5 brain tissue from rapamycin treated versus vehicle treated Tsc1 cKO mice (trypsin).

Supplementary table S7. Proteomic analysis of E16.5 brain tissue from rapamycin treated versus vehicle treated Tsc1 cKO mice (chymotrypsin).

Supplementary table S8. Phosphoproteomic analysis of E16.5 brain tissue from rapamycin treated versus vehicle treated Tsc1 cKO mice (chymotrypsin).

Supplementary table S9. Previously reported mTORC1 targets identified in our analysis. Novel phosphorylation sites on these proteins are underlined.

Supplementary table S10. mTORC1 dependent phosphosites and disorders probability scores in known mTORC1 targets identified in the TSC mouse model.

Supplementary table S11. Cellular functions of the novel mTORC1 targets identified in the mouse TSC model.

Supplementary table S12. Proteomic analysis of TSC1 KO versus control human NPCs (trypsin).

Supplementary table S13. Proteomic analysis of TSC1 KO versus control human NPCs (chymotrypsin).

Supplementary table S14. Phosphoproteomic analysis of TSC1 KO versus control human NPCs (trypsin).

Supplementary table S15. Phosphoproteomic analysis of TSC1 KO versus control human NPCs (chymotrypsin).

Supplementary table S16. Proteomic analysis of TSC1 KO rapamycin versus vehicle treated human NPCs (trypsin).

Supplementary table S17. Proteomic analysis of TSC1 KO rapamycin treated versus vehicle treated human NPCs (chymotrypsin).

Supplementary table S18. Phosphoproteomic analysis of TSC1 KO rapamycin treated versus vehicle treated human NPCs (trypsin).

Supplementary table S19. Phosphoproteomic analysis of TSC1 KO rapamycin treated versus vehicle treated human NPCs (chymotrypsin).

Supplementary table S20. RNA-seq analysis of TSC1 KO versus control human NPCs.

Supplementary table S21. Phosphosites and proteins that were significantly increased in TSC1 KO versus control NPCs and significantly decreased in rapamycin treated TSC1 KO NPCs. Disorders probability scores and whether the phosphosite is significantly increased in SEGA tissue also shown.

Supplementary table S22. Results of rMATS analysis for genes with significantly altered splicing in TSC1 KO versus control human NPCs.

Supplementary table S24. Results of Splicify analysis of TSC1 KO versus control human NPCs using trypsin digest proteomic data. Supplementary table S24. Details of primers and plasmids used.

Supplementary table S25. Details of primary and secondary antibodies used.

## References

1. G. Y. Liu, D. M. Sabatini, mTOR at the nexus of nutrition, growth, ageing and disease. Nature reviews. Molecular cell biology 21, 183–203 (2020).

2. C. H. Aylett et al., Architecture of human mTOR complex 1. Science 351, 48–52 (2016).

3. R. Cafferkey et al., Dominant missense mutations in a novel yeast protein related to mammalian phosphatidylinositol 3-kinase and VPS34 abrogate rapamycin cytotoxicity. Mol Cell Biol 13, 6012–6023 (1993).

4. J. Heitman, N. R. Movva, M. N. Hall, Targets for cell cycle arrest by the immunosuppressant rapamycin in yeast. Science 253, 905–909 (1991).

5. E. J. Brown et al., A mammalian protein targeted by G1-arresting rapamycin-receptor complex. Nature 369, 756–758 (1994).

6. M. I. Chiu, H. Katz, V. Berlin, RAPT1, a mammalian homolog of yeast Tor, interacts with the FKBP12/rapamycin complex. Proc Natl Acad Sci U S A 91, 12574–12578 (1994).

7. D. M. Sabatini, H. Erdjument-Bromage, M. Lui, P. Tempst, S. H. Snyder, RAFT1: a mammalian protein that binds to FKBP12 in a rapamycin-dependent fashion and is homologous to yeast TORs. Cell 78, 35–43 (1994).

8. C. J. Sabers et al., Isolation of a protein target of the FKBP12-rapamycin complex in mammalian cells. J Biol Chem 270, 815–822 (1995).

9. J. O. Lipton, M. Sahin, The neurology of mTOR. Neuron 84, 275–291 (2014).

10. A. R. Tee, J. R. Sampson, D. K. Pal, J. M. Bateman, The role of mTOR signalling in neurogenesis, insights from tuberous sclerosis complex. Semin Cell Dev Biol 52, 12–20 (2016).

11. K. Hentges, K. Thompson, A. Peterson, The flat-top gene is required for the expansion and regionalization of the telencephalic primordium. Development 126, 1601–1609 (1999).

12. J. M. Bateman, H. McNeill, Temporal control of differentiation by the insulin receptor/tor pathway in Drosophila. Cell 119, 87–96 (2004).

13. J. M. Bateman, H. McNeill, Insulin/IGF signalling in neurogenesis. Cellular and Molecular Life Sciences 63, 1701–1705 (2006).

14. A. Avet-Rochex et al., Unkempt is negatively regulated by mTOR and uncouples neuronal differentiation from growth control. PLoS Genet 10, e1004624 (2014).

15. K. T. Maierbrugger, R. Sousa-Nunes, J. M. Bateman, The mTOR pathway component Unkempt regulates neural stem cell and neural progenitor cell cycle in the Drosophila central nervous system. Dev Biol 461, 55–65 (2020).

16. H. McNeill, G. M. Craig, J. M. Bateman, Regulation of neurogenesis and epidermal growth factor receptor signaling by the insulin receptor/target of rapamycin pathway in Drosophila. Genetics 179, 843–853 (2008).

17. F. LiCausi, N. W. Hartman, Role of mTOR Complexes in Neurogenesis. International journal of molecular sciences 19, (2018).

18. A. Polchi, A. Magini, D. D. Meo, B. Tancini, C. Emiliani, mTOR Signaling and Neural Stem Cells: The Tuberous Sclerosis Complex Model. International journal of molecular sciences 19, (2018).

19. M. Girodengo, S. K. Ultanir, J. M. Bateman, Mechanistic target of rapamycin signaling in human nervous system development and disease. Front Mol Neurosci 15, 1005631 (2022).

20. P. B. Crino, mTOR signaling in epilepsy: insights from malformations of cortical development. Cold Spring Harb Perspect Med 5, (2015).

21. L. Mantoan Ritter et al., mTOR pathway diseases: challenges and opportunities from bench to bedside and the mTOR node. Orphanet journal of rare diseases 20, 256 (2025).

22. P. Curatolo, R. Moavero, P. J. de Vries, Neurological and neuropsychiatric aspects of tuberous sclerosis complex. Lancet Neurol 14, 733–745 (2015).

23. H. Northrup, D. A. Krueger, Tuberous sclerosis complex diagnostic criteria update: recommendations of the 2012 Iinternational Tuberous Sclerosis Complex Consensus Conference. Pediatric neurology 49, 243–254 (2013).

24. P. P. Hsu et al., The mTOR-regulated phosphoproteome reveals a mechanism of mTORC1-mediated inhibition of growth factor signaling. Science 332, 1317–1322 (2011).

25. A. M. Robitaille et al., Quantitative phosphoproteomics reveal mTORC1 activates de novo pyrimidine synthesis. Science 339, 1320–1323 (2013).

26. Y. Yu et al., Phosphoproteomic analysis identifies Grb10 as an mTORC1 substrate that negatively regulates insulin signaling. Science 332, 1322–1326 (2011).

27. S. Anderl, M. Freeland, D. J. Kwiatkowski, J. Goto, Therapeutic value of prenatal rapamycin treatment in a mouse brain model of tuberous sclerosis complex. Hum Mol Genet 20, 4597–4604 (2011).

28. P. Baskaran et al., Phosphorylation of the novel mTOR substrate Unkempt regulates cellular morphogenesis. J Biol Chem, 102788 (2022).

29. J. Goto et al., Regulable neural progenitor-specific Tsc1 loss yields giant cells with organellar dysfunction in a model of tuberous sclerosis complex. Proc Natl Acad Sci U S A 108, E1070–1079 (2011).

30. G. Benvenuto et al., The tuberous sclerosis-1 (TSC1) gene product hamartin suppresses cell growth and augments the expression of the TSC2 product tuberin by inhibiting its ubiquitination. Oncogene 19, 6306–6316 (2000).

31. J. Huang, B. D. Manning, The TSC1-TSC2 complex: a molecular switchboard controlling cell growth. The Biochemical journal 412, 179–190 (2008).

32. G. G. Chiang, R. T. Abraham, Phosphorylation of mammalian target of rapamycin (mTOR) at Ser-2448 is mediated by p70S6 kinase. J Biol Chem 280, 25485–25490 (2005).

33. J. Huang, S. Wu, C. L. Wu, B. D. Manning, Signaling events downstream of mammalian target of rapamycin complex 2 are attenuated in cells and tumors deficient for the tuberous sclerosis complex tumor suppressors. Cancer Res 69, 6107–6114 (2009).

34. S. Battaglioni, D. Benjamin, M. Wälchli, T. Maier, M. N. Hall, mTOR substrate phosphorylation in growth control. Cell 185, 1814–1836 (2022).

35. M. Bonucci et al., mTOR and S6K1 drive polycystic kidney by the control of Afadin-dependent oriented cell division. Nature communications 11, 3200 (2020).

36. J. L. Johnson et al., An atlas of substrate specificities for the human serine/threonine kinome. Nature 613, 759–766 (2023).

37. A. Bah, J. D. Forman-Kay, Modulation of Intrinsically Disordered Protein Function by Post-translational Modifications. J Biol Chem 291, 6696–6705 (2016).

38. E. A. Newcombe, E. Delaforge, R. Hartmann-Petersen, K. Skriver, B. B. Kragelund, How phosphorylation impacts intrinsically disordered proteins and their function. Essays in biochemistry 66, 901–913 (2022).

39. K. Nagahama et al., Setd1a Insufficiency in Mice Attenuates Excitatory Synaptic Function and Recapitulates Schizophrenia-Related Behavioral Abnormalities. Cell reports 32, 108126 (2020).

40. M. Murtaza, L. A. Jolly, J. Gecz, S. A. Wood, La FAM fatale: USP9X in development and disease. Cell Mol Life Sci 72, 2075–2089 (2015).

41. J. D. Blair, D. Hockemeyer, H. S. Bateup, Genetically engineered human cortical spheroid models of tuberous sclerosis. Nat Med 24, 1568–1578 (2018).

42. D. Adhya et al., Atypical Neurogenesis in Induced Pluripotent Stem Cells From Autistic Individuals. Biological psychiatry 89, 486–496 (2021).

43. M. Livingstone, M. Bidinosti, Rapamycin-insensitive mTORC1 activity controls eIF4E:4E-BP1 binding. F1000Res 1, 4 (2012).

44. X. Wang, A. Beugnet, M. Murakami, S. Yamanaka, C. G. Proud, Distinct signaling events downstream of mTOR cooperate to mediate the effects of amino acids and insulin on initiation factor 4E-binding proteins. Mol Cell Biol 25, 2558–2572 (2005).

45. R. Blazev et al., Phosphoproteomics of three exercise modalities identifies canonical signaling and C18ORF25 as an AMPK substrate regulating skeletal muscle function. Cell Metab 34, 1561–1577.e1569 (2022).

46. L. Erkman et al., A POU domain transcription factor-dependent program regulates axon pathfinding in the vertebrate visual system. Neuron 28, 779–792 (2000).

47. P. B. Moloney, G. L. Cavalleri, N. Delanty, Epilepsy in the mTORopathies: opportunities for precision medicine. Brain Commun 3, fcab222 (2021).

48. L. Subramanian, M. E. Calcagnotto, M. F. Paredes, Cortical Malformations: Lessons in Human Brain Development. Front Cell Neurosci 13, 576 (2019).

49. M. Girodengo et al., The phosphoproteomic landscape of the neurological manifestations in tuberous sclerosis complex. Acta neuropathologica 151, (2026).

50. A. Bankolé et al., mTOR controls ependymal cell differentiation by targeting the alternative cell cycle and centrosomal proteins. EMBO reports 26, 3075–3105 (2025).

51. P. Agrawal, Y. T. Chen, B. Schilling, B. W. Gibson, R. E. Hughes, Ubiquitin-specific peptidase 9, X-linked (USP9X) modulates activity of mammalian target of rapamycin (mTOR). J Biol Chem 287, 21164–21175 (2012).

52. L. A. Jolly et al., Missense variant contribution to USP9X-female syndrome. NPJ Genom Med 5, 53 (2020).

53. M. R. Reijnders et al., De Novo Loss-of-Function Mutations in USP9X Cause a Female-Specific Recognizable Syndrome with Developmental Delay and Congenital Malformations. Am J Hum Genet 98, 373–381 (2016).

54. C. C. Homan et al., Mutations in USP9X are associated with X-linked intellectual disability and disrupt neuronal cell migration and growth. Am J Hum Genet 94, 470–478 (2014).

55. M. Dietachmayr et al., Antagonistic activities of CDC14B and CDK1 on USP9X regulate WT1-dependent mitotic transcription and survival. Nature communications 11, 1268 (2020).

56. A. Moritz et al., Akt-RSK-S6 kinase signaling networks activated by oncogenic receptor tyrosine kinases. Sci Signal 3, ra64 (2010).

57. L. A. Jolly, V. Taylor, S. A. Wood, USP9X enhances the polarity and self-renewal of embryonic stem cell-derived neural progenitors. Mol Biol Cell 20, 2015–2029 (2009).

58. C. R. Bridges et al., USP9X deubiquitylating enzyme maintains RAPTOR protein levels, mTORC1 signalling and proliferation in neural progenitors. Scientific reports 7, 391 (2017).

59. T. Bonacci, M. J. Emanuele, Dissenting degradation: Deubiquitinases in cell cycle and cancer. Seminars in cancer biology 67, 145–158 (2020).

60. S. D. Sisoudiya et al., Identification of USP9X as a leukemia susceptibility gene. Blood Adv 7, 4563–4575 (2023).

61. S. Shen et al., rMATS: robust and flexible detection of differential alternative splicing from replicate RNA-Seq data. Proc Natl Acad Sci U S A 111, E5593–5601 (2014).

62. K. Vitting-Seerup, A. Sandelin, IsoformSwitchAnalyzeR: analysis of changes in genome-wide patterns of alternative splicing and its functional consequences. *Bioinformatics (Oxford*, England) 35, 4469–4471 (2019).

63. M. A. Komor et al., Identification of Differentially Expressed Splice Variants by the Proteogenomic Pipeline Splicify. Mol Cell Proteomics 16, 1850–1863 (2017).

64. N. W. Hartman et al., mTORC1 targets the translational repressor 4E-BP2, but not S6 kinase 1/2, to regulate neural stem cell self-renewal in vivo. Cell reports 5, 433–444 (2013).

65. T. V. Lin, L. Hsieh, T. Kimura, T. J. Malone, A. Bordey, Normalizing translation through 4E-BP prevents mTOR-driven cortical mislamination and ameliorates aberrant neuron integration. Proc Natl Acad Sci U S A 113, 11330–11335 (2016).

66. B. D. Semple, K. Blomgren, K. Gimlin, D. M. Ferriero, L. J. Noble-Haeusslein, Brain development in rodents and humans: Identifying benchmarks of maturation and vulnerability to injury across species. Prog Neurobiol 106-107, 1–16 (2013).

67. G. L. McKinsey et al., Dlx1&2-dependent expression of Zfhx1b (Sip1, Zeb2) regulates the fate switch between cortical and striatal interneurons. Neuron 77, 83–98 (2013).

68. K. Nikouei, A. B. Muñoz-Manchado, J. Hjerling-Leffler, BCL11B/CTIP2 is highly expressed in GABAergic interneurons of the mouse somatosensory cortex. J Chem Neuroanat 71, 1–5 (2016).

69. S. V. Hegarty, A. M. Sullivan, G. W. O’Keeffe, Zeb2: A multifunctional regulator of nervous system development. Prog Neurobiol 132, 81–95 (2015).

70. S. Gueroussov et al., Regulatory Expansion in Mammals of Multivalent hnRNP Assemblies that Globally Control Alternative Splicing. Cell 170, 324–339.e323 (2017).

71. J. Ule, B. J. Blencowe, Alternative Splicing Regulatory Networks: Functions, Mechanisms, and Evolution. Mol Cell 76, 329–345 (2019).

72. F. Roşianu et al., Loss of NDR1/2 kinases impairs endomembrane trafficking and autophagy leading to neurodegeneration. Life Sci Alliance 6, (2023).

73. X. Jiang et al., Sensitive and Accurate Quantitation of Phosphopeptides Using TMT Isobaric Labeling Technique. Journal of proteome research 16, 4244–4252 (2017).

74. A. J. Storey et al., ProteoViz: a tool for the analysis and interactive visualization of phosphoproteomics data. Mol Omics 16, 316–326 (2020).

75. M. Gierlinski, F. Gastaldello, C. Cole, G. J. Barton, *Proteus*: an R package for downstream analysis of *MaxQuant* output. bioRxiv, 416511 (2018).

76. E. Maes et al., CONSTANd : A Normalization Method for Isobaric Labeled Spectra by Constrained Optimization. Mol Cell Proteomics 15, 2779–2790 (2016).

77. G. Erdős, Z. Dosztányi, AIUPred: combining energy estimation with deep learning for the enhanced prediction of protein disorder. Nucleic Acids Res 52, W176–w181 (2024).

78. D. Szklarczyk et al., The STRING database in 2023: protein-protein association networks and functional enrichment analyses for any sequenced genome of interest. Nucleic Acids Res 51, D638–d646 (2023).

79. C. B. Pantazis et al., A reference human induced pluripotent stem cell line for large-scale collaborative studies. Cell stem cell 29, 1685–1702.e1622 (2022).

80. A. Bhat et al., Attenuated transcriptional response to pro-inflammatory cytokines in schizophrenia hiPSC-derived neural progenitor cells. Brain Behav Immun 105, 82–97 (2022).

81. C. Shum et al., Δ(9)-tetrahydrocannabinol and 2-AG decreases neurite outgrowth and differentially affects ERK1/2 and Akt signaling in hiPSC-derived cortical neurons. Molecular and cellular neurosciences 103, 103463 (2020).

82. P. A. Ewels et al., The nf-core framework for community-curated bioinformatics pipelines. Nature biotechnology 38, 276–278 (2020).

83. M. I. Love, W. Huber, S. Anders, Moderated estimation of fold change and dispersion for RNA-seq data with DESeq2. Genome Biol 15, 550 (2014).

84. J. Schindelin et al., Fiji: an open-source platform for biological-image analysis. Nat Methods 9, 676–682 (2012).

85. Y. Perez-Riverol et al., The PRIDE database at 20 years: 2025 update. Nucleic Acids Res 53, D543–d553 (2025).

86. E. Erkut et al., A cardiovascular, craniofacial, and neurodevelopmental disorder caused by loss-of-function variants in the eIF3 complex component genes EIF3A and EIF3B. Am J Hum Genet 112, 2625–2642 (2025).

87. M. Magaña-Acosta, V. Valadez-Graham, ATRX: From Chromatin Remodeling to Disease. Genesis 63, e70031 (2025).

88. F. Greenberg et al., Multi-disciplinary clinical study of Smith-Magenis syndrome (deletion 17p11.2). American journal of medical genetics 62, 247–254 (1996).

89. V. Endris et al., The novel Rho-GTPase activating gene MEGAP/ srGAP3 has a putative role in severe mental retardation. Proc Natl Acad Sci U S A 99, 11754–11759 (2002).

90. F. Qiao et al., A De Novo heterozygous frameshift mutation identified in BCL11B causes neurodevelopmental disorder by whole exome sequencing. Mol Genet Genomic Med 7, e897 (2019).

91. T. Singh et al., Rare loss-of-function variants in SETD1A are associated with schizophrenia and developmental disorders. Nat Neurosci 19, 571–577 (2016).

92. W. Deb et al., DE NOVO VARIANTS IN THE POLY(RC)-BINDING PROTEIN GENE PCBP1 CAUSE A NEURODEVELOPMENTAL DISORDER. medRxiv, (2025).

93. N. Smal et al., Burden re-analysis of neurodevelopmental disorder cohorts for prioritization of candidate genes. European journal of human genetics : EJHG 32, 1378–1386 (2024).

94. B. Yuan et al., Clinical exome sequencing reveals locus heterogeneity and phenotypic variability of cohesinopathies. Genet Med 21, 663–675 (2019).

95. D. Samanta, SYNGAP1-related disorder: pathophysiology, epilepsy, cognitive and behavioral phenotypes, and precision therapeutic approaches. Epilepsy & behavior : E&B 182, 111098 (2026).

96. S. E. Raible et al., Clinical and molecular spectrum of CHOPS syndrome. Am J Med Genet A 179, 1126–1138 (2019).

97. K. Tatton-Brown et al., Weaver syndrome and EZH2 mutations: Clarifying the clinical phenotype. Am J Med Genet A 161a, 2972–2980 (2013).

98. K. Yamada et al., Nonsense and frameshift mutations in ZFHX1B, encoding Smad-interacting protein 1, cause a complex developmental disorder with a great variety of clinical features. Am J Hum Genet 69, 1178–1185 (2001).

99. N. C. Bramswig et al., Identification of new TRIP12 variants and detailed clinical evaluation of individuals with non-syndromic intellectual disability with or without autism. Hum Genet 136, 179–192 (2017).

100. R. Marin et al., A new developmental and epileptic encephalopathy: PUM1-neurodevelopmental disorder with epilepsy with myoclonic-atonic seizures. Seizure 131, 16–21 (2025).

101. P. M. Tamhankar et al., Lenz-Majewski syndrome: Report of a case with novel mutation in PTDSS1 gene. Eur J Med Genet 58, 392–399 (2015).

102. B. G. Ng et al., Biallelic mutations in CAD, impair de novo pyrimidine biosynthesis and decrease glycosylation precursors. Hum Mol Genet 24, 3050–3057 (2015).

103. C. Wang et al., Mutations in SLC20A2 link familial idiopathic basal ganglia calcification with phosphate homeostasis. Nat Genet 44, 254–256 (2012).

104. Z. El Amrani et al., Complex translocation leading to13q interstitial deletion in a Moroccan child with retinoblastoma and intellectual disability. Ophthalmic Genet 43, 689–692 (2022).

105. X. Jia et al., De novo variants in genes regulating stress granule assembly associate with neurodevelopmental disorders. Sci Adv 8, eabo7112 (2022).

106. R. A. Alkhater, S. W. Scherer, B. A. Minassian, S. Walker, PI4K2A deficiency in an intellectual disability, epilepsy, myoclonus, akathisia syndrome. Annals of clinical and translational neurology 5, 1617–1621 (2018).

