## Supplemental table S25 for "Cross-species phosphoproteomic mapping of mTORC1 signalling reveals RNA splicing regulation during neurodevelopment"

| **Antibody** | **Origin (species)** | **Concentration (western blots)** | **Concentration (ICC)** | **Reference** |
| --- | --- | --- | --- | --- |
| 4E-BP1 | Rabbit | 1:1000-2000 | n/a | 9644; Cell Signaling |
| 4E-BP2 | Rabbit | 1:1000-2000 | 1:400 | 2845; Cell Signaling |
| FLAG | Rat | 1:1000-2000 | n/a | MA1-142; Invitrogen |
| Hamartin (TSC1) | Rabbit | 1:1000 | n/a | ab270967; Abcam |
| Ki67 | Rabbit | n/a | 1:250 | ab15580; Abcam |
| mTOR | Rabbit | 1:2000 | n/a | 2983; Cell Signaling |
| mTOR | Rabbit | 1:2000 | n/a | 2972; Cell Signaling |
| NANOG | Rabbit | n/a | 1:250 | ab109250; Abcam |
| Nestin (10C2) | Mouse | n/a | 1:500 | 14-9843; Invitrogen |
| OCT4 | Rabbit | n/a | 1:1000 | ab19857; Abcam |
| PAX6 | Rabbit | n/a | 1:350 | ab195045; Abcam |
| phospho-4E-BP S65 | Rabbit | 1:2000 | n/a | 9451; Cell Signaling |
| phospho-4E-BP S65 | Rabbit | 1:2000 | n/a | 13443; Cell Signaling |
| phospho-4E-BP T37/46 | Rabbit | 1:1000-2000 | 1:200 | 2855; Cell Signaling |
| Phospho-mTOR S2448 | Rabbit | 1:1000-1:5000 | n/a | 5536; Cell Signaling |
| Phospho-PKCα S657 | Rabbit | 1:1000 | n/a | ab180848; Abcam |
| Phospho-RPS6 S235/236 | Rabbit | 1:1000-2000 | n/a | 2211; Cell Signaling |
| Phospho-RPS6 S240/244 | Rabbit | 1:1000-2000 | n/a | 4838; Cell Signaling |
| phospho-RPS6 S240/244 | Rabbit | 1:1000-2000 | 1:800 | 5364; Cell Signaling |
| Phospho-S6K1 S371 | Rabbit | 1:2000 | n/a | 9208; Cell Signaling |
| Phospho-S6K1 T389 | Rabbit | 1:1000-2000 | n/a | 9234; Cell Signaling |
| Phospho-USP9X S2443 | Rabbit | 1:1000 | n/a | Covalab |
| Phospho-USP9X S2547 | Rabbit | 1:1000 | n/a | Covalab |
| PKCα | Rabbit | 1:1000-12,000 | n/a | ab32376; Abcam |
| RAPTOR | Rabbit | 1:800 | n/a | 2280; Cell Signaling |
| Rheb | Rabbit | 1:3000 | n/a | ab92313; Abcam |
| RPS6 | Rabbit | 1:1000-2000 | 1:250 | 2217; Cell Signaling |
| S6K1 | Rabbit | 1:1000-2000 | n/a | 9202; Cell Signaling |
| SOX2 | Rabbit | n/a | 1:250 | ab97959; Abcam |
| SSEA4 | Mouse | n/a | 1:250 | ab16287; Abcam |
| TRA-1-60 | Mouse | n/a | 1:250 | ab16288; Abcam |
| TUBB3 | Mouse | n/a | 1:200 | 801201; BioLegend |
| Tuberin (TSC2) | Rabbit | 1:1000 | n/a | ab32554; Abcam |
| USP9X | Rabbit | 1:2000 | n/a | A301-351A; Bethyl |
| USP9X | Rabbit | 1:2000 | 1:100 | NBP1-4831; NovusBio |
| α-Tubulin | Mouse | 1:10,000 | n/a | T9026; Sigma-Aldrich |
| THP | Rabbit | 1:5000 | n/a | ab92570; Abcam |

Primary antibodies

Secondary antibodies

| **Antibody** | **Concentration (western blots)** | **Concentration (ICC)** | **Reference** |
| --- | --- | --- | --- |
| Mouse HRP-conjugated | 1:20,000 | n/a | 715-035-151; Jackson |
| Rabbit HRP-conjugated | 1:20,000 | n/a | 711-035-152; Jackson |
| Rat HRP-conjugated | 1:20,000 | n/a | 712-035-153; Jackson |
| Mouse AlexaFluor® 488 | n/a | 1:500-1000 | 715-545-151; Jackson |
| Mouse AlexaFluor® 568 | n/a | 1:500-1000 | A10037; Invitrogen |
| Mouse AlexaFluor® 647 | n/a | 1:500-1000 | 715-605-151; Jackson |
| Rabbit AlexaFluor® 488 | n/a | 1:500-1000 | 711-545-152; Jackson |
| Rabbit AlexaFluor® 568 | n/a | 1:500-1000 | A10042; Invitrogen |
| Rabbit AlexaFluor® 647 | n/a | 1:500-1000 | 711-605-152; Jackson |
