## Supplemental table S24 for "Cross-species phosphoproteomic mapping of mTORC1 signalling reveals RNA splicing regulation during neurodevelopment"

| Primers | | |
| --- | --- | --- |
| **Primer** | **Use** | **Sequence** |
| Nestin-Cre_FWD | Nestin-Cre mice genotyping | TTGCTAAAGCGCTACATAGGA |
| Nestin-Cre_REV | Nestin-Cre mice genotyping | GCCTTATTGTGGAAGGACTG |
| Tsc1^fl/fl^_FWD | *Tsc1^fl/fl^* mice genotyping | GTCACGACCGTAGGAGAAGC |
| Tsc1^fl/fl^_REV | *Tsc1^fl/fl^* mice genotyping | GAATCAACCCCACAGAGCAT |
| S2443A_FWD | S2443A mutagenesis | **G**CTCCCCCAGTGCAAAGCAAT |
| S2443A_REV | S2443A mutagenesis | CCAATTGTTGTAAGTGTACTGAGG  ATTGCC |
| S2547A_FWD | S2547A mutagenesis | CCCCACCTCAAACCAAG |
| S2547A_REV | S2547A mutagenesis | **C**TACTTCTTCACTGCCTTCATA |
| USP9X_Seq_14 | S2443A & S2547A sequencing (plasmids, hiPSCs) | GGAATGGCTTGGAGATGA |
| USP9X_S2443A_FWD | USP9X hiPSC genotyping | CCTGGGCAGTGGAATGGCTTGG |
| USP9X_S2443A_REV | USP9X hiPSC genotyping | GCCACCATGCCCAGCCGAATAA |
| USP9X_S2547A_FWD | USP9X hiPSC genotyping | AGGCCCAGCAGCACATCACATG |
| USP9X_S2547A_REV | USP9X hiPSC genotyping | CACTTTCCACTGCAGGCGGGTT |
| USP9Y_OffTarget_FWD | USP9X hiPSC genotyping | TCCAGATGACAGAGATGGGCTGT |
| USP9Y_OffTarget_REV | USP9X hiPSC genotyping | CCATCTGCTGTCCATTGGTAGCC |
| TSC1_FWD | TSC1 hiPSC genotyping | GCTTCACACCCGCTGTACAAT |
| TSC1_REV | TSC1 hiPSC genotyping | ACACAGCGAAGGAAGTGTCT |
| *TSC1* Exon17 Fwd | TSC1 gRNA | AGACTTCCTGCCAGGCTTCCCT |
| *TSC1* Exon 17 Rev | TSC1 gRNA | CTGACTGGCTTCACACCCGCTG |
| attB_Fragment_FWD | Gateway cloning | GGGGACAAGTTTGTACAAAAAAGC  AGGCTTAATGCAGTCTTCAATCAAT  GGTAATCCTC |
| attB_Fragment_REV | Gateway cloning | GGGGACCACTTTGTACAAGAAAGC  TGGGTTTCATTGATCCTTGGTTTGA  GGTGGG |
| M13 *TSC1* Exon 17 Fwd | MinION sequencing | GTAAAACGACGGCCAAGACTTCCTGCCAGGCTTCCCT |
| M13 *TSC1* Exon 17 Rev | MinION sequencing | GGAAACAGCTATGACCATGCTGACTGGCTTCACACCCGCTG |
| Barcode Fwd 1 | MinION sequencing | AAGAAAGTTGTCGGTGTCTTTGTGGTAAAACGACGGCCA |
| Barcode Fwd 2 | MinION sequencing | TCGATTCCGTTTGTAGTCGTCTGTGTAAAACGACGGCCA |
| Barcode Fwd 3 | MinION sequencing | GAGTCTTGTGTCCCAGTTACCAGGGTAAAACGACGGCCA |
| Barcode Fwd 4 | MinION sequencing | TTCGGATTCTATCGTGTTTCCCTAGTAAAACGACGGCCA |
| Barcode Fwd 5 | MinION sequencing | CTTGTCCAGGGTTTGTGTAACCTTGTAAAACGACGGCCA |
| Barcode Fwd 6 | MinION sequencing | TTCTCGCAAAGGCAGAAAGTAGTCGTAAAACGACGGCCA |
| Barcode Fwd 7 | MinION sequencing | GTGTTACCGTGGGAATGAATCCTTGTAAAACGACGGCCA |
| Barcode Fwd 8 | MinION sequencing | TTCAGGGAACAAACCAAGTTACGTGTAAAACGACGGCCA |
| Barcode Fwd 9 | MinION sequencing | AACTAGGCACAGCGAGTCTTGGTTGTAAAACGACGGCCA |
| Barcode Fwd 10 | MinION sequencing | AAGCGTTGAAACCTTTGTCCTCTCGTAAAACGACGGCCA |
| Barcode Fwd 11 | MinION sequencing | GTTTCATCTATCGGAGGGAATGGAGTAAAACGACGGCCA |
| Barcode Fwd 12 | MinION sequencing | CAGGTAGAAAGAAGCAGAATCGGAGTAAAACGACGGCCA |
| Barcode Rev A | MinION sequencing | AGAACGACTTCCATACTCGTGTGAGGAAACAGCTATGACCATG |
| Barcode Rev B | MinION sequencing | AACGAGTCTCTTGGGACCCATAGAGGAAACAGCTATGACCATG |
| Barcode Rev C | MinION sequencing | AGGTCTACCTCGCTAACACCACTGGGAAACAGCTATGACCATG |
| Barcode Rev D | MinION sequencing | CGTCAACTGACAGTGGTTCGTACTGGAAACAGCTATGACCATG |
| Barcode Rev E | MinION sequencing | ACCCTCCAGGAAAGTACCTCTGATGGAAACAGCTATGACCATG |
| Barocde Rev F | MinION sequencing | CCAAACCCAACAACCTAGATAGGCGGAAACAGCTATGACCATG |
| Barcode Rev G | MinION sequencing | GTTCCTCGTGCAGTGTCAAGAGATGGAAACAGCTATGACCATG |
| Barcode Rev H | MinION sequencing | TTGCGTCCTGTTACGAGAACTCATGGAAACAGCTATGACCATG |

| **Vectors/Plasmids** | **Antibiotic resistance** | **Uses** | **Source** |
| --- | --- | --- | --- |
| pCAG | Ampicillin | Transfection (HEK293T) | Gift from Dr Matthias Krause |
| pCAG-mCherry | Ampicillin | Transfection (HEK293T) | Gift from Dr Molly Strom |
|  |  | CoIP |  |
| pCAG-RhebCA | Ampicillin | Transfection (HEK293T) | Gift from Dr Angelique Bordey |
|  |  | CoIP |  |
| CMV-FLAG | Ampicillin | Transfection (HEK293T) | Gift from Prof Guillaume Hautbergue |
|  |  | CoIP |  |
| CMV-GFP | Ampicillin | Transfection (HEK293T) | Gift from Dr Molly Strom |
|  |  | CoIP |  |
| pEGFPC1-USP9X-WT | Neomycin-Kanamycin | Transfection (HEK293T) | DU10181; MRC PPU |
|  |  | CoIP |  |
|  |  | Site-directed mutagenesis |  |
| pEGFPC1-USP9X-S2443A | Neomycin-Kanamycin | Site-directed mutagenesis | Generated via site-directed mutagenesis from pEGFPC1-USP9X-WT |
| pEGFPC1-USP9X-Double | Neomycin-Kanamycin | Transfection (HEK293T) | Generated via site-directed mutagenesis from pEGFPC1-USP9X-S2443A |
|  |  | CoIP |  |
| pCDNA5-FRT-FLAG-(USP9X aa2245-end)-WT | Ampicillin | Transfection (HEK293T) | DU30069; MRC PPU |
|  |  | Site-directed mutagenesis |  |
|  |  | Gateway cloning |  |
| pCDNA5-FRT-FLAG-(USP9X aa2245-end)-S2443A | Ampicillin | Transfection (HEK293T) | Generated via site-directed mutagenesis from pCDNA5-FRT-FLAG-(USP9X aa2245-end)-WT |
|  |  | Site-directed mutagenesis |  |
|  |  | Gateway cloning |  |
| pCDNA5-FRT-FLAG-(USP9X aa2245-end)-S2547A | Ampicillin | Transfection (HEK293T) | Generated via site-directed mutagenesis from pCDNA5-FRT-FLAG-(USP9X aa2245-end)-WT |
|  |  | Gateway cloning |  |
| pCDNA5-FRT-FLAG-(USP9X aa2245-end)-Double | Ampicillin | Transfection (HEK293T) | Generated via site-directed mutagenesis from pCDNA5-FRT-FLAG-(USP9X aa2245-end)-S2443A |
|  |  | Gateway cloning |  |
| pDONR^™^ 207 (attP1 and attP2) | Gentamycin | Gateway cloning | Invitrogen |
| pDest-565 (T7-6xHis/GST) | Ampicillin | Gateway cloning | 11520; Addgene |
| T7-6xHis/GST-(USP9X aa2245-end)-WT | Ampicillin | Bacterial protein purification | Generated via Gateway cloning using pCDNA5-FRT-FLAG-(USP9X aa2245-end)-WT |
| T7-6xHis/GST-(USP9X aa2245-end)-S2443A | Ampicillin | Bacterial protein purification | Generated via Gateway cloning using pCDNA5-FRT-FLAG-(USP9X aa2245-end)-S2443A |
| T7-6xHis/GST-(USP9X aa2245-end)-S2547A | Ampicillin | Bacterial protein purification | Generated via Gateway cloning using pCDNA5-FRT-FLAG-(USP9X aa2245-end)-S2547A |
| T7-6xHis/GST-(USP9X aa2245-end)-Double | Ampicillin | Bacterial protein purification | Generated via Gateway cloning using pCDNA5-FRT-FLAG-(USP9X aa2245-end)-Double |
