## Supplemental table S11 for "Cross-species phosphoproteomic mapping of mTORC1 signalling reveals RNA splicing regulation during neurodevelopment"

|  | **mTORC1 target** | **Full name** | **Cellular functions** | **References** |
| --- | --- | --- | --- | --- |
| Transcriptional regulation | ATRX | Transcriptional regulator ATRX | Transcriptional regulation and chromatin remodeling | (Episkopou, Draskovic et al. 2014) |
|  | BCL11B (also known as CTIP2) | B-cell lymphoma/leukemia 11B  (COUP-TF-interacting protein 2) | Transcription regulation | (García-Aznar, Alonso Alvarez et al. 2024) |
|  | ESF1 | 18S rRNA factor 1 | Transcription regulation | (Peng, Krogan et al. 2004) |
|  | GPBP1 | Vasculin | Transactivating transcription factor | (Bijnens, Gils et al. 2003) |
|  | HDGFL2 | Hepatoma-derived growth factor-related protein 2 | Epigenetic regulation, DNA repair | (Baude, Aaes et al. 2016, Zhu, Lan et al. 2020) |
|  | NSD3 | Nuclear SET domain-containing protein 3 | Epigenetic regulation | (Rathert 2021) |
|  | PDS5A | Sister chromatid cohesion protein PDS5 homolog A | Epigenetic regulation, DNA repair, DNA replication | (Zhang, Coutinho et al. 2021) |
|  | RFX7 | Regulatory factor X7 | Transcription regulation | (Castro, Chelbi et al. 2018) |
|  | SETD1A | SET domain-containing protein 1A | Transcription regulation, Epigenetic regulation, DNA repair | (Alsulami, Munawar et al. 2019) |
| RNA metabolism | STRAP | Serine-threonine kinase receptor-associated protein | spliceosome-associated factor, translation, scaffolding factor | (Karfa 2025) |
|  | PCBP1 | Poly(rC)-binding protein 1 | Multifunctional RNA-binding protein | (Rithvik, Wadhavane et al. 2025) |
|  | PCBP2 | Poly(rC)-binding protein 2 | Multifunctional RNA-binding protein | (Yuan, Chen et al. 2021) |
|  | PYM1 | Partner of Y14 and mago | Disassembly factor for the Exon Junction Complex | (Gehring, Lamprinaki et al. 2009) |
|  | SMG7 | Nonsense-mediated mRNA decay factor SMG7 | Nonsense-mediated mRNA decay | (Schweingruber, Rufener et al. 2013) |
| Autophagy | ATG2B | Autophagy-related 2B | Autophagy, Lipid transport | (Zheng, Li et al. 2017, Valverde, Yu et al. 2019) |
|  | SMCR8 | Smith-Magenis syndrome chromosomal region candidate gene 8 protein | Autophagy | (Jung and Behrends 2020) |
| Cytoskeleton modulation | NAV1 | Neuron navigator 1 | Microtubule cytoskeleton dynamics regulation | (Powers, Hevner et al. 2022, Sandeep, Sharma et al. 2023) |
|  | SRGAP3 | SLIT-ROBO Rho GTPase-activating protein 3 | Cytoskeletal reorganisation | (Bacon, Endris et al. 2013) |
| Deubiquiti-nation | USP33 | Ubiquitin-specific protease 33 | Deubiquitination, Cell cycle | (Zhu, Chen et al. 2025) |
|  | USP9X | Ubiquitin-specific peptidase 9, X-linked | Deubiquitination, Cell cycle, Cell survival, Cytoskeleton dynamics | (Murtaza, Jolly et al. 2015, Meng, Hong et al. 2023) |
| Cell division | AHCTF1 | Elys | Nuclear pore complex assembly | (Shevelyov 2020) |
|  | CCSER1 | Serine-rich coiled-coil domain-containing protein 1 | Cell division | (Patel, Scrimieri et al. 2013, Kang and Park 2019) |
|  | NUMA1 | Nuclear mitotic apparatus protein 1 | Mitotic spindle assembly | (Kiyomitsu and Boerner 2021) |
| Translation | EIF2AK3 (PERK) | Eukaryotic translation initiation factor 2-alpha kinase 3 | Phosphorylates eIF2α in the unfolded protein response | (Hetz 2012) |
|  | EIF3A | Eukaryotic initiation factor 3A | Translation initiation | (Masutani, Sonenberg et al. 2007, Lee, Kranzusch et al. 2015) |
|  | LRRC47 | Leucine-rich repeat-containing protein 47 | Translation regulation | (Ameismeier, Zemp et al. 2020) |
|  | PRRC2B | Protein PRRC2B | RNA-binding protein; protein translation | (Jiang, Hedaya et al. 2023) |
| Others | AAK1 | AP2-associated protein kinase-1 | Endocytosis | (Conner and Schmid 2002) |
|  | C2CD5 | C2 domain-containing protein 5 | Protein trafficking | (Gavini, Cook et al. 2020) |
|  | SEC14L1 | SEC14-like protein 1 | Innate immunity, Cholinergic transporter regulation | (Ribeiro, Ferreira et al. 2007, Li, Di et al. 2013) |
|  | SGTA | Small glutamine-rich tetratricopeptide repeat-containing protein alpha | Misfolded protein chaperone | Philp, Butler et al. (2013) |
|  | SYAP1 | Synapse-associated protein 1 | Promotes mTORC2-mediated phosphorylation of AKT1 | (C, Schmitt et al. 2019) |

Supplementary table S11. Cellular functions of novel mTORC1 targets.

**References**

. "Alzheimer’s Disease International World Alzheimer Report 2010." from <http://www.alz.co.uk/research/worldreport>.

Alsulami, M., N. Munawar, E. Dillon, G. Oliviero, K. Wynne, M. Alsolami, C. Moss, O. G. P, F. O'Meara, D. Cotter and G. Cagney (2019). "SETD1A Methyltransferase Is Physically and Functionally Linked to the DNA Damage Repair Protein RAD18." Mol Cell Proteomics **18**(7): 1428-1436.

Ameismeier, M., I. Zemp, J. van den Heuvel, M. Thoms, O. Berninghausen, U. Kutay and R. Beckmann (2020). "Structural basis for the final steps of human 40S ribosome maturation." Nature **587**(7835): 683-687.

Bacon, C., V. Endris and G. A. Rappold (2013). "The cellular function of srGAP3 and its role in neuronal morphogenesis." Mech Dev **130**(6-8): 391-395.

Baude, A., T. L. Aaes, B. Zhai, N. Al-Nakouzi, H. Z. Oo, M. Daugaard, M. Rohde and M. Jaattela (2016). "Hepatoma-derived growth factor-related protein 2 promotes DNA repair by homologous recombination." Nucleic Acids Res **44**(5): 2214-2226.

Bijnens, A. P., A. Gils, B. Jutten, B. C. Faber, S. Heeneman, P. J. Kitslaar, J. H. Tordoir, C. J. de Vries, A. A. Kroon, M. J. Daemen and K. B. Cleutjens (2003). "Vasculin, a novel vascular protein differentially expressed in human atherogenesis." Blood **102**(8): 2803-2810.

C, R. v. C., D. Schmitt, T. Rülicke, M. Sendtner, R. Blum and E. Buchner (2019). "An essential role of the mouse synapse-associated protein Syap1 in circuits for spontaneous motor activity and rotarod balance." Biol Open **8**(6).

Castro, W., S. T. Chelbi, C. Niogret, C. Ramon-Barros, S. P. M. Welten, K. Osterheld, H. Wang, G. Rota, L. Morgado, E. Vivier, M. E. Raeber, O. Boyman, M. Delorenzi, D. Barras, P. C. Ho, A. Oxenius and G. Guarda (2018). "The transcription factor Rfx7 limits metabolism of NK cells and promotes their maintenance and immunity." Nat Immunol **19**(8): 809-820.

Conner, S. D. and S. L. Schmid (2002). "Identification of an adaptor-associated kinase, AAK1, as a regulator of clathrin-mediated endocytosis." J Cell Biol **156**(5): 921-929.

Episkopou, H., I. Draskovic, A. Van Beneden, G. Tilman, M. Mattiussi, M. Gobin, N. Arnoult, A. Londoño-Vallejo and A. Decottignies (2014). "Alternative Lengthening of Telomeres is characterized by reduced compaction of telomeric chromatin." Nucleic Acids Res **42**(7): 4391-4405.

García-Aznar, J. M., S. Alonso Alvarez and T. Bernal Del Castillo (2024). "Pivotal role of BCL11B in the immune, hematopoietic and nervous systems: a review of the BCL11B-associated phenotypes from the genetic perspective." Genes Immun **25**(3): 232-241.

Gavini, C. K., T. M. Cook, D. J. Rademacher and V. Mansuy-Aubert (2020). "Hypothalamic C2-domain protein involved in MC4R trafficking and control of energy balance." Metabolism **102**: 153990.

Gehring, N. H., S. Lamprinaki, A. E. Kulozik and M. W. Hentze (2009). "Disassembly of exon junction complexes by PYM." Cell **137**(3): 536-548.

Hetz, C. (2012). "The unfolded protein response: controlling cell fate decisions under ER stress and beyond." Nat Rev Mol Cell Biol **13**(2): 89-102.

Jiang, F., O. M. Hedaya, E. Khor, J. Wu, M. Auguste and P. Yao (2023). "RNA binding protein PRRC2B mediates translation of specific mRNAs and regulates cell cycle progression." Nucleic Acids Res **51**(11): 5831-5846.

Jung, J. and C. Behrends (2020). "Multifaceted role of SMCR8 as autophagy regulator." Small GTPases **11**(1): 53-61.

Kang, S. U. and J. T. Park (2019). "Functional evaluation of alternative splicing in the FAM190A gene." Genes Genomics **41**(2): 193-199.

Karfa, S., Saurav, S., Feng, B., Li, S., Law, B.K., Datta, P.K. (2025). "The Role of Serine-Threonine Kinase Receptor-Associated Protein (STRAP) Signaling in Cancer." Cells **14**: 854.

Kiyomitsu, T. and S. Boerner (2021). "The Nuclear Mitotic Apparatus (NuMA) Protein: A Key Player for Nuclear Formation, Spindle Assembly, and Spindle Positioning." Front Cell Dev Biol **9**: 653801.

Lee, A. S., P. J. Kranzusch and J. H. Cate (2015). "eIF3 targets cell-proliferation messenger RNAs for translational activation or repression." Nature **522**(7554): 111-114.

Li, M. T., W. Di, H. Xu, Y. K. Yang, H. W. Chen, F. X. Zhang, Z. H. Zhai and D. Y. Chen (2013). "Negative regulation of RIG-I-mediated innate antiviral signaling by SEC14L1." J Virol **87**(18): 10037-10046.

Masutani, M., N. Sonenberg, S. Yokoyama and H. Imataka (2007). "Reconstitution reveals the functional core of mammalian eIF3." EMBO J **26**(14): 3373-3383.

Meng, Y., C. Hong, S. Yang, Z. Qin, L. Yang and Y. Huang (2023). "Roles of USP9X in cellular functions and tumorigenesis (Review)." Oncol Lett **26**(6): 506.

Murtaza, M., L. A. Jolly, J. Gecz and S. A. Wood (2015). "La FAM fatale: USP9X in development and disease." Cell Mol Life Sci **72**(11): 2075-2089.

Patel, K., F. Scrimieri, S. Ghosh, J. Zhong, M. S. Kim, Y. R. Ren, R. A. Morgan, C. A. Iacobuzio-Donahue, A. Pandey and S. E. Kern (2013). "FAM190A deficiency creates a cell division defect." Am J Pathol **183**(1): 296-303.

Peng, W. T., N. J. Krogan, D. P. Richards, J. F. Greenblatt and T. R. Hughes (2004). "ESF1 is required for 18S rRNA synthesis in Saccharomyces cerevisiae." Nucleic Acids Res **32**(6): 1993-1999.

Philp, L. K., M. S. Butler, T. E. Hickey, L. M. Butler, W. D. Tilley and T. K. Day (2013). "SGTA: a new player in the molecular co-chaperone game." Horm Cancer **4**(6): 343-357.

Powers, R. M., R. F. Hevner and S. Halpain (2022). "The Neuron Navigators: Structure, function, and evolutionary history." Front Mol Neurosci **15**: 1099554.

Rathert, P. (2021). "Structure, Activity and Function of the NSD3 Protein Lysine Methyltransferase." Life (Basel) **11**(8).

Ribeiro, F. M., L. T. Ferreira, S. Marion, S. Fontes, M. Gomez, S. S. Ferguson, M. A. Prado and V. F. Prado (2007). "SEC14-like protein 1 interacts with cholinergic transporters." Neurochem Int **50**(2): 356-364.

Rithvik, A., S. Wadhavane and M. Rasool (2025). "Decoding poly (RC)-binding protein 1 (PCBP1), the underrated guard at the foothill of ferroptosis." Pathol Res Pract **266**: 155771.

Sandeep, P., P. Sharma, K. Luhach, N. Dhiman, H. Kharkwal and B. Sharma (2023). "Neuron navigators: A novel frontier with physiological and pathological implications." Mol Cell Neurosci **127**: 103905.

Schweingruber, C., S. C. Rufener, D. Zünd, A. Yamashita and O. Mühlemann (2013). "Nonsense-mediated mRNA decay - mechanisms of substrate mRNA recognition and degradation in mammalian cells." Biochim Biophys Acta **1829**(6-7): 612-623.

Shevelyov, Y. Y. (2020). "The Role of Nucleoporin Elys in Nuclear Pore Complex Assembly and Regulation of Genome Architecture." Int J Mol Sci **21**(24).

Valverde, D. P., S. Yu, V. Boggavarapu, N. Kumar, J. A. Lees, T. Walz, K. M. Reinisch and T. J. Melia (2019). "ATG2 transports lipids to promote autophagosome biogenesis." J Cell Biol **218**(6): 1787-1798.

Yuan, C., M. Chen and X. Cai (2021). "Advances in poly(rC)-binding protein 2: Structure, molecular function, and roles in cancer." Biomed Pharmacother **139**: 111719.

Zhang, N., L. E. Coutinho and D. Pati (2021). "PDS5A and PDS5B in Cohesin Function and Human Disease." Int J Mol Sci **22**(11).

Zheng, J. X., Y. Li, Y. H. Ding, J. J. Liu, M. J. Zhang, M. Q. Dong, H. W. Wang and L. Yu (2017). "Architecture of the ATG2B-WDR45 complex and an aromatic Y/HF motif crucial for complex formation." Autophagy **13**(11): 1870-1883.

Zhu, X., B. Lan, X. Yi, C. He, L. Dang, X. Zhou, Y. Lu, Y. Sun, Z. Liu, X. Bai, K. Zhang, B. Li, M. J. Li, Y. Chen and L. Zhang (2020). "HRP2-DPF3a-BAF complex coordinates histone modification and chromatin remodeling to regulate myogenic gene transcription." Nucleic Acids Res **48**(12): 6563-6582.

Zhu, Y., Z. Chen, K. Niu, M. Li, Y. Deng, J. Zhang, D. Wei, J. Wang and Y. Zhao (2025). "USP33 Regulates DNA Damage Response and Carcinogenesis Through Deubiquitylating and Stabilising p53." Cell Prolif **58**(5): e13793.
