## Supplemental table S9 for "Cross-species phosphoproteomic mapping of mTORC1 signalling reveals RNA splicing regulation during neurodevelopment"

| **mTORC1 target** | **Full name** | **Direct or indirect** | **Site(s) in the literature** | **Site(s) identified in this study** | **References** |
| --- | --- | --- | --- | --- | --- |
| DNAJC2 | DnaJ homolog subfamily C member 2 | Indirect | S47 | S47, T48, S49 | (Barilari, Bonfils et al. 2017) |
| EEF2K | Eukaryotic elongation factor 2 kinase | Direct | S78, S396 | S76, S77 (human S77, S78) | (Battaglioni, Benjamin et al. 2022) |
| 4E-BP2 | Eukaryotic translation initiation factor 4E-binding protein 2 | Direct | T37, T46, S65, T70 | T46, S65, T70 | (Battaglioni, Benjamin et al. 2022) |
| EIF4G1 | Eukaryotic translation initiation factor 4 gamma 1 | Direct & Indirect | S1108, S1148, S1189, S1192 | S1102 (human S1148) | (Raught, Gingras et al. 2000, Ramirez-Valle, Braunstein et al. 2008) |
| FLCN | Folliculin | Indirect | S302 | S302, T313 | (Piao, Kobayashi et al. 2009) |
| LARP1 | La-related protein 1 | Direct | 26 S/T residues, including S766 and S774 | S743 (human S766),  S751 (human S774) | (Fonseca, Jia et al. 2018, Jia, Lahr et al. 2021, Battaglioni, Benjamin et al. 2022) |
| MAF1 | Repressor of RNA polymerase III transcription MAF1 homolog | Direct | S60, S68, S75 | S75 | (Battaglioni, Benjamin et al. 2022) |
| MTOR | Serine/threonine-protein kinase mTOR | Indirect | S2448 | S2448 | (Holz and Blenis 2005) |
| RPS6 | Ribosomal protein S6 | Indirect | S235, S236, S240, S244 | S235, S236, S240, T241, S242, S244, S246, S247 | (Biever, Valjent et al. 2015) |
| S6K1 | Ribosomal protein S6 kinase β1 | Direct | S371 (human S394), T389, T412 | T390 (mouse T389, human T389),  S398, T399 | (Battaglioni, Benjamin et al. 2022) |
| SQSTM1 (also known as P62) | Sequestosome-1 | Direct | S351 (human S349) | S357 (human S355) | (Ichimura, Waguri et al. 2013, Battaglioni, Benjamin et al. 2022) |
| SRPK2 | SRSF protein kinase 2 | Indirect | S494 | S488 (human S494), S490, S491, T492 | (Lee, Zheng et al. 2017) |

Supplementary table S9. Previously reported mTORC1 targets identified in this study. Novel phosphorylation sites on these proteins identified are underlined.

Barilari, M., G. Bonfils, C. Treins, V. Koka, D. De Villeneuve, S. Fabrega and M. Pende (2017). "ZRF1 is a novel S6 kinase substrate that drives the senescence programme." EMBO J **36**(6): 736-750.

Battaglioni, S., D. Benjamin, M. Wälchli, T. Maier and M. N. Hall (2022). "mTOR substrate phosphorylation in growth control." Cell **185**(11): 1814-1836.

Biever, A., E. Valjent and E. Puighermanal (2015). "Ribosomal Protein S6 Phosphorylation in the Nervous System: From Regulation to Function." Front Mol Neurosci **8**: 75.

Fonseca, B. D., J.-J. Jia, A. K. Hollensen, R. Pointet, H.-D. Hoang, M. R. Niklaus, I. A. Pena, R. M. Lahr, E. M. Smith, J. Hearnden, X.-D. Wang, A.-D. Yang, G. Celucci, T. E. Graber, C. Dajadian, Y. Yu, C. K. Damgaard, A. J. Berman and T. Alain (2018). "LARP1 is a major phosphorylation substrate of mTORC1." bioRxiv.

Holz, M. K. and J. Blenis (2005). "Identification of S6 kinase 1 as a novel mammalian target of rapamycin (mTOR)-phosphorylating kinase." J Biol Chem **280**(28): 26089-26093.

Ichimura, Y., S. Waguri, Y. S. Sou, S. Kageyama, J. Hasegawa, R. Ishimura, T. Saito, Y. Yang, T. Kouno, T. Fukutomi, T. Hoshii, A. Hirao, K. Takagi, T. Mizushima, H. Motohashi, M. S. Lee, T. Yoshimori, K. Tanaka, M. Yamamoto and M. Komatsu (2013). "Phosphorylation of p62 activates the Keap1-Nrf2 pathway during selective autophagy." Mol Cell **51**(5): 618-631.

Jia, J. J., R. M. Lahr, M. T. Solgaard, B. J. Moraes, R. Pointet, A. D. Yang, G. Celucci, T. E. Graber, H. D. Hoang, M. R. Niklaus, I. A. Pena, A. K. Hollensen, E. M. Smith, M. Chaker-Margot, L. Anton, C. Dajadian, M. Livingstone, J. Hearnden, X. D. Wang, Y. Yu, T. Maier, C. K. Damgaard, A. J. Berman, T. Alain and B. D. Fonseca (2021). "mTORC1 promotes TOP mRNA translation through site-specific phosphorylation of LARP1." Nucleic Acids Res **49**(6): 3461-3489.

Lee, G., Y. Zheng, S. Cho, C. Jang, C. England, J. M. Dempsey, Y. Yu, X. Liu, L. He, P. M. Cavaliere, A. Chavez, E. Zhang, M. Isik, A. Couvillon, N. E. Dephoure, T. K. Blackwell, J. J. Yu, J. D. Rabinowitz, L. C. Cantley and J. Blenis (2017). "Post-transcriptional Regulation of De Novo Lipogenesis by mTORC1-S6K1-SRPK2 Signaling." Cell **171**(7): 1545-1558 e1518.

Piao, X., T. Kobayashi, L. Wang, M. Shiono, Y. Takagi, G. Sun, M. Abe, Y. Hagiwara, D. Zhang, K. Okimoto, M. Kouchi, I. Matsumoto and O. Hino (2009). "Regulation of folliculin (the BHD gene product) phosphorylation by Tsc2-mTOR pathway." Biochem Biophys Res Commun **389**(1): 16-21.

Ramirez-Valle, F., S. Braunstein, J. Zavadil, S. C. Formenti and R. J. Schneider (2008). "eIF4GI links nutrient sensing by mTOR to cell proliferation and inhibition of autophagy." J Cell Biol **181**(2): 293-307.

Raught, B., A. C. Gingras, S. P. Gygi, H. Imataka, S. Morino, A. Gradi, R. Aebersold and N. Sonenberg (2000). "Serum-stimulated, rapamycin-sensitive phosphorylation sites in the eukaryotic translation initiation factor 4GI." Embo j **19**(3): 434-444.
